# A framework for evaluating correspondence between brain images using anatomical fiducials

**DOI:** 10.1101/460675

**Authors:** Jonathan C. Lau, Andrew G. Parrent, John Demarco, Geetika Gupta, Jason Kai, Olivia W. Stanley, Tristan Kuehn, Patrick J. Park, Kayla Ferko, Ali R. Khan, Terry M. Peters

## Abstract

Accurate spatial correspondence between template and subject images is a crucial step in neuroimaging studies and clinical applications like stereotactic neurosurgery. In the absence of a robust quantitative approach, we sought to propose and validate a set of point landmarks, anatomical fiducials (AFIDs), that could be quickly, accurately, and reliably placed on magnetic resonance images of the human brain. Using several publicly available brain templates and individual participant datasets, novice users could be trained to place a set of 32 AFIDs with millimetric accuracy. Furthermore, the utility of the AFIDs protocol is demonstrated for evaluating subject-to-template and template-to-template registration. Specifically, we found that commonly used voxel overlap metrics were relatively insensitive to focal misregistrations compared to AFID point-based measures. Our entire protocol and study framework leverages open resources and tools, and has been developed with full transparency in mind so that others may freely use, adopt, and modify. This protocol holds value for a broad number of applications including alignment of brain images and teaching neuroanatomy.

## Introduction

Establishing spatial correspondence between images is a crucial step in neuroimaging studies enabling fusion of multimodal information, analysis of focal morphological differences, and comparison of within- and between-study data in a common coordinate space. Stereotaxy arose as a result of questions raised by scientists and surgeons interested in the physiology and treatment of focal brain structures (A. C. Evans, Janke, Collins, & Baillet, 2012; Horsley & Clarke, 1908; Peters, 2006). Jean Talairach played a crucial role, observing consistent anatomical features on lateral pneumoencephalograms (Dandy, 1918), or “air studies”, that could be consistently localized, specifically the anterior commissure (AC) and posterior commissure (PC) (Schaltenbrand & Wahren, 1977; J Talairach, David, Tournoux, Corredor, & Kvasina, 1957), and could thus be mapped to prepared post-mortem brain sections in a 3D coordinate system. The AC-PC line has remained important in the era since magnetic resonance imaging (MRI) has risen to prominence for aligning brain images to create population atlases (Collins, Neelin, Peters, & Evans, 1994; A. Evans et al., 1992; Jean Talairach & Tournoux, 1988) as well as to project data from structural and functional investigations. Further optimizations enabled by deformable registration have led to atlas enhancements (Fonov et al., 2011) where many more structural features are preserved. The adoption of standard templates has allowed researchers to compile cytoarchitectonic, functional, and structural data across studies via image-based meta-analysis of peak coordinates and statistical maps (Eickhoff et al., 2009; Gorgolewski et al., 2015; Yarkoni, Poldrack, Nichols, Van Essen, & Wager, 2011).

Ever since the first linearly aligned population templates (A. Evans et al., 1992; Jean Talairach & Tournoux, 1988), there have been a number of advances in the development of robust higher order nonlinear registration tools. As the options became more numerous, several studies investigated the performance of the different nonlinear registration algorithms (Chakravarty et al., 2009; A. C. Evans et al., 2012; Hellier et al., 2003; Klein et al., 2009). Over the past decade, the most common metrics used to evaluate spatial correspondence are related to voxel overlap between regions-of-interest (ROIs) segmented in both reference and target images. Typically, large subcortical structures well-visualized on standard structural MRIs such as the globus pallidus (pallidum), striatum, and thalamus are used (Chakravarty et al., 2009; Chakravarty, Sadikot, Germann, Bertrand, & Collins, 2008; Klein et al., 2009). While these measures are effective for evaluating spatial correspondence on the macroscale, here we argue that they remain relatively coarse measures of registration quality and are insensitive to focal misregistration between images. In addition, they do not permit facile identification or description of where these local biases are occurring. These issues are particularly critical as technical advancements in both imaging and stereotaxy are enabling more accurate therapeutic modulation of brain regions where several millimeters could represent the difference between optimal therapy and complications.

In this paper, we sought inspiration from classical stereotactic methods (Schaltenbrand & Wahren, 1977; J Talairach et al., 1957), and propose that point-based distances provide a more sensitive metric by which brain image correspondence can be evaluated. Anatomical points have been referred to in the literature using a variety of terms including fiducials, landmarks, markups (sometimes used in combination) but ultimately involve representing an anatomical feature by a three-dimensional (x,y,z) Cartesian coordinate. For this manuscript, we have chosen to use the term *AFIDs*, short for *anatomical fiducials*, “fiducia” being Latin for *trust* or *confidence*. We argue that the advent of automatic segmentation-based methods has led to a relative underemphasis of point correspondence between brain structures. We first sought to determine whether we could define a set of AFIDs that were both consistently identifiable across multiple datasets while also providing a distributed sampling about the brain. Following this, we demonstrate how AFIDs are complementary to segmentation-based metrics for providing a quantitative report of spatial correspondence between structural magnetic resonance images of the brain using more intuitive distance-based measures of alignment. Central to this work was the development of our protocol using an open source framework, enabling reproducibility across sites and centers. The overall study organization is shown schematically in Fig 1.

**Fig 1.**
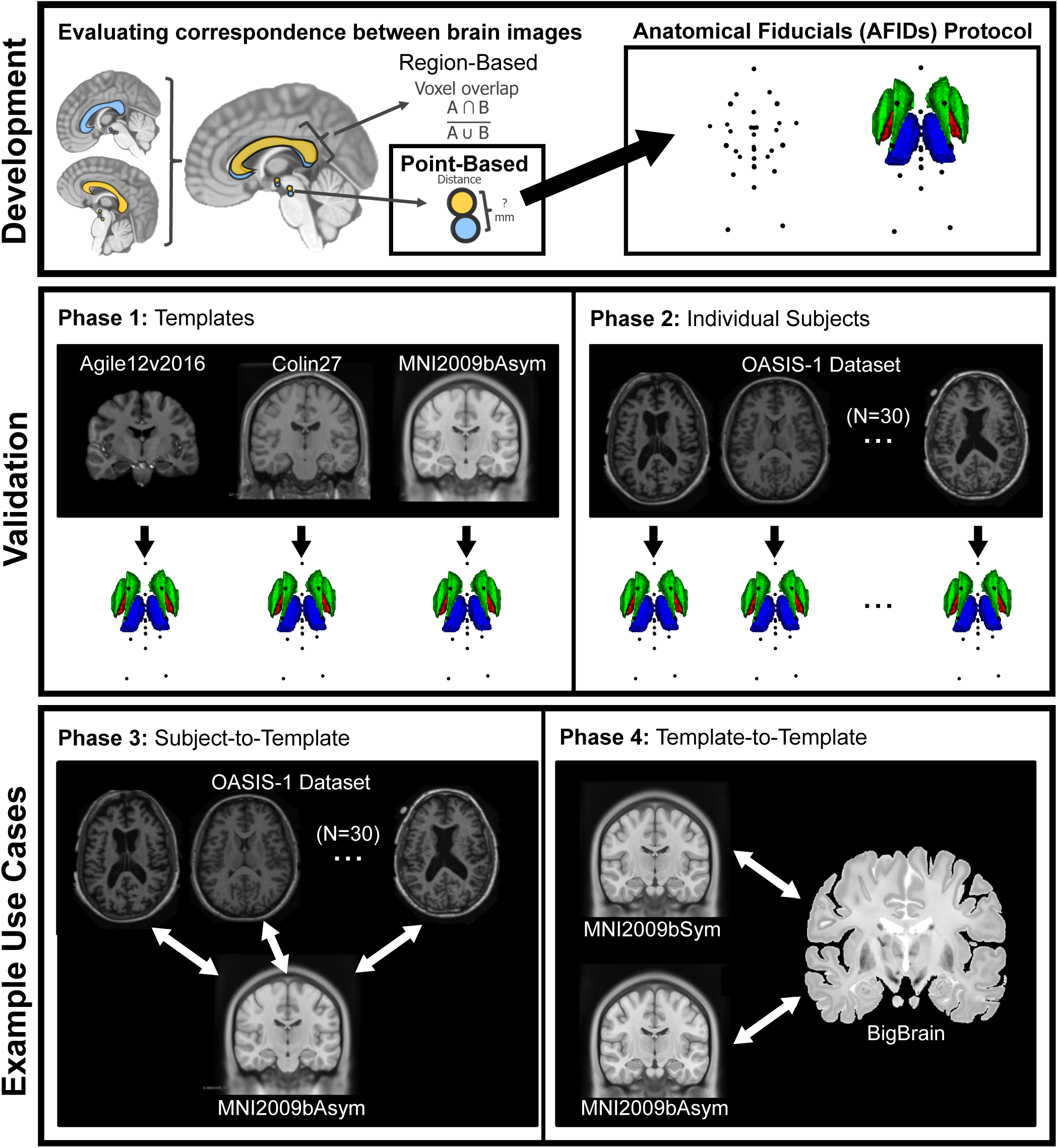
Metrics for evaluating spatial correspondence between brain images include voxel overlap (i.e. ROI-based) metrics as well as point-based distance metrics. The proposed framework involves the identification of point-based anatomical fiducials (AFIDs) in a series of brain images, which provide an intuitive millimetric estimate of correspondence error between images and is also a useful tool for teaching neuroanatomy.

## Methods

### Protocol development

A series of anatomical fiducials (AFIDs) were identified by the lead author (JCL; 10 years experience in neuroanatomy) in consultation with an experienced neurosurgeon (AGP; 20+ years experience practicing stereotactic and functional neurosurgery) with consensus achieved on a set of 32 points (see Fig 2; RRID:SCR_016623). AFIDs could generally be classified as midline (10/32 = 31.25%) or lateral (22/32; i.e. 11 structures that could be placed on each of the left and right sides). Regions prone to geometric distortion were avoided (Lau et al., 2018). We limited our initial set of AFID locations to deep brain regions where less inter-subject variability exists (millimeter scale) compared to the cortical sulci and gyri (centimeter scale) (Thompson, Schwartz, Lin, Khan, & Toga, 1996).

**Fig 2.**
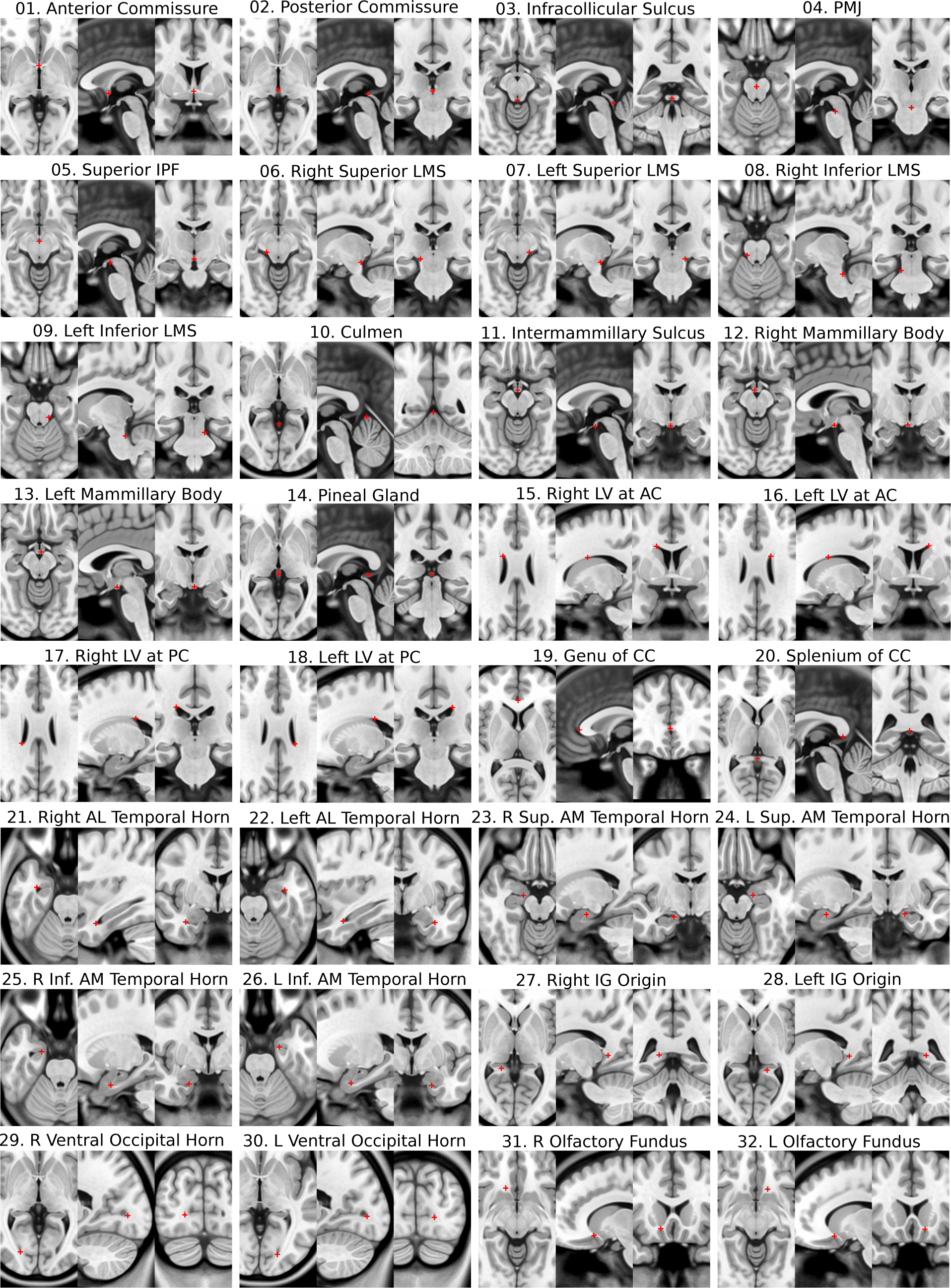
Each of the 32 anatomical fiducials in the protocol is demonstrated with crosshairs at the representative location in MNI2009bAsym space using the standard cardinal planes. AC = anterior commissure; PC = posterior commissure; AL = anterolateral; AM = anteromedial; IG = indusium griseum; IPF = interpeduncular fossa; LMS = lateral mesencephalic sulcus; LV = lateral ventricle; PMJ = pontomesenphalic junction.

The AFID points were placed using the Markups Module of 3D Slicer version 4.6.2 (Fedorov et al., 2012) (RRID:SCR_005619). One key feature of 3D Slicer is that it allows markup points to be placed in the 3D coordinate system of the software as opposed to the voxel coordinate system of the image being annotated permitting more refined (sub-voxel) localization. Images are automatically linearly interpolated by the software on zoom. After importing the structural MRI scan to be annotated into 3D Slicer, the anterior commissure (AC) and posterior commissure (PC) points were placed—specifically at the center of each commissure rather than the intraventricular edge. After defining an additional midline point (typically the pontomesencephalic junction or intermamillary sulcus), an AC-PC transformation was performed using the built-in Slicer module (AC-PC Transform). For all subsequent AFID placements, the AC-PC aligned image was used. The entire protocol is shown in MNI2009bAsym space in Fig 2.

The rest of the methods are organized into four separate phases (see Fig 1). Phase 1 involved AFID placement in three open access brain templates. Phase 2 involved further placement of the AFIDs in individual subject scans. In Phase 3, AFIDs were used to evaluate subject-to-template registration; and finally, in Phase 4, they were used to assess template-to-template registration quality.

For validation and assessment, we adopted the terminology of Fitzpatrick and colleagues (Fitzpatrick & West, 2001; Fitzpatrick, West, & Maurer, 1998) who defined fiducial localization error (FLE) and fiducial registration error (FRE) as metrics used to evaluate the real-world accuracy of image-guidance systems used in neurosurgery. FLE is defined as error related to the placement (i.e. localization) of fiducials, while FRE is defined as error related to registration. This body of work has been most concerned with describing the correspondence between preoperative images of a patient and the physical location of the patient and surgical site in the operating room. Here, we use these terms to describe (virtual, image-based) *anatomical* fiducials (AFIDs) annotated in structural T1-weighted MRI scans.

### Phase 1: Protocol validation for brain templates

Novice participants (N=8) were trained over a series of neuroanatomy tutorials to place AFIDs on a number of publicly available brain images: Agile12v2016 (Lau et al., 2017; Wang et al., 2016), Colin27 (Holmes et al., 1998), MNI2009bAsym (nonlinear asymmetric; version 2009b; RRID:SCR_008796) (Fonov et al., 2011). Each participant then performed 4 rating sessions independently for each template, for a total of 12 point sets resulting in a total of 96 protocols. We computed several different metrics for describing the accuracy (and reliability) of our proposed protocol, all of which are variations of *anatomical fiducial localization error* (AFLE): *mean AFLE, intra-rater AFLE*, and *inter-rater AFLE* as shown in Fig 3.

**Fig 3.**
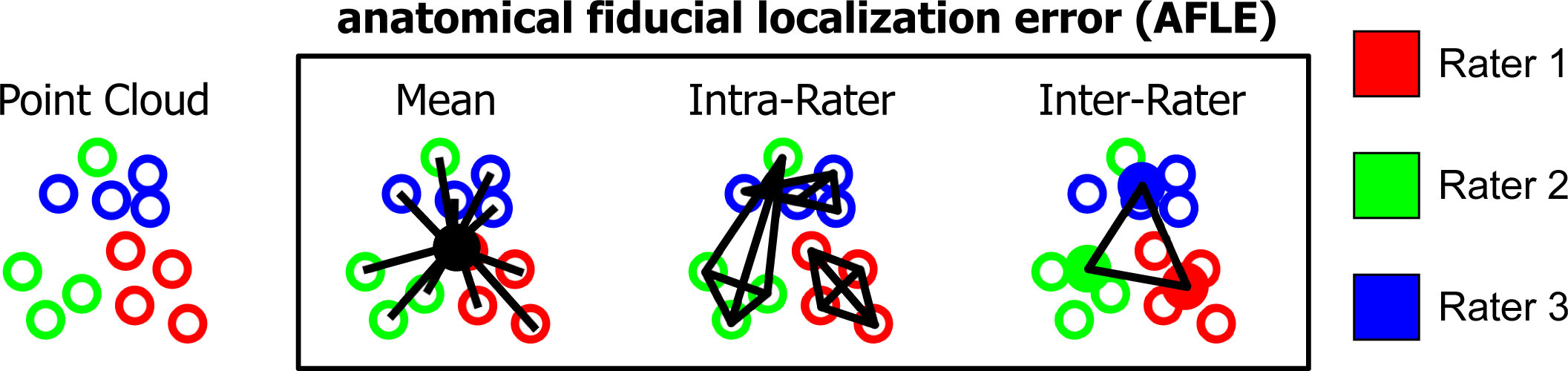
Metrics used for validating AFID placements are shown here in schematic form. Mean, intra-rater, and inter-rater AFLE can be computed for an image that has been rated by multiple raters multiple times.

To compute the *mean AFLE*, the mean AFID coordinate for each brain image was used as an approximation of the ideal coordinate location. Mean AFLE was calculated as the Euclidean distance between the individual position and the group mean. We furthermore calculated *intra-rater AFLE* as the mean pairwise distance between AFIDs placed by the same rater. The individual measures were averaged across all raters as a summary metric. To calculate *inter-rater AFLE*, a mean coordinate was computed by averaging the coordinates for each rater as an estimate of the ideal coordinate location for the rater; the mean pairwise distance between AFIDs placed across raters was then calculated as a summary metric. We summarized global and location-specific mean AFLE according to a number of variables: template (group versus individual), rating session (1-4), rater, and AFID.

Time required to complete placement for a single MRI was documented by each rater. Outliers were defined as any fiducials deviating from the mean fiducial point by greater than 10 mm. Furthermore, patterns of variability in AFID placement were assessed using K-means clustering of fiducial locations (point clouds) relative to the mean fiducial location.

### Phase 2: Protocol validation for individual subjects

The same participants and the lead author (total N=9) performed additional AFID placement on a series of 30 independent brain images from the OASIS-1 database (Marcus, Fotenos, Csernansky, Morris, & Buckner, 2010) (RRID:SCR_007385). Subjects from the OASIS-1 database were selected from the broad range of ages encountered in the database, restricted to cognitively intact (MMSE 30) participants. Although we controlled for normal cognition by MMSE, we selected for qualitatively challenging images with more complex anatomy (asymmetric anatomy and/or variably-sized ventricles). Details on the 30 scans are provided in the S2 file and organized into the Brain Imaging Data Structure (BIDS) format (Gorgolewski, Auer, Calhoun, Craddock, & Das, 2016) (RRID:SCR_016124).

Each of the 9 participants placed 10 independent protocols (90 protocols; 2880 individual points). Each of the 30 MRI scans from the OASIS-1 database had AFIDs placed by 3 raters to establish *inter-rater AFLE* (as described in Methods Section Phase 1: Protocol Validation for Brain Templates). Intra-rater AFLE was not evaluated in Phase 2. Quality of rigid registration was visually inspected by an experienced rater (JL).

#### Region-of-interest segmentation

BIDS formatting permitted automatic processing of each of the included OASIS-1 subjects using fMRIPrep version 1.1.1 (Esteban et al., 2018; Gorgolewski et al., 2017) (RRID:SCR_016216) with anatomical image processing only. Briefly, the fMRIPrep pipeline involves linear and deformable registration to the MNI2009cAsym template (Avants, Epstein, Grossman, & Gee, 2008; Fonov et al., 2011) then processing of the structural MRI through Freesurfer for cortical surface and subcortical volumetric labeling (Dale, Fischl, & Sereno, 1999; Bruce Fischl, 2012) (RRID:SCR_001847). We focused on using ROIs commonly used in the literature to evaluate quality of registration in the subcortex (Chakravarty et al., 2009; Hellier et al., 2003; Klein et al., 2009), i.e. the pallidum, striatum, and thalamus provided as part of the fMRIPrep output run through FreeSurfer. The striatum label required combining the ipsilateral caudate nucleus, accumbens, and putamen labels.

#### Online Validator

In order to better automate the examination of individual fiducial placements by novice raters, an online validator tool was developed (https://github.com/afids). The alpha version is a webpage permitting trainees to upload their own file containing fiducial placements and calculating the Euclidean error for each fiducial they marked relative to a predefined template. These templates are selected from the linked AFIDs repository itself and will be extensible as the project grows. This tool will allow users to compare their results against ground truth results facilitating training.

### Phase 3: Evaluating subject-to-template registration

We evaluated the quality of subject-to-template registration using the output provided as part of fMRIPrep version 1.1.1 using conventional ROI-based metrics (i.e. voxel overlap) as well as distance metrics derived from our manual annotations from Phases 1 and 2. The default template for fMRIPrep 1.1.1 was the MNI2009cAsym template. We started by visually inspecting the images qualitatively from the output fMRIPrep html pages. For each individual subject scan, we used the mean fiducial location as the optimal location calculated in Phase 2. The distance between the individual subject AFID location and the corresponding mean AFID location in the template was computed and defined as the *anatomical fiducial registration error* (AFRE) and computed for linear transformation alone (lin) and combined linear and nonlinear transformation (nlin). Our definition of AFRE differs from the FRE used by Fitzpatrick whose framework for neuronavigation was necessarily limited to rigid-body transformations (Fitzpatrick et al., 1998). This was compared with ROI-based measures of spatial correspondence, specifically, the Jaccard similarity coefficient 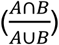 and the Dice kappa coefficient 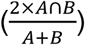, where A and B are the number of voxels in the source and reference images, respectively.

We were able to use the points placed in Phase 1 for the MNI2009bAsym template since the only difference between the MNI2009bAsym and MNI2009cAsym templates was the resampling from 0.5 mm to 1 mm isotropic resolution. AFRE was computed for each AFID location and OASIS-1 subject, along with voxel overlap for the pallidum, striatum, and thalamus. Comparisons between AFRE and voxel overlap were made using Kendall’s tau.

### Phase 4: Evaluating template-to-template registration

BigBrain is a publicly available ultrahigh-resolution (20 micron) human brain model that has enabled bridging of macroscale anatomy with near cellular anatomy (Amunts et al., 2013) (RRID:SCR_001593). A deformable mapping provided by the MNI group has permitted the exploration of high-resolution BigBrain neuroanatomy in MNI2009bSym space (BigBrainRelease.2015; Last modified August 21, 2016; accessed August 2, 2018; Available at: ftp://bigbrain.loris.ca/BigBrainRelease.2015/3D_Volumes/MNI-ICBM152_Space/). In this manuscript, we refer to the registered BigBrain image as BigBrainSym. We quantify the spatial correspondence between BigBrainSym and MNI2009bSym as well as BigBrainSym and MNI2009bAsym templates using the AFIDs protocol to determine whether any significant AFRE could be identified. For MNI2009bAsym, we used mean coordinates for each AFID using rater data from Phase 1. BigBrainSym and MNI2009bSym templates were annotated *de novo* by three experienced raters (GG, JL, KF). The mean AFID coordinate was used as an approximation of the ideal coordinate location for each template. Spatial correspondence was estimated as the AFRE (i.e. Euclidean distance between points) for each AFID. Correlation between AFLE and AFRE were assessed using Kendall’s tau.

### Source code and data availability

All data analysis was performed using R-project version 3.5.1. The AFIDs protocol, raw and processed data, processing scripts, and scripts used in this manuscript are available at: https://github.com/afids. The templates used in this study and salient features of these templates are summarized in Table 1.

**Table 1.** Summary of templates used in this study.

| Template | N | Features | References |
| --- | --- | --- | --- |
| Agile12v2016 | 12 | Combined T1w and T2w template acquired at 7-Tesla. | (Lau et al., 2017; Wang et al., 2016) |
| BigBrainSym | 1 | Ultra-high resolution histological template registered to MNI2009bSym. | (Amunts et al., 2013) |
| Colin27 | 1 | Single subject scanned 27 times at 1.5-Tesla. | (Holmes et al., 1998) |
| MNI2009bAsym | 152 | Population template most commonly used in the literature. | (Fonov et al., 2011) |
| MNI2009bSym | 152 | Symmetric version of MNI2009bAsym. | (Fonov et al., 2011) |
| OASIS-1 | 1 | Dataset of publicly available T1w anatomical MRI scans. | (Marcus et al., 2010) |

N = number of subjects for each template (Note: some are single subject rather than population templates).

## Results

### Phase 1: Protocol validation for brain templates

The 8 raters had a mean experience of 11.5 +/- 11.2 months in medical imaging (range: 0-24 months), 14.3 +/- 17.0 months in neuroanatomy (range: 0-48 months), and 7.0 +/- 8.8 months in 3D Slicer (range: 0-24 months). During the template validation phase, the raters placed a total of 3072 individual points (number of sessions = 4; templates = 3; points = 32). Average placement time for a single brain image was estimated at between 20-40 minutes. Thus, a total of 1920-3840 minutes (or 32-64 hours) were logged in this phase of the study. The mean, intra-rater, and inter-rater AFLE metrics are summarized in Table 2.

**Table 2.** Summary of fiducial localization error across brain templates.

| Template | Before QC |  | After QC |  |  |  |
| --- | --- | --- | --- | --- | --- | --- |
|  | mean AFLE (mm) | # of outliers (%) | mean AFLE (mm) | # of outliers (%) | intra-rater AFLE (mm) | inter-rater AFLE (mm) |
| **Agile12v2016** | 1.10 +/- 1.59 | 3/1024 (0.29%) | 1.01 +/- 0.93 | 0/1021 (0.00%) | 1.13 +/- 0.86 | 1.14 +/- 0.48 |
| **Colin27** | 1.71 +/- 2.78 | 20/1024 (1.95%) | 1.11 +/- 1.05 | 1/1004 (0.10%) | 1.14 +/- 0.92 | 1.36 +/- 0.88 |
| **MNI2009bAsym** | 0.99 +/- 1.11 | 1/1024 (0.10%) | 0.97 +/- 0.80 | 0/1023 (0.00%) | 1.03 +/- 0.78 | 1.07 +/- 0.46 |
| **Total** | **1.27 +/- 1.98** | **24/3072 (0.78%)** | **1.03 +/- 0.94** | **1/3048 (0.03%)** | **1.10 +/- 0.86** | **1.19 +/- 0.64** |

For the raw data, the mean AFLE was 1.27 +/- 1.98 mm (1.10 +/- 1.59 mm for Agile12v2016; 1.71 +/- 2.78 mm for Colin27; 0.99 +/- 1.11 mm for MNI2009bAsym). Using a threshold of mean AFLE greater than 10 mm from the group mean, we identified 24 outliers out of 3072 independent points (0.78%). 20/24 (83.33%) of outliers were the result of variable placement in the bilateral ventral occipital horns (i.e. AFID29 and AFID30) of the Colin27 template. One pair (2/24; 8.33%) of outliers was due to left-right mislabeling (indusium griseum; AFID27 and AFID28). One additional point was mislabeled; i.e. the left anterolateral temporal horn point (AFID22) was placed at the left inferior anteromedial horn location (AFID26). After quality control (QC) and filtering outliers, mean AFLE improved to 1.03 +/- 0.94 mm (1.01 +/- 0.93 mm for Agile12v2016; 1.11 +/- 1.05 mm for Colin27; 0.97 +/- 0.80 mm for MNI2009bAsym).

Intra-rater AFLE was 1.10 +/- 0.86 mm (1.13 +/- 0.86 mm for Agile12v2016; 1.14 +/- 0.92 mm for Colin27; 1.03 +/- 0.78 mm); and inter-rater AFLE was 1.19 +/- 0.65 mm (1.15 +/- 0.49 mm for Agile12v2016; 1.36 +/- 0.88 mm for Colin27; 1.07 +/- 0.46 mm for MNI2009bAsym). Mean, intra-rater, and inter-rater AFLE for each AFID post-QC are summarized in the Supporting Information S1 File.

All subsequent analyses were performed using the mean AFLE metric. We performed a one-way analysis of variance observing evidence of statistically different variance between templates (F-value = 7.88; p-value < 0.001). Differences in mean AFLE between templates were identified on subgroup analysis for the right superior lateral mesencephalic sulcus (AFID06), culmen (AFID10), genu of the corpus callosum (AFID19), and left superior anteromedial temporal horn (AFID24), suggesting differences between templates that may contribute to errors in placement. The results for each AFID are also summarized in the Supporting Information S1 File.

Furthermore, we observed several distinct patterns of AFID placement using K-means clustering of fiducial locations (point clouds) relative to the mean fiducial location (see Fig 4). We identified three different general patterns of point cloud distributions ranging from highly anisotropic to moderately anisotropic to isotropic.

**Fig 4.**
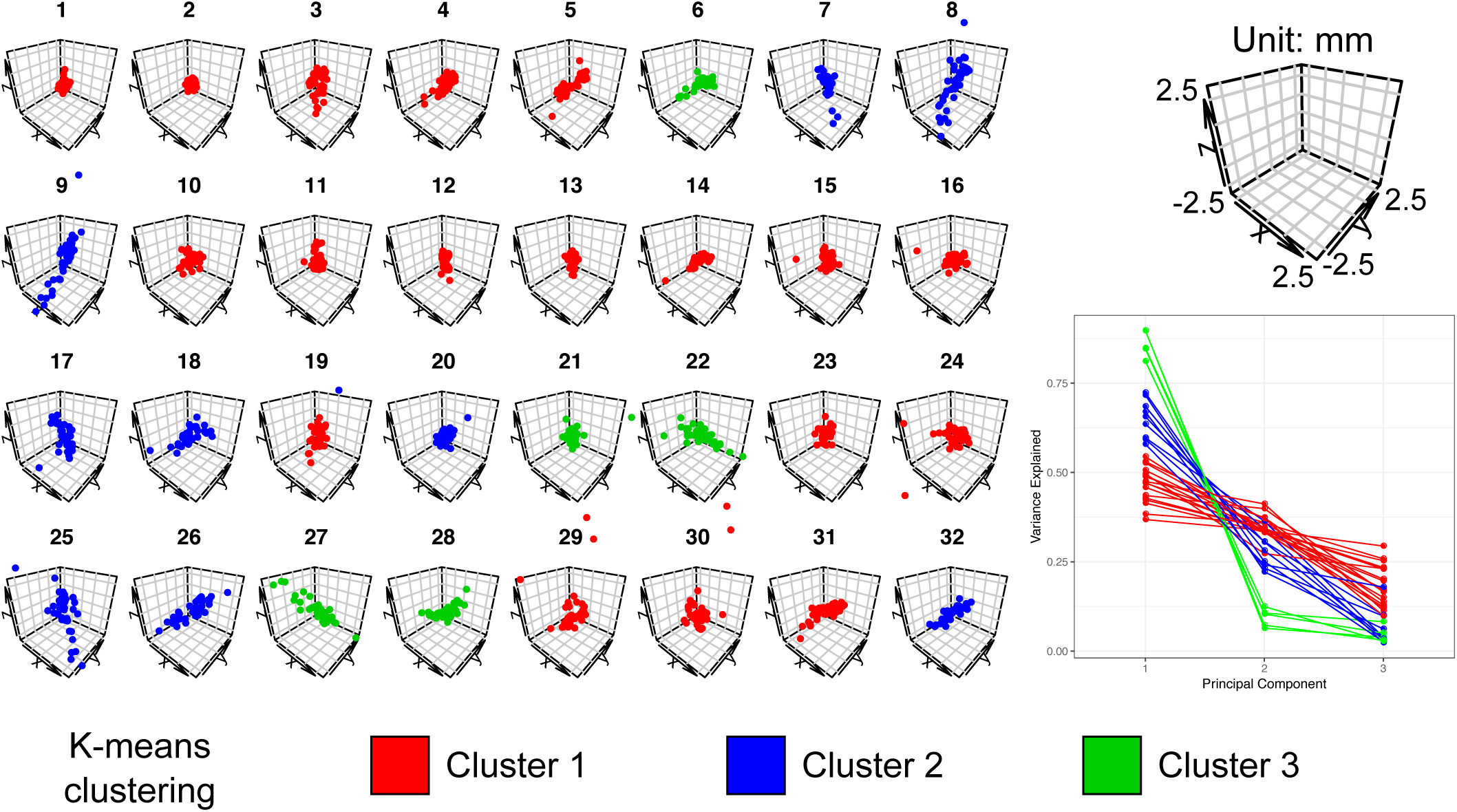
K-means clustering of point clouds relative to the mean fiducial location for each of the 32 AFIDs (left). Principle components analysis (bottom right) revealed three different general patterns were identified ranging from highly isotropic (Cluster 1: red) to moderately anisotropic (Cluster 2: blue) to anisotropic (Cluster 3: green). Results are shown for the MNI2009bAsym template. See the Supplementary Materials for similar plots for Agile12v2016, Colin27, and the templates combined.

### Phase 2: Protocol validation for individual subjects

During the individual subject validation phase, 9 participants completed 10 AFID protocols (= 90 total protocols) and a total of 2880 individual points distributed equally among 30 OASIS-1 datasets. We identified 28 outliers (0.97%), defined as individual point placements greater than 1 cm (10 mm) away from the group mean. 8/28 outliers (28.57%) were the result of mislabeled points: three pairs of lateral (non-midline) AFIDs and only one pair due to gross mislabeling of the target AFID structure (placement in bilateral frontal ventricular horns rather than occipital horns). Beyond left-right swapping, the AFIDs most susceptible to outliers were the following points: bilateral ventral occipital horns (AFID29-30) and bilateral indusium griseum origins (AFID27-28). Mean AFLE across the 30 scans and points was 1.28 +/- 3.03 mm improving to 0.94 +/- 0.73 after filtering out the outliers. Inter-rater AFLE was 1.58 +/- 1.02 mm across all AFIDs. Mean AFLE and inter-rater AFLE are summarized for each AFID in Table 3 and subject in the Supporting Information S2 file.

**Table 3.** Mean and inter-rater fiducial localization error pre- and post-QC for the included OASIS-1 subjects for all AFIDs.

| AFID | Description | Pre-QC | Post-QC |  |
| --- | --- | --- | --- | --- |
| | | Mean AFLE<br>mean ± sd (max) | Mean AFLE<br>mean ± sd (max) | Inter-Rater AFLE<br>mean ± sd (max) |
| 01 | AC | 0.36±0.21 (1.29) | 0.36±0.21 (1.29) | 0.60±0.25 (1.38) |
| 02 | PC | 0.34±0.16 (0.88) | 0.34±0.16 (0.88) | 0.57±0.21 (1.22) |
| 03 | infracollicular sulcus | 0.78±0.48 (3.07) | 0.78±0.48 (3.07) | 1.34±0.64 (3.84) |
| 04 | PMJ | 0.83±0.49 (2.44) | 0.83±0.49 (2.44) | 1.41±0.55 (2.55) |
| 05 | superior interpeduncular fossa | 1.20±0.75 (3.50) | 1.20±0.75 (3.50) | 2.04±0.90 (4.25) |
| 06 | R superior LMS | 1.30±1.74 (14.25) | 1.01±0.55 (2.85) | 1.70±0.68 (3.13) |
| 07 | L superior LMS | 1.36±1.71 (13.99) | 1.06±0.61 (3.45) | 1.72±0.71 (3.89) |
| 08 | R inferior LMS | 1.13±0.75 (5.13) | 1.03±0.57 (2.99) | 1.77±0.74 (3.43) |
| 09 | L inferior LMS | 1.10±0.80 (5.31) | 1.01±0.62 (2.72) | 1.71±0.86 (3.71) |
| 10 | culmen | 0.99±0.99 (5.66) | 0.83±0.62 (3.07) | 1.35±0.82 (3.42) |
| 11 | intermamillary sulcus | 0.60±0.31 (1.62) | 0.60±0.31 (1.62) | 1.02±0.41 (1.86) |
| 12 | R MB | 0.40±0.23 (1.11) | 0.40±0.23 (1.11) | 0.69±0.32 (1.52) |
| 13 | L MB | 0.36±0.20 (1.20) | 0.36±0.20 (1.20) | 0.62±0.29 (1.62) |
| 14 | pineal gland | 0.68±0.47 (1.98) | 0.68±0.47 (1.98) | 1.16±0.69 (2.63) |
| 15 | R LV at AC | 1.00±0.90 (5.28) | 0.91±0.72 (4.45) | 1.55±1.08 (5.86) |
| 16 | L LV at AC | 1.01±0.80 (4.53) | 0.94±0.70 (4.53) | 1.60±1.08 (5.47) |
| 17 | R LV at PC | 0.92±0.54 (3.42) | 0.92±0.54 (3.42) | 1.54±0.77 (3.84) |
| 18 | L LV at PC | 0.87±0.42 (2.20) | 0.87±0.42 (2.20) | 1.46±0.55 (2.80) |
| 19 | genu of CC | 0.97±0.81 (5.16) | 0.89±0.63 (3.69) | 1.50±0.89 (4.30) |
| 20 | splenium | 0.54±0.25 (1.24) | 0.54±0.25 (1.24) | 0.91±0.35 (1.66) |
| 21 | R AL temporal horn | 1.44±1.09 (7.01) | 1.30±0.86 (4.45) | 2.21±1.13 (5.92) |
| 22 | L AL temporal horn | 1.22±0.77 (4.11) | 1.22±0.77 (4.11) | 2.04±1.01 (4.47) |
| 23 | R superior AM temporal horn | 1.28±1.27 (8.22) | 1.12±0.88 (4.69) | 1.86±1.19 (4.97) |
| 24 | L superior AM temporal horn | 1.09±1.22 (7.54) | 0.83±0.61 (3.66) | 1.39±0.85 (4.60) |
| 25 | R inferior AM temporal horn | 1.69±1.43 (9.03) | 1.44±0.91 (4.72) | 2.39±1.23 (5.07) |
| 26 | L inferior AM temporal horn | 1.99±1.75 (8.79) | 1.49±1.09 (4.70) | 2.42±1.47 (6.64) |
| 27 | R indusium griseum origin | 3.13±4.19 (23.44) | 1.77±0.99 (4.77) | 2.95±1.20 (5.75) |
| 28 | L indusium griseum origin | 2.99±4.30 (24.30) | 1.68±1.00 (5.00) | 2.75±1.29 (5.78) |
| 29 | R ventral occipital horn | 3.64±10.36 (78.74) | 0.69±0.39 (2.11) | 1.14±0.54 (2.53) |
| 30 | L ventral occipital horn | 3.43±10.38 (80.42) | 0.86±0.67 (4.94) | 1.39±0.98 (5.72) |
| 31 | R olfactory sulcal fundus | 0.99±0.53 (2.29) | 0.99±0.53 (2.29) | 1.71±0.60 (2.84) |
| 32 | L olfactory sulcal fundus | 1.21±0.74 (4.53) | 1.21±0.74 (4.53) | 2.11±0.92 (5.81) |

### Phase 3: Evaluating subject-to-template registration

The following section uses the AFIDs to evaluate the quality of spatial correspondence between the Phase 2 subject data with the MNI2009cAsym template as processed through fMRIPrep. FMRIPrep ran successfully on 30/30 datasets (100%). Visual inspection of the fMRIPrep generated reports revealed no gross misregistrations between MNI2009c and the individual subject scans although a pattern of worse deformable registration in subjects with enlarged ventricles was observed. The rest of this section is concerned with examining the comparative utility of conventional voxel overlap (ROI-based) metrics against the point-based (AFRE) metric proposed in this study (see Fig 5A).

**Fig 5.**
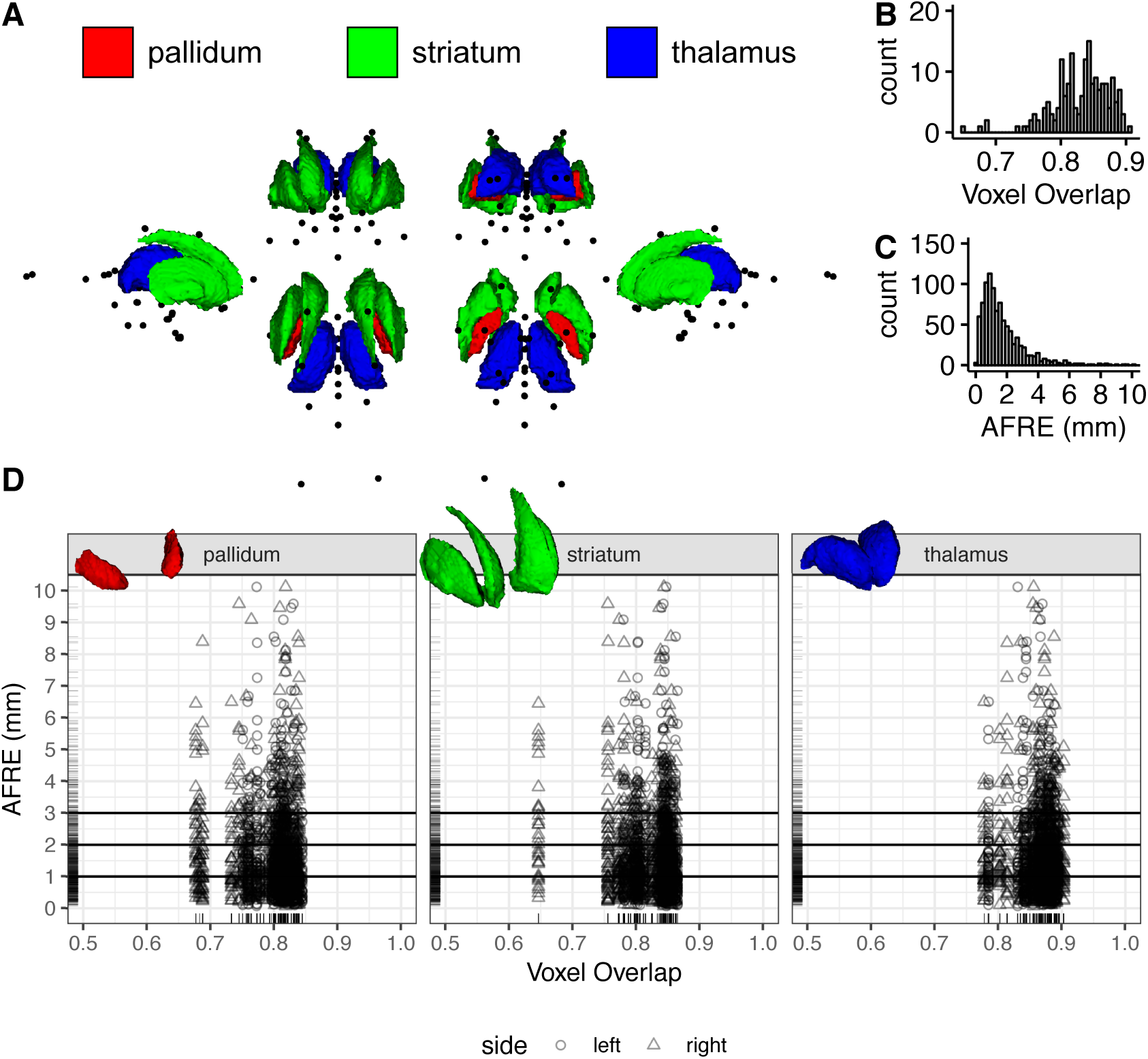
A comparison of voxel overlap and distance metrics for establishing spatial correspondence between brain regions as evaluated on fMRIPrep output. (A) Multiple views showing the location of AFIDs (black dots) relative to three commonly used ROIs used in voxel overlap measures (the pallidum, striatum, and thalamus). (B,C) The histograms for voxel overlap (Jaccard index) and AFRE, respectively. The distribution for AFRE is more unimodal with a more interpretable dynamic range (in mm) compared to voxel overlap. Trellis plots demonstrate evidence of focal misregistrations identified by AFRE not apparent when looking at ROI-based voxel overlap alone (D).

Improvements in overlap were identified when going from linear to combined linear/nonlinear transformations (Table 4). Some heterogeneity in values was noted between ROIs with voxel overlap measures observed to be lowest for the pallidum (the smallest structure evaluated). All Jaccard values after nonlinear transformation were greater than 0.7 (greater than 0.8 for Dice kappa), generally considered to represent good correspondence between two registered images. For simplicity, we report the Jaccard coefficient as our measure of voxel overlap for all subsequent analyses.

**Table 4.** Voxel overlap (Jaccard and Kappa) of the pallidum, striatum, and thalamus after linear registration only and combined linear/nonlinear registration.

| roi | side | Jaccard |  |  | Kappa |  |  |
| --- | --- | --- | --- | --- | --- | --- | --- |
|  |  | lin | nlin |  | lin | nlin |  |
| pallidum | left | 0.54±0.13 | 0.80±0.03 | * | 0.69±0.11 | 0.89±0.02 | * |
|  | right | 0.55±0.12 | 0.79±0.05 | * | 0.70±0.11 | 0.88±0.03 | * |
| striatum | left | 0.53±0.14 | 0.83±0.03 | * | 0.68±0.13 | 0.91±0.02 | * |
|  | right | 0.55±0.15 | 0.82±0.05 | * | 0.70±0.13 | 0.90±0.03 | * |
| thalamus | left | 0.70±0.11 | 0.86±0.03 | * | 0.82±0.08 | 0.93±0.02 | * |
|  | right | 0.69±0.11 | 0.87±0.03 | * | 0.81±0.08 | 0.93±0.02 | * |

\* significant after FDR corrected (q-value < 0.05)

Mean AFRE improved from 3.40 +/- 2.55 mm with linear transformation alone to 1.80 +/- 2.09 with combined linear/nonlinear transformation (p-value < 0.001). AFRE was significantly decreased with nonlinear registration for all AFIDs except the pineal gland (AFID14). AFRE was observed to be higher than mean AFLE measures (see Phase 2: 0.93 +/- 0.73 mm) across the same subjects providing evidence that registration error is detectable beyond the limits of localization error. The number of outlier AFIDs with AFRE > 3 mm (more than 2 standard deviations above the mean AFLE found in Phase 2 for the same subjects) was 135/960 (14.06%), representing 22/32 (68.75%) unique AFIDs identified as misregistered. Each independent OASIS-1 subject had at least one AFID with AFRE > 3 mm with a mean maximum AFRE of 7.5 mm (Range: 3.16-32.78 mm). Although AFLE and AFRE were statistically correlated, the effect size was small (Kendall tau = 0.15; p-value < 0.001; Supporting Information S3 file).

Subgroup analysis for each AFID is summarized in Table 5. AC and PC had the lowest mean AFRE at 0.36 +/- 0.21 and 0.57 +/- 0.29 mm, respectively. However, registration errors as high as 1.64 mm were observed for PC. The ventricles appeared particularly difficult to align on subgroup analysis of the AFIDs. The highest AFRE among all 32 AFIDs was observed for the right and left ventral occipital horns (AFID29-30) at 3.44 +/- 5.77 and 4.51 +/- 6.28 mm respectively with errors in certain cases over 20 mm (OAS1_0109 and OAS1_0203; Supporting Information S3 file). Similarly, the lateral ventricle features (AFID15-18) also demonstrated high AFRE ranging from 2.11-3.01 mm on average and up to 7 mm or more. Finally, the alignment of the temporal horn features (AFID21-26) also support this observation with mean errors of 1.67-2.41 mm with observed errors over 5 mm.

**Table 5.** AFRE after linear registration alone and combined linear/nonlinear registration.

| AFID | Description | Mean AFRE<br>mean ± sd (max) | | |
| --- | --- | --- | --- | --- |
|  |  | lin | nlin |  |
| 01 | AC | 2.15±0.97 (4.96) | 0.36±0.21 (0.99) | * |
| 02 | PC | 1.83±0.96 (4.58) | 0.57±0.29 (1.64) | * |
| 03 | infracollicular sulcus | 2.20±1.23 (5.71) | 0.93±0.53 (2.11) | * |
| 04 | PMJ | 2.50±1.36 (6.06) | 0.68±0.43 (2.13) | * |
| 05 | superior interpeduncular fossa | 2.35±1.06 (4.75) | 0.76±0.37 (1.69) | * |
| 06 | R superior LMS | 2.07±0.95 (4.32) | 1.17±0.74 (3.52) | * |
| 07 | L superior LMS | 2.03±0.85 (4.22) | 1.43±0.77 (2.88) | * |
| 08 | R inferior LMS | 2.45±1.37 (7.50) | 1.78±1.11 (5.41) | * |
| 09 | L inferior LMS | 2.54±1.26 (6.63) | 1.83±0.96 (3.99) | * |
| 10 | culmen | 4.50±2.93 (12.72) | 2.73±2.81 (10.12) | * |
| 11 | intermamillary sulcus | 2.81±1.62 (6.30) | 1.44±0.60 (2.73) | * |
| 12 | R MB | 2.72±1.67 (6.90) | 0.93±0.48 (1.90) | * |
| 13 | L MB | 2.84±1.70 (6.14) | 1.01±0.62 (2.93) | * |
| 14 | pineal gland | 2.53±1.39 (5.70) | 2.01±1.24 (6.16) | |
| 15 | R LV at AC | 4.44±1.84 (7.90) | 2.70±1.59 (7.85) | * |
| 16 | L LV at AC | 4.50±1.95 (8.40) | 2.11±1.72 (7.92) | * |
| 17 | R LV at PC | 4.81±2.54 (10.07) | 2.96±2.42 (9.46) | * |
| 18 | L LV at PC | 4.80±2.64 (10.34) | 3.01±2.22 (8.13) | * |
| 19 | genu of CC | 3.73±1.82 (7.88) | 1.56±0.76 (3.32) | * |
| 20 | splenium | 2.96±1.88 (7.57) | 0.97±0.60 (2.93) | * |
| 21 | R AL temporal horn | 3.79±1.71 (7.50) | 1.70±1.09 (5.23) | * |
| 22 | L AL temporal horn | 3.62±1.45 (6.98) | 1.67±0.98 (4.31) | * |
| 23 | R superior AM temporal horn | 3.34±1.63 (7.25) | 1.93±1.34 (6.85) | * |
| 24 | L superior AM temporal horn | 3.44±1.80 (8.20) | 1.67±1.25 (5.80) | * |
| 25 | R inferior AM temporal horn | 4.02±1.97 (8.32) | 2.41±1.16 (5.61) | * |
| 26 | L inferior AM temporal horn | 4.13±1.70 (8.20) | 2.21±1.09 (4.84) | * |
| 27 | R indusium griseum origin | 3.36±2.07 (8.46) | 2.06±1.49 (6.40) | * |
| 28 | L indusium griseum origin | 3.60±1.68 (8.83) | 2.05±1.37 (5.00) | * |
| 29 | R ventral occipital horn | 5.86±6.32 (36.26) | 3.44±5.77 (32.78) | * |
| 30 | L ventral occipital horn | 6.99±6.72 (33.74) | 4.51±6.28 (29.76) | * |
| 31 | R olfactory sulcal fundus | 2.83±1.36 (7.50) | 1.37±0.95 (3.44) | * |
| 32 | L olfactory sulcal fundus | 2.94±1.28 (6.49) | 1.57±0.84 (3.41) | * |

\* significant after FDR corrected (q-value < 0.05)

AFRE was negatively correlated with voxel overlap but the estimates were small (tau = −0.02; p-value = 0.03). Subgroup analysis demonstrated the same negative trends for the right pallidum and striatum but these results did not survive multiple comparisons correction (Fig 5D). No correlation between voxel overlap measures and individual AFID AFREs survived multiple comparisons correction. Comparing histograms, AFRE demonstrated a more unimodal distribution peaking between 1-2 mm (Fig 5B) while voxel overlap exhibited two peaks within the 0.8-0.9 range (Fig 5C). The AFRE plot also demonstrated a longer tail up to 10 mm, thus permitting a broader dynamic range in which to judge the quality of registration. In contrast, voxel overlap metrics were sparse in the lower range making interpretation more difficult. Finally, we observed that even where voxel overlap was high, suggesting good spatial correspondence, high AFRE values were also observed for certain AFIDs (see Fig 5D). These represent focal AFID locations where two images are misregistered despite stable voxel overlap results (Fig 6).

**Fig 6.**
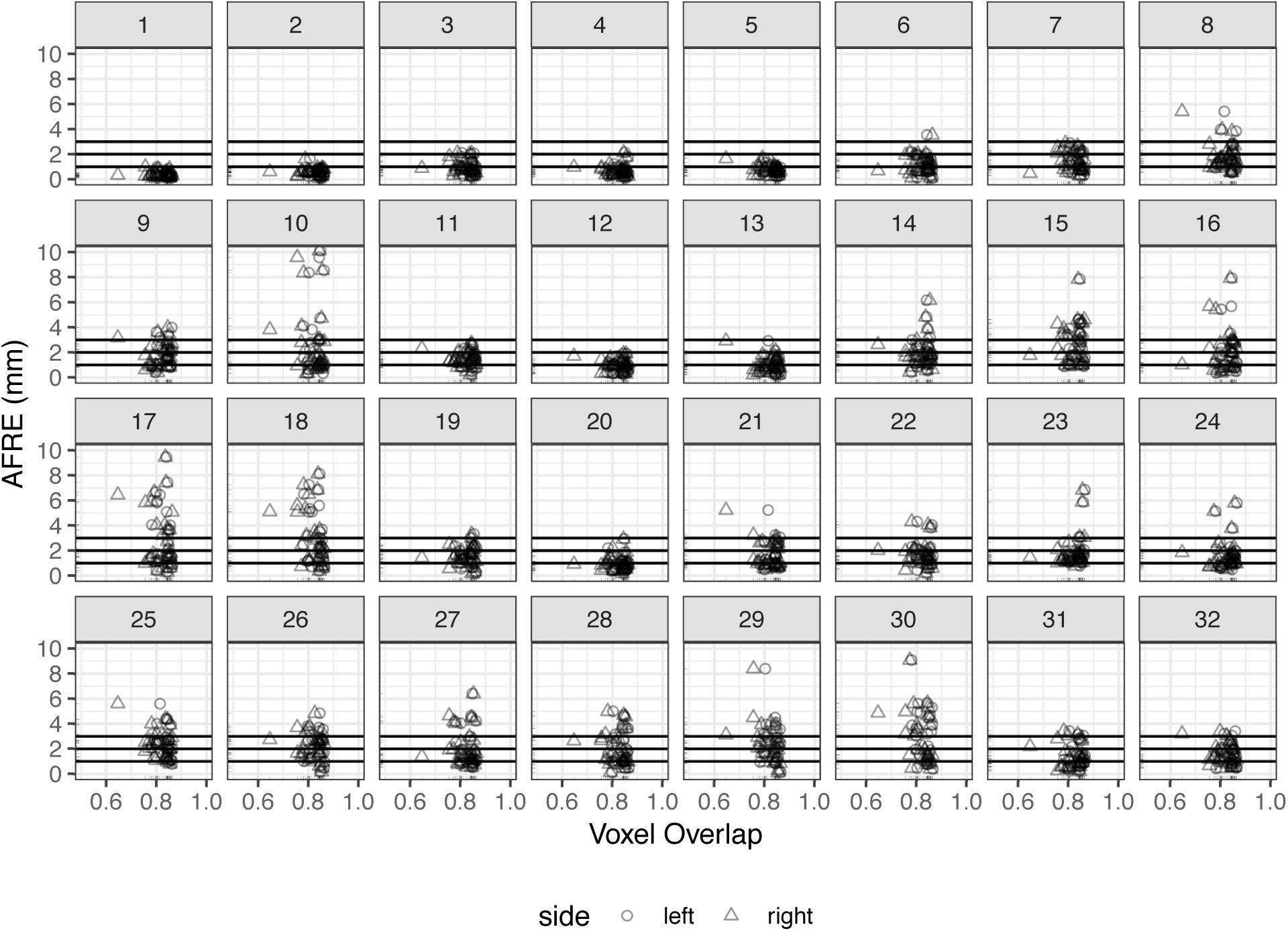
Investigating relationships between voxel overlap of the striatum and AFRE for each AFID. Focal misregistrations are identified using AFRE for the following AFIDs: 8-10, 14-18, 21-30. The most commonly misregistered regions include the inferior mesencephalon, superior vermis, pineal gland, indusium griseum, and ventricular regions. Horizontal lines are used to demarcate tiers of AFLE error above which AFRE values are beyond a threshold of localization error alone, i.e. the top horizontal line at 3 mm represents more than 2 standard deviations beyond the mean AFLE. Separate plots for the pallidum and thalamus ROIs are provided in the Supporting Information S3 file.

### Phase 4: Evaluating template-to-template registration

Mean AFLE for BigBrainSym and MNI2009bSym was 0.59 +/- 0.40 mm combined with no outliers (BigBrainSym: 0.63 +/- 0.50 mm; MNI2009bSym: 0.55 +/- 0.26 mm). We highlighted AFRE values beyond a threshold of 2 mm given this represents more than 2 standard deviations beyond the mean AFLE in the templates being studied. AFRE values beyond this minimum were flagged as highlighting focal misregistrations between templates.

The mean AFRE between BigBrainSym and MNI2009bSym was 2.16 +/- 1.99 mm and between BigBrainSym and MNI2009bAsym was 2.30 +/- 1.83 mm, both above threshold. The largest error was 9.27 mm (MNI2009bSym) and 9.38 mm (MNI2009bAsym), found at the culmen (AFID10). Out of the 32 AFIDs defined, 11 (34.4%) were above threshold for the symmetric template and 12 (37.5%) for the asymmetric template. The most prominent misregistrations tended to occur in the posterior brainstem with the infracollicular sulcus (AFID03) and pineal gland (AFID14) quantified as 6.36 mm and 4.42 mm AFRE, respectively. These registration errors can be seen in Fig 7 and are summarized by AFID in Table 6. In addition, AFRE up to 2.78 mm were observed for AFIDs placed along the lateral mesencephalic sulcus (AFID06-09) and at the superior interpeduncular fossa (AFID05), which represent features demarcating the lateral and superior bounds of midbrain registration. Registration differences between these templates was also above threshold for the left lateral ventricle at the anterior commissure (AFID16), splenium (AFID20), left anterolateral temporal horn (AFID22), bilateral ventral occipital horns (AFID29-30), and bilateral olfactory sulcal fundi (AFID31-32). No correlation between AFRE and AFLE was found using BigBrainSym AFLE (tau = 0.071; p-value = 0.57) or MNI2009bSym AFLE (tau = −0.046; p-value = 0.71). Interestingly, AFRE was somewhat lower with MNI2009bAsym in many midline AFIDs but higher for certain lateral landmarks, i.e. the left inferior anteromedial temporal horn and bilateral origin of the indusium griseum (AFID26-28).

**Table 6.** AFIDs demonstrating evidence of template-to-template misregistration for BigBrainSym with MNI2009bSym and BigBrainSym with MNI2009bAsym as well as correspondence differences between MNI2009bAsym and MNI2009bSym.

| AFID | Description | AFRE (mm) |  |  |  | Distance** (mm) |  |
| --- | --- | --- | --- | --- | --- | --- | --- |
|  |  | BigBrainSym vs MNI2009bSym |  | BigBrainSym vs MNI2009bAsym |  | MNI2009bAsym vs MNI2009bSym |  |
| 03 | infracollicular sulcus | 6.36 | * | 5.48 | * | 0.98 |  |
| 09 | L inferior LMS | 2.78 | * | 2.48 | * | 0.68 |  |
| 10 | culmen | 9.27 | * | 9.39 | * | 0.21 |  |
| 14 | pineal gland | 4.42 | * | 4.16 | * | 0.41 |  |
| 16 | L LV at AC | 2.05 | * | 1.22 |  | 0.86 |  |
| 20 | splenium | 2.23 | * | 2.20 | * | 0.10 |  |
| 22 | L AL temporal horn | 4.69 | * | 3.44 | * | 2.45 | * |
| 26 | L inferior AM temporal horn | 1.88 |  | 2.58 | * | 0.98 |  |
| 27 | R indusium griseum origin | 1.21 |  | 3.60 | * | 2.81 | * |
| 28 | L indusium griseum origin | 0.74 |  | 2.88 | * | 2.29 | * |
| 29 | R ventral occipital horn | 2.54 | * | 3.99 | * | 1.63 |  |
| 30 | L ventral occipital horn | 5.88 | * | 4.22 | * | 2.00 | * |
| 31 | R olfactory sulcal fundus | 2.62 | * | 1.84 |  | 1.10 |  |
| 32 | L olfactory sulcal fundus | 3.06 | * | 4.21 | * | 1.24 |  |

\* AFRE > 2 mm
\*\* Distance between fiducials (not truly a registration error since templates are designed to be in different spaces)

**Fig 7.**
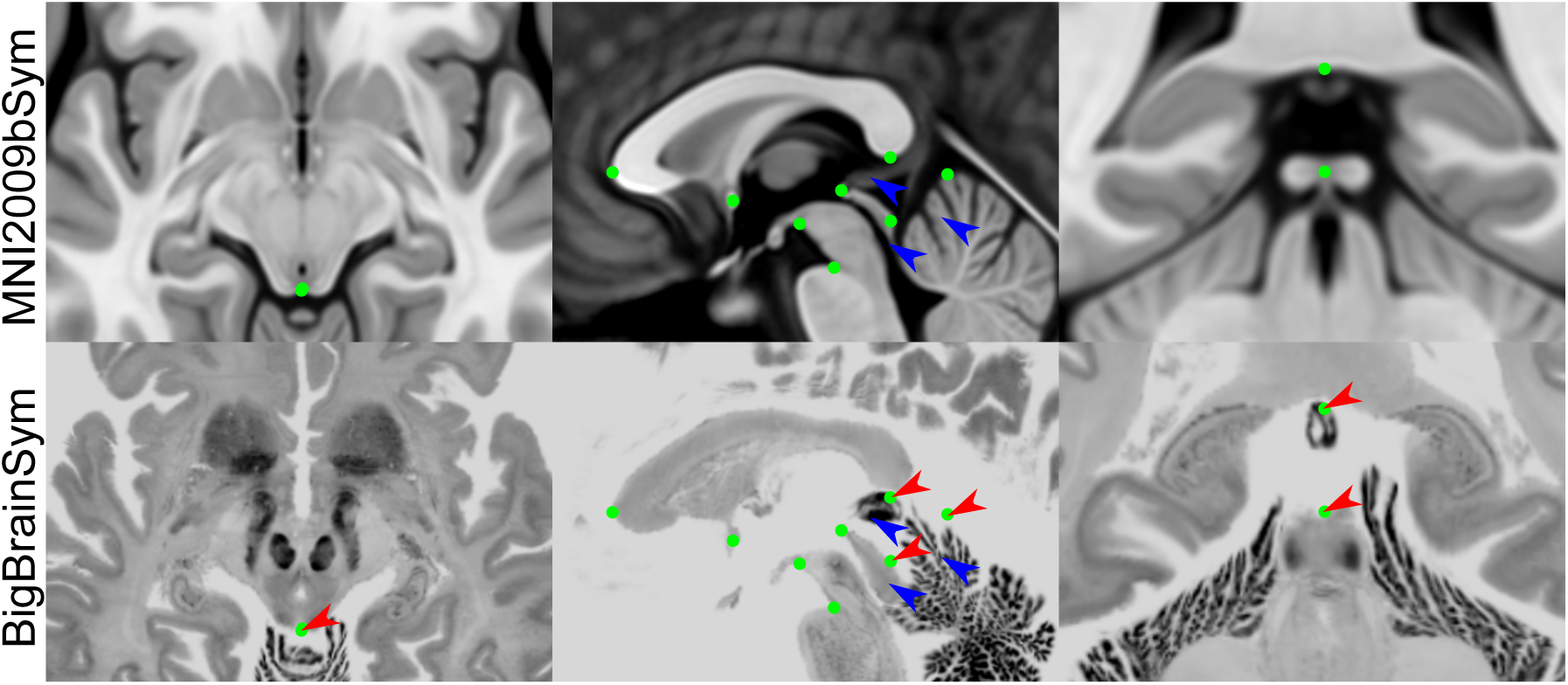
Select views demonstrating registration errors between BigBrainSym and MNI2009bSym. The green dots represent the optimal AFID coordinates in MNI2009bSym space superimposed in both templates to provide a basis for comparing registration differences. While many of the midline AFIDs are stable across both templates, the infracollicular sulcus, pineal gland, splenium, and culmen are misregistered in BigBrainSym (red arrows). The AFIDs draw attention to registration differences in the BigBrainSym space in the tectal plate, pineal gland, and superior vermis (blue arrows).

Finally, we explored the differences in correspondence between the MNI2009bSym and MNI2009bAsym. Note that these differences are not registration errors per se, as the two are not meant to be in the exact same coordinate space. The differences were generally more subtle (0.88 +/- 0.68 mm) but 4 AFIDs (12.5%) were found to be above threshold. As expected, correspondence differences greater than 2 mm occurred in lateral rather than midline AFIDs, specifically at the left anterolateral temporal horn (AFID22), bilateral origins of the indusium griseum (AFID27-28), and left lateral ventral occipital horn (AFID30). No correlations between correspondence and AFLE were found (tau = 0.210; p-value = 0.09).

## Discussion

The present findings demonstrate that a series of anatomical fiducials, referred to here as AFIDs, can be consistently placed on standard structural MR images and can be used to quantify the degree of spatial alignment between brain images in millimeters. We found that AFIDs are reproducible, not overtly manually intensive (20-40 minutes once trained), and more sensitive to local registration errors than standard voxel overlap measures. Our entire protocol and study framework leverages open resources and tools, and has been developed with full transparency in mind so that others may freely use, adopt, and modify.

The work presented here is inspired heavily by classical stereotactic methods (J Talairach et al., 1957), where point-based correspondence has been used to align brain templates with patient anatomy to enable atlas-based surgical targeting. The anterior and posterior commissure were originally identified as prominent intraventricular features based on pneumoencephalography (air studies) and contrast ventriculography, prior to the invention of computed tomography or MRI. The AC and PC have proven to be reliable features on MRI and were adopted by neuroscientists for the alignment of brain images to templates, in what is referred to as the Talairach grid normalization procedure (Brett, Johnsrude, & Owen, 2002; A. Evans et al., 1992; Jean Talairach & Tournoux, 1988). The advent of robust and openly available software for automatic or semi-automatic labeling of regions-of-interest in brain images has led to a relative underemphasis of point-based alignment. We demonstrate here that point-based metrics are more sensitive to focal misregistrations than voxel overlap measures and quantified in millimeters.

Tolerance to focal misregistration in images undoubtedly will depend on the application; but there is no doubt that poor image correspondence can result in inaccurate (and possibly erroneous) predictions and conclusions in neuroimaging studies. Our results evaluating correspondence error in an fMRI preprocessing pipeline revealed local template misregistrations of 1.80 +/- 2.09 mm. For many fMRI or diffusion-based applications, this mean error is about the size of a voxel; and thus may be within an acceptable tolerance. However, mean maximum errors of over 7 mm were also observed and may begin to impact the sensitivity to discovery as well as the accuracy of localization of affected brain regions in a task or connectivity analyses. These misregistrations also may affect the interpretation of voxel-based and deformation-based morphometry studies that seek to investigate subtle shape differences between study populations. Finally, minimizing registration error becomes particularly critical for analyses pertaining to stereotactic interventions like deep brain stimulation (DBS) where millimeters can represent the difference between optimal therapy and side effects.

### Protocol development and validation

After a single training session, novice raters could place AFIDs at a mean AFLE of approximately 1-1.5 mm across all AFID points. Placement error varied from one template to another and among AFIDs (Supporting Information S1 file). Raters had the least amount of error with placements for the MNI2009bAsym and Agile12v2016 templates. In contrast, fiducial placement errors were higher when raters were asked to place AFIDs for individual subjects, i.e. Colin27 as well as the OASIS-1 database. Repeatability was assessed using measures of intra-rater and inter-rater AFLE. Intra-rater AFLE was lowest for the MNI2009bAsym and highest in Colin27 (Table 2). Inter-rater AFLE was again lowest for MNI2009bAsym and highest in Colin27 and the OASIS-1 datasets. This demonstrates how AFIDs are more difficult to place due to individual variability versus in population templates where the individual nuances of these features may be effectively blurred out. Overall, the placement error remains acceptable (1-2 mm) among all annotated images.

The AC and PC were the most reliably identifiable AFIDs with mean AFLE of less than 0.5 mm and inter-rater AFLE of 0.5-1 +/- 0.3 mm observed. These results compared favorably to an analysis of experienced neurosurgeons by Pallaravam and colleagues placing the same AC-PC points where they observed a point placement error (equivalent to the inter-rater AFLE metric used here) that was surprisingly higher at 1-2 mm +/- 1.5 mm (Pallavaram et al., 2008). We speculate that the higher variability in the referenced study was the lack of restriction on how the AC-PC landmarks were placed; that is, some stereotactic neurosurgeons continue to use the intraventricular edge of each commissure, which was the classical technique used by Talairach during air studies, while others used the center of each commissure (Horn et al., 2017). The distance from the center to the ventricular edge can be several millimeters likely accounting for this difference. Overall, our findings demonstrate that enforcing certain practices such as using the center of each commissure play an important role in the consistency and standardization of fiducial placement.

In contrast, certain fiducial points contributed substantially to worse overall estimates of fiducial localization error. The ventricular features in general had higher placement errors than other regions. In particular, the bilateral ventral occipital horns (AFID29-30) had higher placement errors. Placement was particularly inaccurate for individual subjects where the ventricular atrium tapered completely in many individual subject studies (including Colin27), and thus the posterior continuation into the occipital horn was sometimes difficult to visualize or resolve at all. The bilateral origins of the indusium griseum (AFID27-28) were also difficult for raters to place consistently. Less accuracy likely relate to features that are less salient than other regions and those likely exhibiting higher anatomical variability from one subject to the next.

### Point-based versus ROI-based metrics

Previous work has shown that nonlinear registration improves alignment between structures (Chakravarty et al., 2009; Hellier et al., 2003; Klein et al., 2009), and that the choice of parameters matters. These existing studies have mostly used voxel overlap measures to support their findings. Our results are also in-line with prior work but also demonstrate how AFIDs are complementary and more sensitive than ROI-based metrics for evaluating both local and global spatial correspondence of brain images (see Fig 5).

We were able to compare the relative efficacy of AFRE and voxel overlap for subjects from the OASIS-1 database and several commonly used templates. AFRE had a more unimodal distribution and a longer tail facilitating identification of focal misregistrations between images (Fig 5). On the other hand, the Jaccard histogram was more sparse towards the tail of the distribution suggesting a poorer ability to discriminate. One key advantage of AFRE is its interpretability, representing the distance in millimeters between aligned neuroanatomical structures in two images, compared to voxel overlap, which is a relative measure and unitless. It is commonly perceived in segmentation studies that voxel overlap measures greater than 0.7 represent accurate correspondence between regions. However, our analysis demonstrates that even with generally high overlap after nonlinear registration, focal misregistrations of AFIDs above 7 mm may be identified (Fig 6 and Table 5). Comparing AFRE against other registration quality metrics such as spatial cross-correlation and mutual information is beyond the scope of the current work.

### Subject-to-template registration

We chose to evaluate the subject-to-template registrations computed as part of an fMRI processing pipeline, fMRIPrep (Esteban et al., 2018), as a use case for our AFIDs protocol. Functional MRI studies may not represent the optimal use case due to the relatively coarse spatial resolution relative to the size of misregistration effects we can detect with AFIDs, and because most fMRI researchers are focused on cortical activation while our protocol emphasizes and detects misregistrations in the deep brain regions. Our choice to investigate fMRIPrep registration performance was motivated by their transparent approach to the development of preprocessing software for neuroimaging and BIDS integration (Gorgolewski et al., 2017, 2016). The active developer and support base, as well as growing adoption by many end-users were other contributing factors. Our analysis revealed misregistrations on the order of 1.80 +/- 2.09 mm and as high as over 30 mm that would be more difficult to identify by qualitative evaluation or ROI-based analysis alone.

While this points to potential caution with the use of standardized pipelines like fMRIPrep for template registration, it should be noted that fMRIPrep was designed with a focus on robustness, rather than accuracy. The underlying parameters and processing steps used in fMRIPrep are fully transparent. In addition, the underlying deformable registration software used (Avants et al., 2008) has been demonstrated to achieve high performance in studies using traditional voxel overlap measures (Klein et al., 2009). The focal template misregistrations we have identified in fMRIPrep with AFIDs are meant to serve as a baseline for refinement in future versions that can be compared transparently and potentially incorporated for testing new versions as part of a continuous integration workflow. Using additional image contrasts (Xiao et al., 2017) or subcortical tissue priors (Ewert et al., 2019) to drive template registration have been demonstrated using conventional voxel overlap techniques to result in more optimal registrations that can also be tested using the AFIDs framework.

### Template-to-template registration

We recommend that imaging scientists exercise caution when displaying statistical maps using a template other than the one to which the original deformations were performed. For example, it has become increasingly common to project statistical maps and subject data registered to MNI space using BigBrain for visualization purposes. In this study, we identified clear evidence of registration differences between several templates commonly assumed to be in the same coordinate space: BigBrainSym and MNI2009bSym, and even greater between BigBrainSym and MNI2009bAsym because of the differences in AFID locations in MNI2009bSym and MNI2009bAsym. Specifically, misregistrations as high as over 9 mm have been identified. Many of these errors occur in the midbrain region (Table 6), which would have implications in particular if using BigBrainSym to project locations of electrode implantations. In support of other recent work (Horn et al., 2017), this study highlights the importance of understanding which exact template one is using for processing and analysis: that multiple “MNI” templates exist (with different version dates, types, and symmetry), as do registration differences between these templates.

### Teaching neuroanatomy

Our protocol may also hold particular value for teaching neuroanatomy. In fact, evidence from our study suggests that even relative novices can be trained to place AFIDs accurately, including the AC and PC, with comparable accuracy and variability to trained neurosurgeons (Table 3). By releasing the data acquired in this study, we provide a normative distribution of AFID placements that can be used to quantify how accurately new trainees can place points. These measures can be used to gauge the comprehension of students regarding the specific location of neuroanatomical structures in a quantitative (millimetric) manner and focus efforts on consolidating understanding based on where localization errors were higher. To date, over a series of locally-held workshops and tutorials, over 60 students have been trained to complete the AFIDs protocol. Finally, the online AFIDs validator will facilitate larger scale training. Trainees will be able to check their work and become confident with the protocol by comparing against ground truth labels before using it on their own data.

### Limitations and future work

While we have found the AFIDs proposed to be quite reliable, there is clearly location-related heterogeneity in placement error. We make no claims that this set of anatomical fiducials is optimal and in the future, other locations may prove to be more effective than others. Also, for this first proposed set of AFIDs, we limited our locations to deep structures where less inter-subject variability exists compared to cortical features (Thompson et al., 1996); future extensions could include linking our workflow with cortical surface-based (B. Fischl, 2004) and sulcal-based (Hellier et al., 2003; Mangin et al., 2015; Perrot, Rivière, & Mangin, 2011) methods of spatial correspondence. Development of similar protocols for other neuroimaging modalities such as T2-weighted or diffusion-based contrasts may also be of value. In addition, fiducial localization error may be biased by how the raters were taught to place the fiducials; in our case, we organized an initial interactive tutorial session, and provided text and picture-based resources of how to place the AFIDs. It is also possible that AFLE would be lower if performed by a more experienced group of raters. Also, how AFID placement behaves in the presence of lesional pathology remains an open question. We have made the annotations and images available to allow other groups to propose other AFID locations and descriptions that could be similarly validated. We plan to post any modifications to the protocol as separate versions at the linked repository.

The AFIDs protocol requires correct placement of the anterior commissure (AFID01) and posterior commissure (AFID02) points. We made this decision as it helps to align the brain images into a more standard orientation for subsequent placement of bilateral fiducials. In particular, 4 of the AFIDs are dependent on AC-PC alignment (the lateral ventricles at AC and PC in the coronal plane). In fact, we found on secondary analysis that that error in AFID placements could be compounded by initial error in placement of AC and PC (see Supporting Information S1 file). Fortunately, AC and PC can be placed with high trueness and precision (< 1 mm) (Table 3), consistent with prior studies (Liu & Dawant, 2015). We made the decision to perform AC-PC alignment to permit more accurate placement of lateral AFIDs, which may otherwise have appeared quite oblique from each other if the individual’s head was tilted in the scanner. Thus, on balance, AC-PC alignment probably mitigates placement error in lateral AFIDs compared to placing fiducials in the native MRI space.

Beyond evaluating correspondence, AFIDs could be used for point-based inter-subject or subject-to-template registration. AFIDs used in combination with classic rigid registration algorithms such as Iterative Closest Point (Besl & McKay, 1992) may result in more optimal initial linear registration between images. In addition, point-based deformable registration using (B-splines) may produce more efficient, lower order deformable registrations between two images (Bookstein, 1997). To prevent circular reasoning, we thought this would be best evaluated as independent studies. Finally one compelling extension of this work would be to automate or semi-automate AFID placement, which would enable inclusion of AFID-based metrics in standardized workflows involving template or intersubject registration.

## Conclusions

Our proposed framework consists of the identification of anatomical fiducials, AFIDs, in structural magnetic resonance images of the human brain. Validity has been established using several openly available brain templates and datasets. We found that novice users could be trained to reliably place these points over a series of interactive training sessions to within millimeters of placement accuracy. As an example of different use cases, we examined the utility of our proposed protocol for evaluating subject-to-template and template-to-template registration revealing that AFIDs are sensitive to focal misregistrations that may be missed using other commonly used evaluation methods. This protocol holds value for a broad number of applications including intersubject alignment and teaching neuroanatomy.

## Acknowledgements

The authors would like to thank the many students whose participation in the neuroanatomy tutorials and workshops provided a foundation for the AFIDs framework. JCL is funded through the Western University Clinical Investigator Program accredited by the Royal College of Physicians and Surgeons of Canada and a Canadian Institutes of Health Research Frederick Banting and Charles Best Canada Graduate Doctoral Award Scholarship. Funding for this project was also provided by the Canadian Institute for Health Research CIHR FDN 201409. Infrastructural support was provided by the Canada First Research Excellence Fund to BrainsCAN, Brain Canada, and computational resource through Compute Canada.

## Supporting Information

Additional Supporting Information may be found online in the supporting information tab for this article.

S1 File. Phase 1 Notebook.

S2 File. Phase 2 Notebook.

S3 File. Phase 3 Notebook.

S4 File. Phase 4 Notebook.

S5 File. AFIDS Protocol.

