## Supplementary material for "A framework for evaluating correspondence between brain images using anatomical fiducials": S1_PHASE1_template_validation.pdf

### Phase 1: Protocol Validation for Brain Templates

This notebook contains results validating the AFID protocol on three openly available templates ( `Agile12v2016` , `Colin27` , and `ICBM2009bAsym` ).

The first step is to initialize the variables, define useful functions, and load all the raw fcsv data into `df_raters` .

##### Template Averages

For each template, we calculate the mean value for each AFID32 point and store it in a separate .fcsv file so that it can be loaded back into 3D Slicer.

Deviation of the values by > 10 mm will be classified as an outlier.

##### Phase 1: Raw Data Analysis

Also classify extreme outliers, defined as >= 10 mm from the group mean

'Total: 1.27 +/- 1.98 mm; Outliers: 24/3072 (0.78%)'

'Agile12v2016: 1.10 +/- 1.59 mm; Outliers: 3/1024 (0.29%)'

'Colin27: 1.71 +/- 2.78 mm; Outliers: 20/1024 (1.95%)'

'MNI152NLin2009bAsym: 0.99 +/- 1.11 mm; Outliers: 1/1024 (0.10%)'

### Template Averages: Post-QC

Template averages were recreated after quality control and filtering of outliers.

'Total: 1.03 +/- 0.94 mm; Outliers: 1/3048 (0.03%)'

'Agile12v2016: 1.01 +/- 0.93 mm; Outliers: 0/1021 (0.00%)'

'Colin27: 1.11 +/- 1.05 mm; Outliers: 1/1004 (0.10%)'

'MNI152NLin2009bAsym: 0.97 +/- 0.80 mm; Outliers: 0/1023 (0.00%)'

| AFID | Description | Agile12v2016 Pre-QC | Agile12v2016 Post-QC | Colin27 Pre-QC | Colin27 Post-QC | MNI2009bAsym Pre-QC | MNI2009bAsym Post-QC | Total Pre-QC | Total Post-QC |
|---|---|---|---|---|---|---|---|---|---|
| 01 | AC | 0.33±0.16 (0) | 0.33±0.16 (0) | 0.34±0.29 (0) | 0.34±0.29 (0) | 0.35±0.20 (0) | 0.35±0.20 (0) | 0.34±0.22 (0) | 0.34±0.22 (0) |
| 02 | PC | 0.34±0.19 (0) | 0.34±0.19 (0) | 0.35±0.18 (0) | 0.35±0.18 (0) | 0.33±0.14 (0) | 0.33±0.14 (0) | 0.34±0.17 (0) | 0.34±0.17 (0) |
| 03 | infracollicular sulcus | 1.25±0.47 (0) | 1.25±0.47 (0) | 1.22±0.48 (0) | 1.22±0.48 (0) | 1.08±0.46 (0) | 1.08±0.46 (0) | 1.17±0.47 (0) | 1.17±0.47 (0) |
| 04 | PMJ | 0.83±0.47 (0) | 0.83±0.47 (0) | 0.97±0.65 (0) | 0.97±0.65 (0) | 0.84±0.52 (0) | 0.84±0.52 (0) | 0.87±0.54 (0) | 0.87±0.54 (0) |
| 05 | superior interpeduncular fossa | 1.15±0.61 (0) | 1.15±0.61 (0) | 0.96±0.60 (0) | 0.96±0.60 (0) | 1.12±0.50 (0) | 1.12±0.50 (0) | 1.08±0.57 (0) | 1.08±0.57 (0) |
| 06 | R superior LMS | 0.75±0.48 (0) | 0.75±0.48 (0) | 1.16±0.69 (0) | 1.16±0.69 (0) | 0.68±0.50 (0) | 0.68±0.50 (0) | 0.85±0.59 (0) | 0.85±0.59 (0) |
| 07 | L superior LMS | 0.93±0.59 (0) | 0.93±0.59 (0) | 1.05±0.57 (0) | 1.05±0.57 (0) | 0.91±0.90 (0) | 0.91±0.90 (0) | 0.96±0.71 (0) | 0.96±0.71 (0) |
| 08 | R inferior LMS | 1.55±1.14 (0) | 1.55±1.14 (0) | 1.61±1.07 (0) | 1.61±1.07 (0) | 1.47±0.96 (0) | 1.47±0.96 (0) | 1.54±1.05 (0) | 1.54±1.05 (0) |
| 09 | L inferior LMS | 1.39±1.11 (0) | 1.39±1.11 (0) | 1.79±1.32 (0) | 1.79±1.32 (0) | 1.63±1.19 (0) | 1.63±1.19 (0) | 1.60±1.21 (0) | 1.60±1.21 (0) |
| 10 | culmen | 1.03±0.73 (0) | 1.03±0.73 (0) | 0.68±0.24 (0) | 0.68±0.24 (0) | 0.61±0.32 (0) | 0.61±0.32 (0) | 0.77±0.50 (0) | 0.77±0.50 (0) |
| 11 | intermammillary sulcus | 0.73±0.34 (0) | 0.73±0.34 (0) | 0.68±0.34 (0) | 0.68±0.34 (0) | 0.70±0.38 (0) | 0.70±0.38 (0) | 0.70±0.35 (0) | 0.70±0.35 (0) |
| 12 | R MB | 0.37±0.28 (0) | 0.37±0.28 (0) | 0.44±0.32 (0) | 0.44±0.32 (0) | 0.48±0.34 (0) | 0.48±0.34 (0) | 0.44±0.31 (0) | 0.44±0.31 (0) |
| 13 | L MB | 0.43±0.27 (0) | 0.43±0.27 (0) | 0.53±0.32 (0) | 0.53±0.32 (0) | 0.50±0.31 (0) | 0.50±0.31 (0) | 0.49±0.30 (0) | 0.49±0.30 (0) |
| 14 | pineal gland | 0.70±0.33 (0) | 0.70±0.33 (0) | 0.94±0.33 (0) | 0.94±0.33 (0) | 0.68±0.51 (0) | 0.68±0.51 (0) | 0.77±0.42 (0) | 0.77±0.42 (0) |
| 15 | R LV at AC | 0.99±1.48 (0) | 0.99±1.48 (0) | 0.68±0.42 (0) | 0.68±0.42 (0) | 0.62±0.50 (0) | 0.62±0.50 (0) | 0.75±0.92 (0) | 0.75±0.92 (0) |
| 16 | L LV at AC | 1.06±1.60 (0) | 1.06±1.60 (0) | 0.73±0.42 (0) | 0.73±0.42 (0) | 0.62±0.51 (0) | 0.62±0.51 (0) | 0.79±0.98 (0) | 0.79±0.98 (0) |
| 17 | R LV at PC | 1.13±1.35 (0) | 1.13±1.35 (0) | 1.12±1.01 (0) | 1.12±1.01 (0) | 1.00±0.60 (0) | 1.00±0.60 (0) | 1.08±1.00 (0) | 1.08±1.00 (0) |
| 18 | L LV at PC | 1.23±1.46 (0) | 1.23±1.46 (0) | 1.32±1.02 (0) | 1.32±1.02 (0) | 1.03±0.58 (0) | 1.03±0.58 (0) | 1.18±1.05 (0) | 1.18±1.05 (0) |
| 19 | genu of CC | 1.00±0.46 (0) | 1.00±0.46 (0) | 0.63±0.24 (0) | 0.63±0.24 (0) | 0.78±0.48 (0) | 0.78±0.48 (0) | 0.80±0.44 (0) | 0.80±0.44 (0) |
| 20 | splenium of CC | 0.71±0.39 (0) | 0.71±0.39 (0) | 0.52±0.27 (0) | 0.52±0.27 (0) | 0.80±1.10 (0) | 0.80±1.10 (0) | 0.68±0.73 (0) | 0.68±0.73 (0) |
| 21 | R AL temporal horn | 1.44±1.20 (0) | 1.44±1.20 (0) | 1.52±0.79 (0) | 1.52±0.79 (0) | 1.15±0.89 (0) | 1.15±0.89 (0) | 1.36±0.98 (0) | 1.36±0.98 (0) |
| 22 | L AL temporal horn | 1.64±1.92 (1) | 1.32±0.91 (0) | 1.10±0.56 (0) | 1.10±0.56 (0) | 1.16±0.94 (0) | 1.16±0.94 (0) | 1.29±1.27 (1) | 1.19±0.82 (0) |
| 23 | R superior AM temporal horn | 0.62±0.38 (0) | 0.62±0.38 (0) | 1.31±1.71 (0) | 1.31±1.71 (0) | 0.83±0.91 (0) | 0.83±0.91 (0) | 0.91±1.15 (0) | 0.91±1.15 (0) |
| 24 | L superior AM temporal horn | 0.59±0.39 (0) | 0.59±0.39 (0) | 2.02±1.90 (0) | 2.02±1.90 (0) | 0.95±0.98 (0) | 0.95±0.98 (0) | 1.17±1.36 (0) | 1.17±1.36 (0) |
| 25 | R inferior AM temporal horn | 1.31±1.20 (0) | 1.31±1.20 (0) | 1.49±0.94 (0) | 1.49±0.94 (0) | 1.40±1.00 (0) | 1.40±1.00 (0) | 1.40±1.04 (0) | 1.40±1.04 (0) |
| 26 | L inferior AM temporal horn | 1.36±1.16 (0) | 1.36±1.16 (0) | 1.41±1.06 (0) | 1.41±1.06 (0) | 1.39±0.76 (0) | 1.39±0.76 (0) | 1.38±0.98 (0) | 1.38±0.98 (0) |
| 27 | R indusium griseum origin | 2.58±4.99 (1) | 1.38±0.75 (0) | 1.70±1.08 (0) | 1.70±1.08 (0) | 1.26±0.82 (0) | 1.26±0.82 (0) | 1.81±2.92 (1) | 1.43±0.90 (0) |
| 28 | L indusium griseum origin | 2.57±4.91 (1) | 1.52±1.14 (0) | 2.11±1.44 (0) | 2.11±1.44 (0) | 1.40±0.98 (0) | 1.40±0.98 (0) | 1.99±2.94 (1) | 1.66±1.22 (0) |
| 29 | R ventral occipital horn | 1.59±1.07 (0) | 1.59±1.07 (0) | 11.38±4.82 (10) | 0.80±0.45 (0) | 2.07±4.11 (1) | 1.34±1.25 (0) | 4.84±5.78 (11) | 1.30±1.08 (0) |
| 30 | L ventral occipital horn | 1.09±1.13 (0) | 1.09±1.13 (0) | 10.04±4.78 (10) | 1.63±2.94 (1) | 1.25±1.32 (0) | 1.25±1.32 (0) | 3.96±5.02 (10) | 1.28±1.78 (1) |
| 31 | R olfactory sulcal fundus | 1.17±0.68 (0) | 1.17±0.68 (0) | 1.41±0.95 (0) | 1.41±0.95 (0) | 1.14±0.59 (0) | 1.14±0.59 (0) | 1.23±0.75 (0) | 1.23±0.75 (0) |
| 32 | L olfactory sulcal fundus | 1.23±0.54 (0) | 1.23±0.54 (0) | 1.41±1.00 (0) | 1.41±1.00 (0) | 1.13±0.63 (0) | 1.13±0.63 (0) | 1.25±0.74 (0) | 1.25±0.74 (0) |

### Demographics of Raters

Details regarding experience, etc.

| rater_id | imaging_exp | neuro_exp | slicer_exp | description |
|---|---|---|---|---|
| Rater01 | 24 | 24 | 24 | undergrad_student |
| Rater02 | 0 | 0 | 0 | medical_student |
| Rater03 | 8 | 0 | 8 | undergrad_student |
| Rater04 | 24 | 6 | 0 | grad_student |
| Rater05 | 0 | 24 | 0 | grad_student |
| Rater06 | 24 | 12 | 12 | grad_student |
| Rater07 | 12 | 48 | 12 | grad_student |
| Rater08 | 0 | 0 | 0 | undergrad_student |

'Imaging Experience: 11.5 +/- 11.2 months (Range: 0.0-24.0)'

'Neuroanatomy Experience: 14.2 +/- 17.0 months (Range: 0.0-48.0)'

'3D Slicer Experience: 7.0 +/- 8.8 months (Range: 0.0-24.0)'

#### Secondary Analyses

We first evaluated whether there was any evidence of learning across sessions (excluding session 0 which was completed as part of a group tutorial). There were negative trends in the mean AFLE with increasing session number but these did not meet thresholds of statistical analysis. The first column is the effect, second column is the associated p-value.

Summary text on learning: As a secondary analysis, we explored whether any evidence of learning over the 4 independent rating sessions could be identified (Supporting Information S1 file). Using linear modeling, we identified a general decrease in mean AFLE with increasing session number although this did not meet thresholds of statistical significance (estimate = -0.02 mm/session; p-value = 0.11). These trends were explored on the individual rater level. For two out of 8 raters, AFLE varied with session number. Rater04 demonstrated a general linear improvement of -0.17 mm/session from an initial mean AFLE of 1.64 mm (i.e. the worst performing initial session); however Rater02 worsened at a rate of 0.12 mm/session from an initial mean AFLE of 0.59 mm (i.e. the best performing initial session). No significant effect with individual AFIDs was identified. All subgroup analyses were multiple comparisons corrected using FDR (q-value < 0.05).

| | | |
|---|---|---|
| **(Intercept)** | 1.090 | 0.0000 |
| **session** | -0.024 | 0.1141 |

##### Did specific raters demonstrate any learning?

Because of the trends, we explored further to determine whether any specific raters demonstrated any learning. After multiple comparisons correction, only two raters demonstrated statistically significant change with session number. Rater #4 was observed to start at a baseline increased rating error in the first session (1.64 mm) but demonstrated a decrease in AFLE with session number improving by 0.1-0.2 mm per session based on the linear model (statistically significant improvement). On the contrary, Rater #2 who started with an intercept of 0.59 mm (better than the average) showed worsening of rater error with time.

| rater | (Intercept) | pval_(Intercept) | session | pval_session | pval_session_adjusted | pval_session_significant |
|---|---|---|---|---|---|---|
| 1 | 0.78 | 0 | 0.02 | 0.5644 | 0.6450 | FALSE |
| 2 | 0.59 | 0 | 0.12 | 0.0001 | 0.0009 | TRUE |
| 3 | 1.30 | 0 | -0.06 | 0.1624 | 0.2783 | FALSE |
| 4 | 1.64 | 0 | -0.17 | 0.0002 | 0.0009 | TRUE |
| 5 | 1.05 | 0 | 0.04 | 0.2881 | 0.3841 | FALSE |
| 6 | 1.08 | 0 | -0.04 | 0.1739 | 0.2783 | FALSE |
| 7 | 0.84 | 0 | 0.02 | 0.7086 | 0.7086 | FALSE |
| 8 | 1.45 | 0 | -0.12 | 0.0210 | 0.0560 | FALSE |

##### Did AFLE improve for specific AFIDs?

We wanted to see if specific AFIDs tended to improve with more training (i.e. more sessions). This analysis did not survive multiple comparisons analysis.

| fid | (Intercept) | pval_(Intercept) | session | pval_session | pval_session_adjusted | pval_session_significant |
|---|---|---|---|---|---|---|
| 1 | 0.36 | 0.0000 | -0.01 | 0.7009 | 0.8307 | FALSE |
| 2 | 0.38 | 0.0000 | -0.01 | 0.3531 | 0.7955 | FALSE |
| 3 | 1.28 | 0.0000 | -0.03 | 0.4534 | 0.7955 | FALSE |
| 4 | 0.65 | 0.0000 | 0.09 | 0.0588 | 0.4391 | FALSE |
| 5 | 1.08 | 0.0000 | 0.00 | 0.9802 | 0.9802 | FALSE |
| 6 | 0.71 | 0.0000 | 0.06 | 0.2590 | 0.7296 | FALSE |
| 7 | 0.92 | 0.0000 | 0.01 | 0.8661 | 0.8941 | FALSE |
| 8 | 1.76 | 0.0000 | -0.08 | 0.4408 | 0.7955 | FALSE |
| 9 | 1.82 | 0.0000 | -0.07 | 0.5221 | 0.7955 | FALSE |
| 10 | 0.79 | 0.0000 | -0.01 | 0.8035 | 0.8866 | FALSE |
| 11 | 0.85 | 0.0000 | -0.06 | 0.0686 | 0.4391 | FALSE |
| 12 | 0.47 | 0.0000 | -0.01 | 0.6396 | 0.8307 | FALSE |
| 13 | 0.41 | 0.0000 | 0.03 | 0.2885 | 0.7296 | FALSE |
| 14 | 1.02 | 0.0000 | -0.10 | 0.0076 | 0.2431 | FALSE |
| 15 | 0.68 | 0.0051 | 0.03 | 0.6932 | 0.8307 | FALSE |
| 16 | 0.74 | 0.0044 | 0.03 | 0.7880 | 0.8866 | FALSE |
| 17 | 1.00 | 0.0002 | 0.04 | 0.6840 | 0.8307 | FALSE |
| 18 | 1.25 | 0.0000 | -0.02 | 0.8506 | 0.8941 | FALSE |
| 19 | 0.87 | 0.0000 | -0.02 | 0.5664 | 0.8239 | FALSE |
| 20 | 0.86 | 0.0000 | -0.09 | 0.0188 | 0.3005 | FALSE |
| 21 | 1.22 | 0.0000 | 0.07 | 0.4391 | 0.7955 | FALSE |
| 22 | 1.05 | 0.0000 | 0.07 | 0.3945 | 0.7955 | FALSE |
| 23 | 1.21 | 0.0000 | -0.13 | 0.1677 | 0.7296 | FALSE |
| 24 | 1.34 | 0.0001 | -0.08 | 0.5017 | 0.7955 | FALSE |
| 25 | 1.76 | 0.0000 | -0.13 | 0.1829 | 0.7296 | FALSE |
| 26 | 1.64 | 0.0000 | -0.10 | 0.2964 | 0.7296 | FALSE |
| 27 | 1.04 | 0.0000 | 0.15 | 0.0629 | 0.4391 | FALSE |
| 28 | 1.47 | 0.0000 | 0.08 | 0.4954 | 0.7955 | FALSE |
| 29 | 1.79 | 0.0000 | -0.18 | 0.0984 | 0.5246 | FALSE |
| 30 | 1.82 | 0.0004 | -0.20 | 0.2627 | 0.7296 | FALSE |
| 31 | 1.46 | 0.0000 | -0.09 | 0.2158 | 0.7296 | FALSE |
| 32 | 1.33 | 0.0000 | -0.03 | 0.6341 | 0.8307 | FALSE |

#### Impact of AC and PC on AFLE

Analysis completed at the request of reviewers. We identified that the global AFLE values varied linearly with the accuracy of AC placement but not PC placement.

```
Call:
lm(formula = mean_AFLE ~ AC, data = df_ACPC_impact)

Residuals:
     Min       1Q   Median       3Q      Max
-0.38205 -0.16274 -0.02348  0.11585  0.92690

Coefficients:
            Estimate Std. Error t value Pr(>|t|)
(Intercept)  0.86087    0.04325  19.903  < 2e-16 ***
AC           0.49349    0.10538   4.683 9.51e-06 ***
---
Signif. codes:  0 ‘***’ 0.001 ‘**’ 0.01 ‘*’ 0.05 ‘.’ 0.1 ‘ ’ 1

Residual standard error: 0.2302 on 94 degrees of freedom
Multiple R-squared:  0.1892,    Adjusted R-squared:  0.1805
F-statistic: 21.93 on 1 and 94 DF,  p-value: 9.511e-06
```

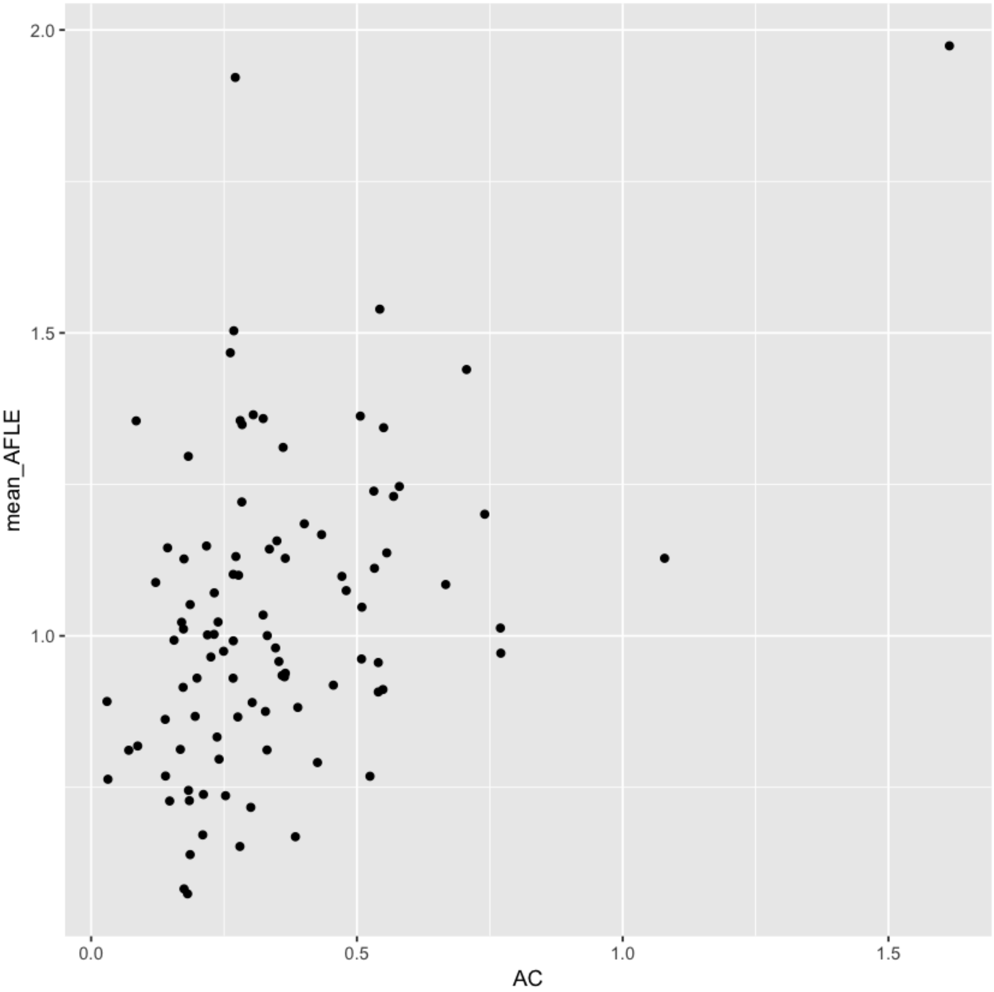

```
Call:
lm(formula = mean_AFLE ~ PC, data = df_ACPC_impact)

Residuals:
     Min       1Q   Median       3Q      Max 
-0.43827 -0.15525 -0.03379  0.11150  0.92972 

Coefficients:
            Estimate Std. Error t value Pr(>|t|)    
(Intercept)  0.97976    0.05771  16.977   <2e-16 ***
PC           0.14993    0.15099   0.993    0.323    
---
Signif. codes:  0 ‘***’ 0.001 ‘**’ 0.01 ‘*’ 0.05 ‘.’ 0.1 ‘ ’ 1

Residual standard error: 0.2543 on 94 degrees of freedom
Multiple R-squared:  0.01038,   Adjusted R-squared:  -0.0001473 
F-statistic: 0.986 on 1 and 94 DF,  p-value: 0.3233
```

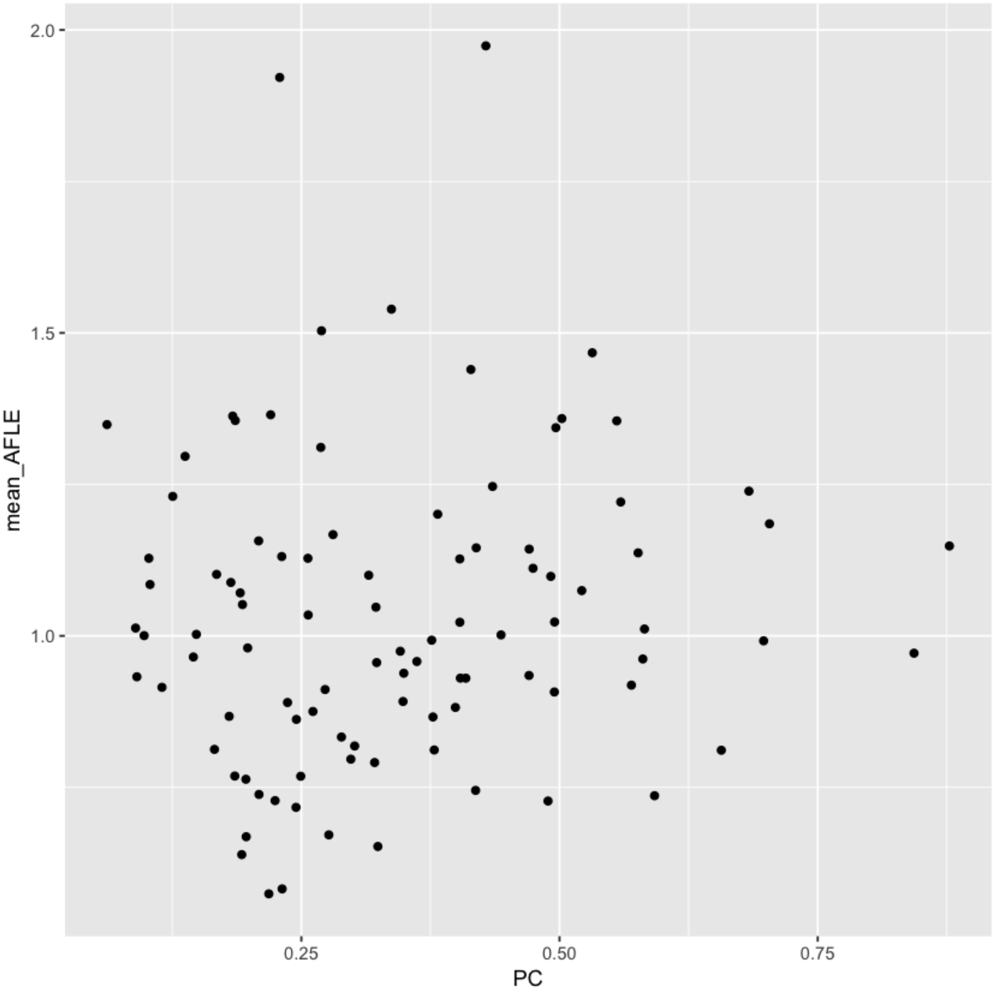

#### Intra-Rater AFLE

| template | mean | sd |
|---|---|---|
| Agile12v2016 | 1.13 | 0.86 |
| Colin27 | 1.14 | 0.92 |
| MNI152NLin2009bAsym | 1.03 | 0.78 |

'Intra-Rater AFLE: 1.10 +/- 0.86 mm'

#### Inter-Rater AFLE

| template | mean | sd |
|---|---|---|
| Agile12v2016 | 1.14 | 0.48 |
| Colin27 | 1.36 | 0.88 |
| MNI152NLin2009bAsym | 1.07 | 0.46 |

'Inter-Rater AFLE: 1.19 +/- 0.64 mm'

#### Summary of Validation Results (Post-QC)

Mean AFLE, Intra-Rater AFLE, Inter-Rater AFLE

| AFID | Description | Agile12v2016 Mean AFLE | Colin27 Mean AFLE | MNI2009bAsym Mean AFLE | Total Mean AFLE | Agile12v2016 Intra-Rater | Colin27 Intra-Rater | MNI2009bAsym Intra-Rater | Total Intra-Rater | Agile12v2016 Inter-Rater | Colin27 Inter-Rater | MNI2009bAsym Inter-Rater | Total Inter-Rater |
|---|---|---|---|---|---|---|---|---|---|---|---|---|---|
| 01 | AC | 0.33±0.16 | 0.34±0.29 | 0.35±0.20 | 0.34±0.22 | 0.41±0.15 | 0.49±0.35 | 0.44±0.23 | 0.45±0.24 | 0.31±0.16 | 0.30±0.12 | 0.37±0.16 | 0.33±0.04 |
| 02 | PC | 0.34±0.19 | 0.35±0.18 | 0.33±0.14 | 0.34±0.17 | 0.43±0.22 | 0.42±0.13 | 0.39±0.17 | 0.41±0.17 | 0.33±0.10 | 0.39±0.13 | 0.34±0.13 | 0.35±0.03 |
| 03 | infracollicular sulcus | 1.25±0.47 | 1.22±0.48 | 1.08±0.46 | 1.17±0.47 | 0.93±0.36 | 0.70±0.32 | 0.70±0.40 | 0.78±0.36 | 1.60±0.72 | 1.67±0.76 | 1.47±0.75 | 1.58±0.10 |
| 04 | PMJ | 0.83±0.47 | 0.97±0.65 | 0.84±0.52 | 0.87±0.54 | 0.80±0.23 | 0.89±0.49 | 0.76±0.28 | 0.81±0.34 | 1.06±0.50 | 1.23±0.74 | 1.17±0.55 | 1.15±0.08 |
| 05 | superior interpeduncular fossa | 1.15±0.61 | 0.96±0.60 | 1.12±0.50 | 1.08±0.57 | 1.04±0.37 | 1.02±0.61 | 1.00±0.51 | 1.02±0.48 | 1.38±0.83 | 1.07±0.59 | 1.20±0.63 | 1.22±0.16 |
| 06 | R superior LMS | 0.75±0.48 | 1.16±0.69 | 0.68±0.50 | 0.85±0.59 | 1.07±0.38 | 1.44±0.50 | 0.91±0.46 | 1.14±0.48 | 0.63±0.28 | 1.17±0.59 | 0.55±0.34 | 0.78±0.34 |
| 07 | L superior LMS | 0.93±0.59 | 1.05±0.57 | 0.91±0.90 | 0.96±0.71 | 1.18±0.32 | 1.04±0.44 | 1.27±0.87 | 1.16±0.58 | 1.03±0.46 | 1.25±0.68 | 0.62±0.31 | 0.97±0.32 |
| 08 | R inferior LMS | 1.55±1.14 | 1.61±1.07 | 1.47±0.96 | 1.54±1.05 | 1.81±1.07 | 1.49±0.77 | 1.35±0.81 | 1.55±0.88 | 1.65±1.08 | 2.06±1.16 | 1.87±1.28 | 1.86±0.21 |
| 09 | L inferior LMS | 1.39±1.11 | 1.79±1.32 | 1.63±1.19 | 1.60±1.21 | 1.66±0.91 | 1.88±1.37 | 1.60±1.22 | 1.71±1.14 | 1.53±1.07 | 2.06±1.29 | 2.05±1.45 | 1.88±0.31 |
| 10 | culmen | 1.03±0.73 | 0.68±0.24 | 0.61±0.32 | 0.77±0.50 | 1.16±0.70 | 0.72±0.22 | 0.61±0.25 | 0.83±0.49 | 1.10±0.41 | 0.77±0.25 | 0.70±0.29 | 0.85±0.21 |
| 11 | intermammillary sulcus | 0.73±0.34 | 0.68±0.34 | 0.70±0.38 | 0.70±0.35 | 0.69±0.41 | 0.74±0.41 | 0.83±0.47 | 0.76±0.42 | 0.82±0.39 | 0.72±0.31 | 0.68±0.36 | 0.74±0.07 |
| 12 | R MB | 0.37±0.28 | 0.44±0.32 | 0.48±0.34 | 0.44±0.31 | 0.51±0.31 | 0.47±0.14 | 0.54±0.34 | 0.51±0.27 | 0.32±0.18 | 0.51±0.42 | 0.51±0.39 | 0.45±0.11 |
| 13 | L MB | 0.43±0.27 | 0.53±0.32 | 0.50±0.31 | 0.49±0.30 | 0.52±0.29 | 0.50±0.15 | 0.58±0.28 | 0.53±0.24 | 0.40±0.19 | 0.63±0.45 | 0.52±0.33 | 0.52±0.12 |
| 14 | pineal gland | 0.70±0.33 | 0.94±0.33 | 0.68±0.51 | 0.77±0.42 | 0.91±0.24 | 1.16±0.37 | 0.83±0.57 | 0.97±0.42 | 0.63±0.25 | 0.70±0.41 | 0.68±0.35 | 0.67±0.04 |
| 15 | R LV at AC | 0.99±1.48 | 0.68±0.42 | 0.62±0.50 | 0.75±0.92 | 1.29±1.50 | 0.74±0.41 | 0.74±0.45 | 0.92±0.93 | 1.10±0.66 | 0.81±0.35 | 0.73±0.30 | 0.88±0.20 |
| 16 | L LV at AC | 1.06±1.60 | 0.73±0.42 | 0.62±0.51 | 0.79±0.98 | 1.34±1.50 | 0.76±0.33 | 0.78±0.41 | 0.96±0.92 | 1.31±1.09 | 0.91±0.35 | 0.77±0.32 | 0.99±0.28 |
| 17 | R LV at PC | 1.13±1.35 | 1.12±1.01 | 1.00±0.60 | 1.08±1.00 | 1.35±1.35 | 1.19±1.04 | 0.90±0.60 | 1.14±1.01 | 1.29±0.65 | 1.38±0.59 | 1.32±0.58 | 1.33±0.05 |
| 18 | L LV at PC | 1.23±1.46 | 1.32±1.02 | 1.03±0.58 | 1.18±1.05 | 1.48±1.29 | 1.18±1.07 | 0.91±0.54 | 1.19±1.00 | 1.50±1.04 | 1.65±0.72 | 1.40±0.61 | 1.52±0.13 |
| 19 | genu of CC | 1.00±0.46 | 0.63±0.24 | 0.78±0.48 | 0.80±0.44 | 1.10±0.66 | 0.62±0.21 | 0.84±0.42 | 0.85±0.49 | 0.99±0.46 | 0.75±0.28 | 0.99±0.48 | 0.91±0.14 |
| 20 | splenium of CC | 0.71±0.39 | 0.52±0.27 | 0.80±1.10 | 0.68±0.73 | 0.90±0.40 | 0.69±0.21 | 0.86±0.52 | 0.81±0.39 | 0.67±0.32 | 0.47±0.15 | 0.67±0.29 | 0.60±0.11 |
| 21 | R AL temporal horn | 1.44±1.20 | 1.52±0.79 | 1.15±0.89 | 1.36±0.98 | 1.55±1.26 | 1.71±0.59 | 1.33±1.12 | 1.53±1.00 | 1.72±1.00 | 1.65±0.80 | 1.32±0.67 | 1.56±0.21 |
| 22 | L AL temporal horn | 1.32±0.91 | 1.10±0.56 | 1.16±0.94 | 1.19±0.82 | 1.32±1.07 | 1.29±0.39 | 1.49±0.93 | 1.36±0.82 | 1.46±0.91 | 1.15±0.52 | 1.31±0.62 | 1.30±0.16 |
| 23 | R superior AM temporal horn | 0.62±0.38 | 1.31±1.71 | 0.83±0.91 | 0.91±1.15 | 0.70±0.36 | 1.73±1.90 | 0.72±0.33 | 1.05±1.19 | 0.69±0.37 | 1.35±0.74 | 0.80±0.34 | 0.95±0.35 |
| 24 | L superior AM temporal horn | 0.59±0.39 | 2.02±1.90 | 0.95±0.98 | 1.17±1.36 | 0.66±0.31 | 2.42±2.12 | 0.80±0.26 | 1.29±1.44 | 0.71±0.36 | 2.08±1.07 | 0.89±0.32 | 1.23±0.74 |
| 25 | R inferior AM temporal horn | 1.31±1.20 | 1.49±0.94 | 1.40±1.00 | 1.40±1.04 | 1.65±0.97 | 1.36±0.81 | 1.44±0.68 | 1.48±0.80 | 1.55±0.86 | 1.80±1.14 | 1.87±1.25 | 1.74±0.17 |
| 26 | L inferior AM temporal horn | 1.36±1.16 | 1.41±1.06 | 1.39±0.76 | 1.38±0.98 | 1.56±0.94 | 1.37±0.99 | 1.40±0.88 | 1.44±0.90 | 1.67±0.88 | 1.70±1.21 | 1.67±0.78 | 1.68±0.01 |
| 27 | R indusium griseum origin | 1.38±0.75 | 1.70±1.08 | 1.26±0.82 | 1.43±0.90 | 1.10±0.54 | 1.55±1.00 | 1.18±0.86 | 1.28±0.81 | 1.71±1.08 | 2.00±1.17 | 1.35±0.78 | 1.69±0.33 |
| 28 | L indusium griseum origin | 1.52±1.14 | 2.11±1.44 | 1.40±0.98 | 1.66±1.22 | 1.37±0.74 | 1.61±1.32 | 1.45±0.75 | 1.47±0.94 | 2.12±1.35 | 2.74±1.76 | 1.59±0.90 | 2.15±0.57 |
| 29 | R ventral occipital horn | 1.59±1.07 | 0.80±0.45 | 1.34±1.25 | 1.30±1.08 | 1.85±1.28 | 0.95±0.24 | 2.08±1.63 | 1.69±1.29 | 1.58±0.85 | 0.93±0.63 | 1.13±0.59 | 1.21±0.33 |
| 30 | L ventral occipital horn | 1.09±1.13 | 1.63±2.94 | 1.25±1.32 | 1.28±1.78 | 1.22±1.15 | 0.86±0.31 | 1.90±1.45 | 1.37±1.16 | 1.36±0.88 | 4.98±6.77 | 1.12±0.60 | 2.49±2.16 |
| 31 | R olfactory sulcal fundus | 1.17±0.68 | 1.41±0.95 | 1.14±0.59 | 1.23±0.75 | 1.44±0.75 | 1.91±0.65 | 1.23±0.56 | 1.53±0.69 | 0.96±0.51 | 1.35±0.68 | 1.20±0.69 | 1.17±0.19 |
| 32 | L olfactory sulcal fundus | 1.23±0.54 | 1.41±1.00 | 1.13±0.63 | 1.25±0.74 | 1.24±0.57 | 1.56±1.13 | 1.07±0.43 | 1.29±0.77 | 1.29±0.71 | 1.41±1.00 | 1.27±0.66 | 1.32±0.08 |

### ANOVA for Templates

A difference in placement error between templates was identified by ANOVA.

'F-value: 7.88; p-value: 0.0004'

| fid | Fval | pval | adjusted | significant |
|---|---|---|---|---|
| 1 | 0.03 | 0.9695 | 0.9767 | FALSE |
| 2 | 0.13 | 0.8760 | 0.9490 | FALSE |
| 3 | 1.45 | 0.2406 | 0.5918 | FALSE |
| 4 | 0.72 | 0.4900 | 0.7215 | FALSE |
| 5 | 1.01 | 0.3696 | 0.6571 | FALSE |
| 6 | 7.28 | 0.0011 | 0.0119 | TRUE |
| 7 | 0.38 | 0.6840 | 0.8755 | FALSE |
| 8 | 0.16 | 0.8535 | 0.9490 | FALSE |
| 9 | 0.90 | 0.4118 | 0.6935 | FALSE |
| 10 | 7.61 | 0.0008 | 0.0119 | TRUE |
| 11 | 0.12 | 0.8897 | 0.9490 | FALSE |
| 12 | 1.05 | 0.3546 | 0.6571 | FALSE |
| 13 | 0.84 | 0.4362 | 0.6979 | FALSE |
| 14 | 4.04 | 0.0206 | 0.1319 | FALSE |
| 15 | 1.57 | 0.2124 | 0.5918 | FALSE |
| 16 | 1.83 | 0.1659 | 0.5310 | FALSE |
| 17 | 0.17 | 0.8398 | 0.9490 | FALSE |
| 18 | 0.71 | 0.4960 | 0.7215 | FALSE |
| 19 | 6.38 | 0.0025 | 0.0198 | TRUE |
| 20 | 1.30 | 0.2779 | 0.5918 | FALSE |
| 21 | 1.39 | 0.2530 | 0.5918 | FALSE |
| 22 | 0.61 | 0.5467 | 0.7289 | FALSE |
| 23 | 3.19 | 0.0456 | 0.1826 | FALSE |
| 24 | 11.61 | 0.0000 | 0.0009 | TRUE |
| 25 | 0.24 | 0.7905 | 0.9490 | FALSE |
| 26 | 0.02 | 0.9767 | 0.9767 | FALSE |
| 27 | 2.22 | 0.1142 | 0.4060 | FALSE |
| 28 | 3.32 | 0.0401 | 0.1826 | FALSE |
| 29 | 3.76 | 0.0271 | 0.1443 | FALSE |
| 30 | 0.61 | 0.5433 | 0.7289 | FALSE |
| 31 | 1.28 | 0.2819 | 0.5918 | FALSE |
| 32 | 1.23 | 0.2959 | 0.5918 | FALSE |

#### K-means clustering of point cloud distributions

Across all templates; and template specific

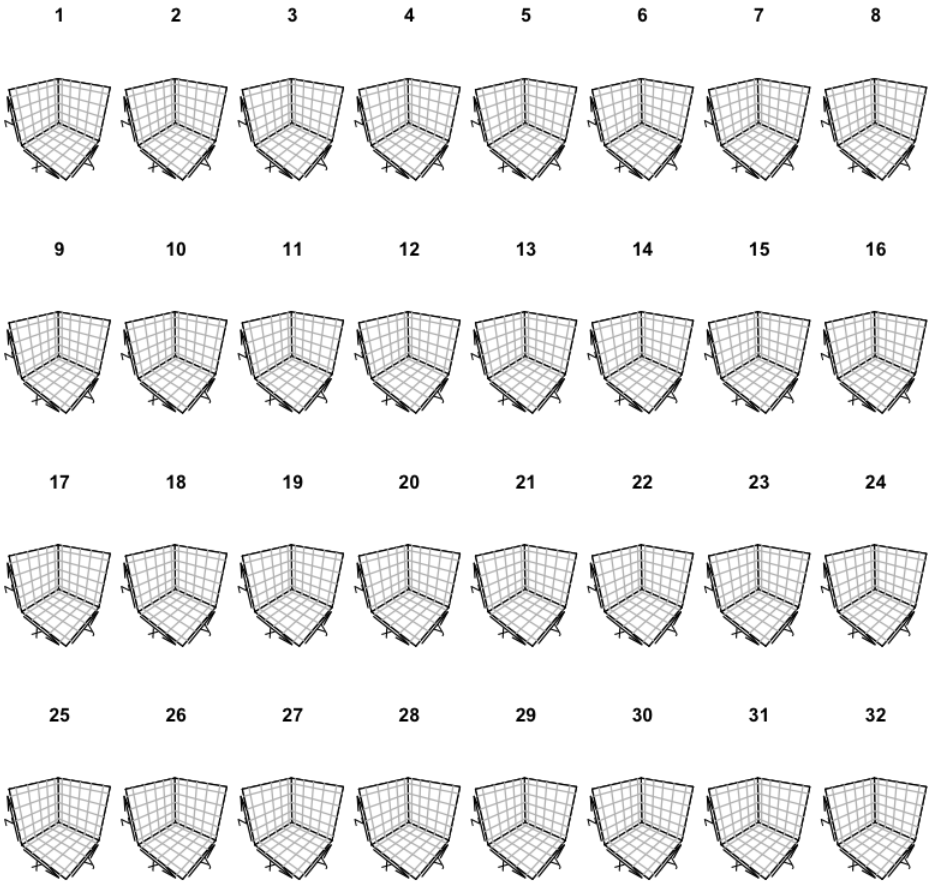

##### Agile12v2016 only

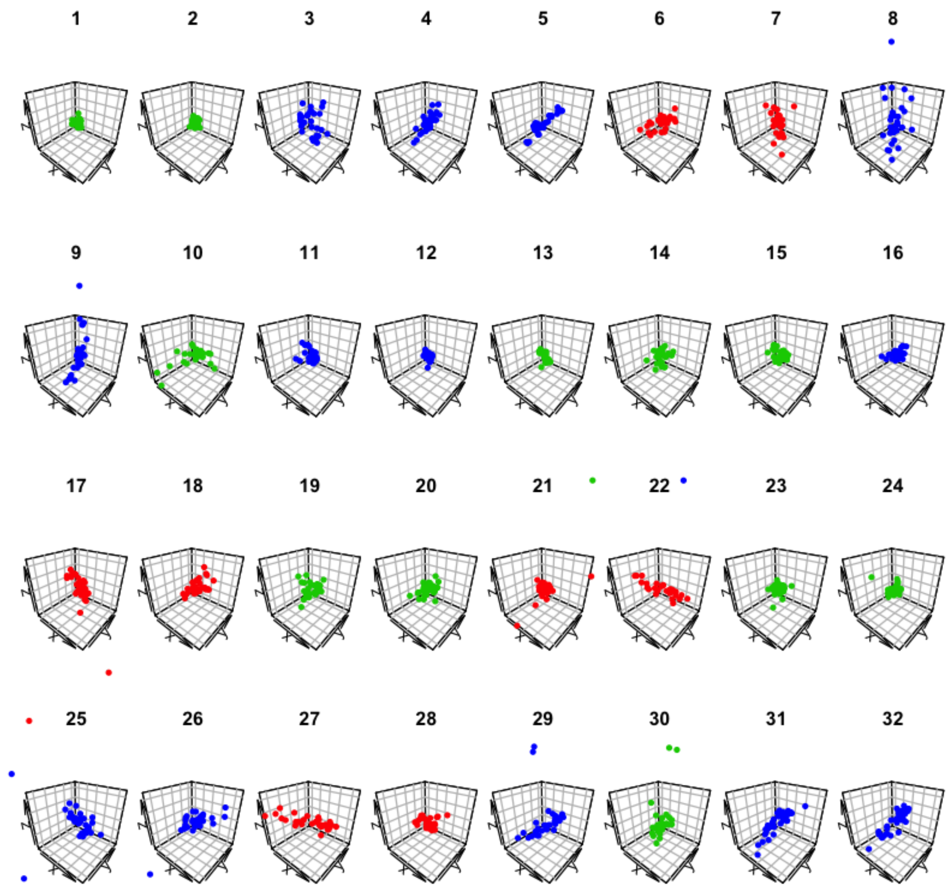

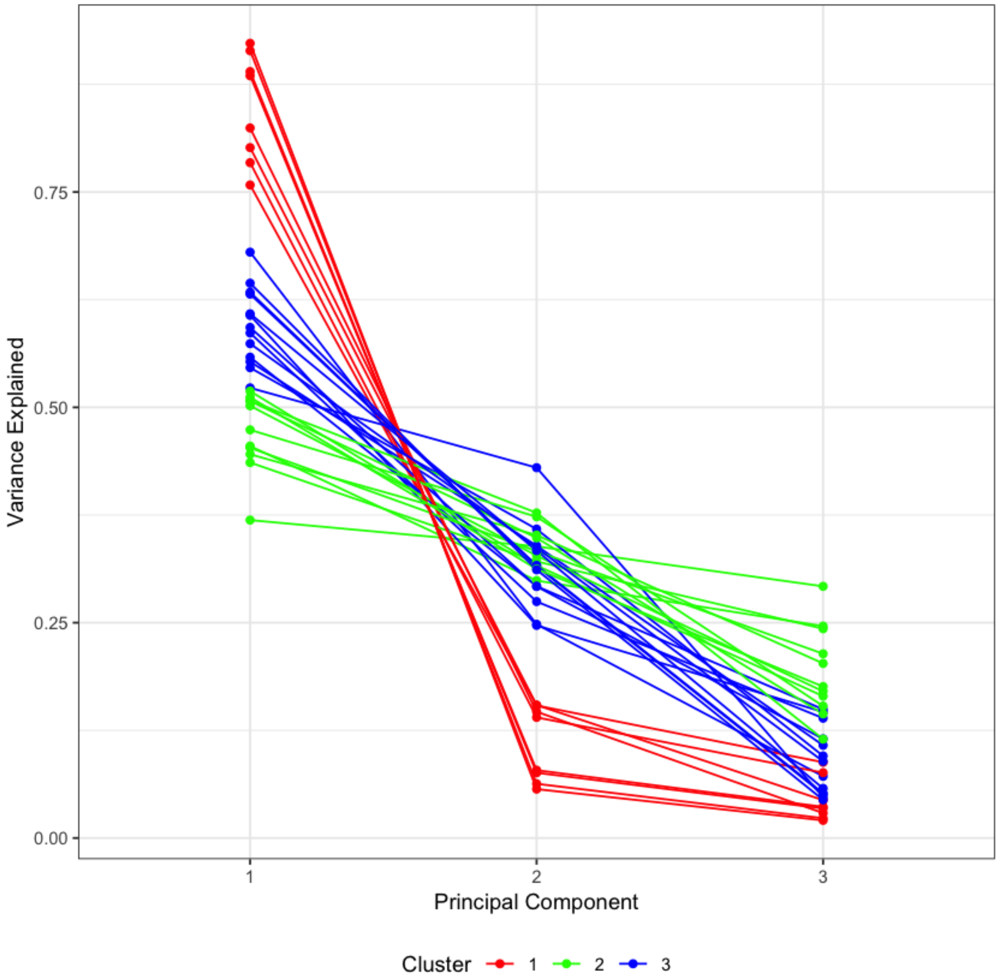

**Colin27 only**

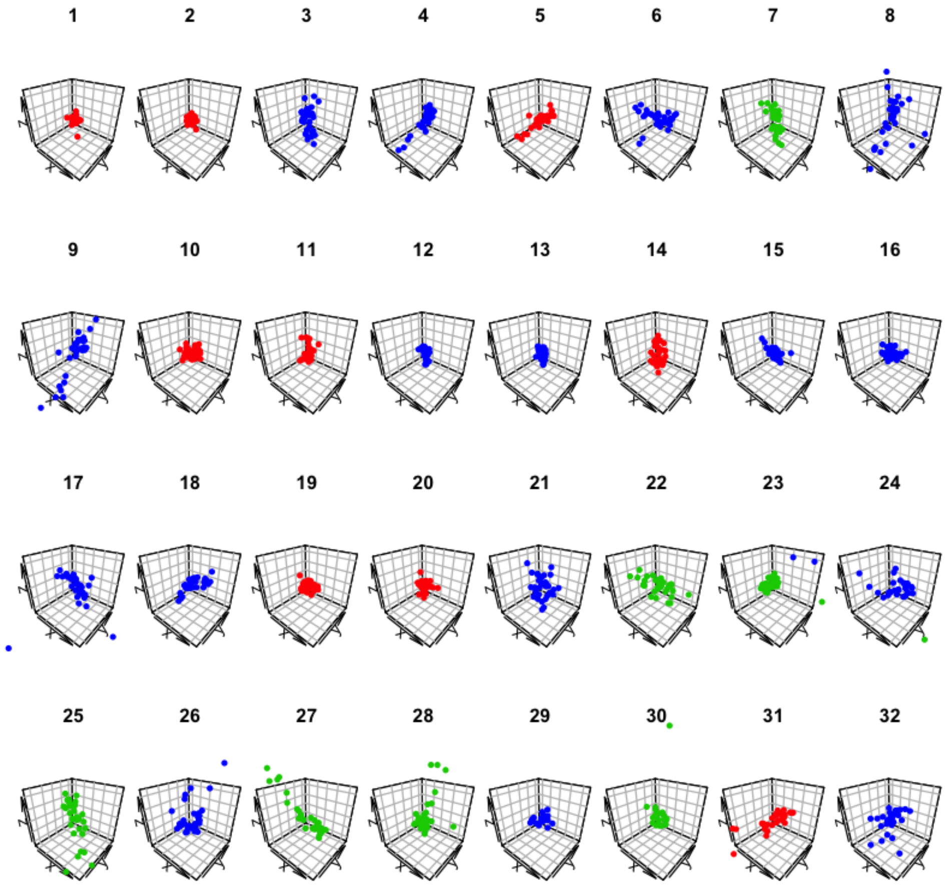

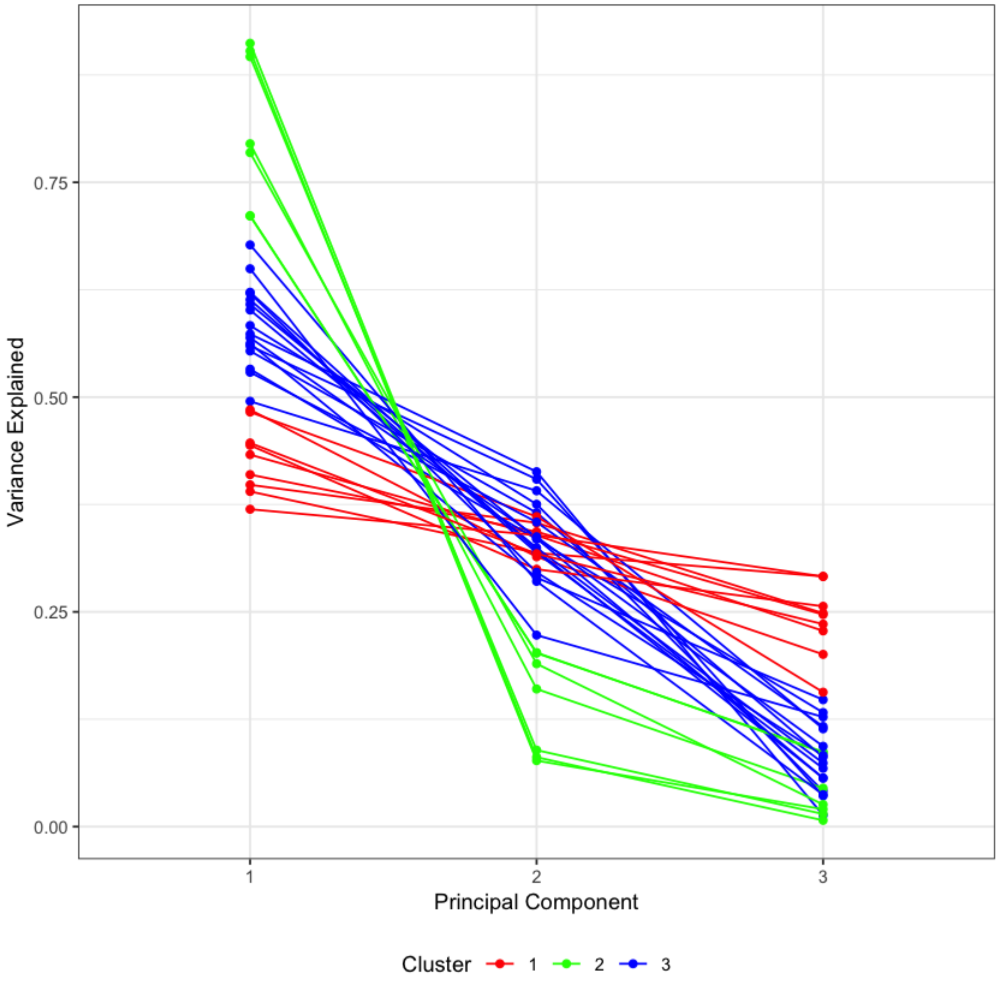

#### MNI152NLin2009bAsym Only

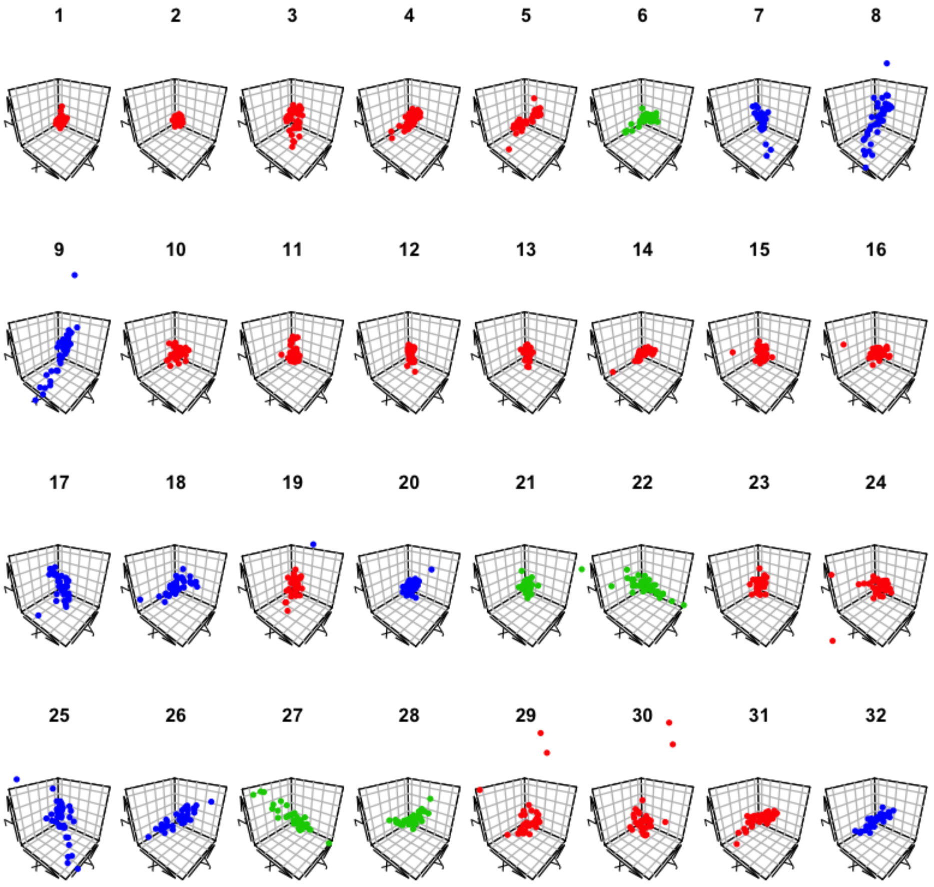

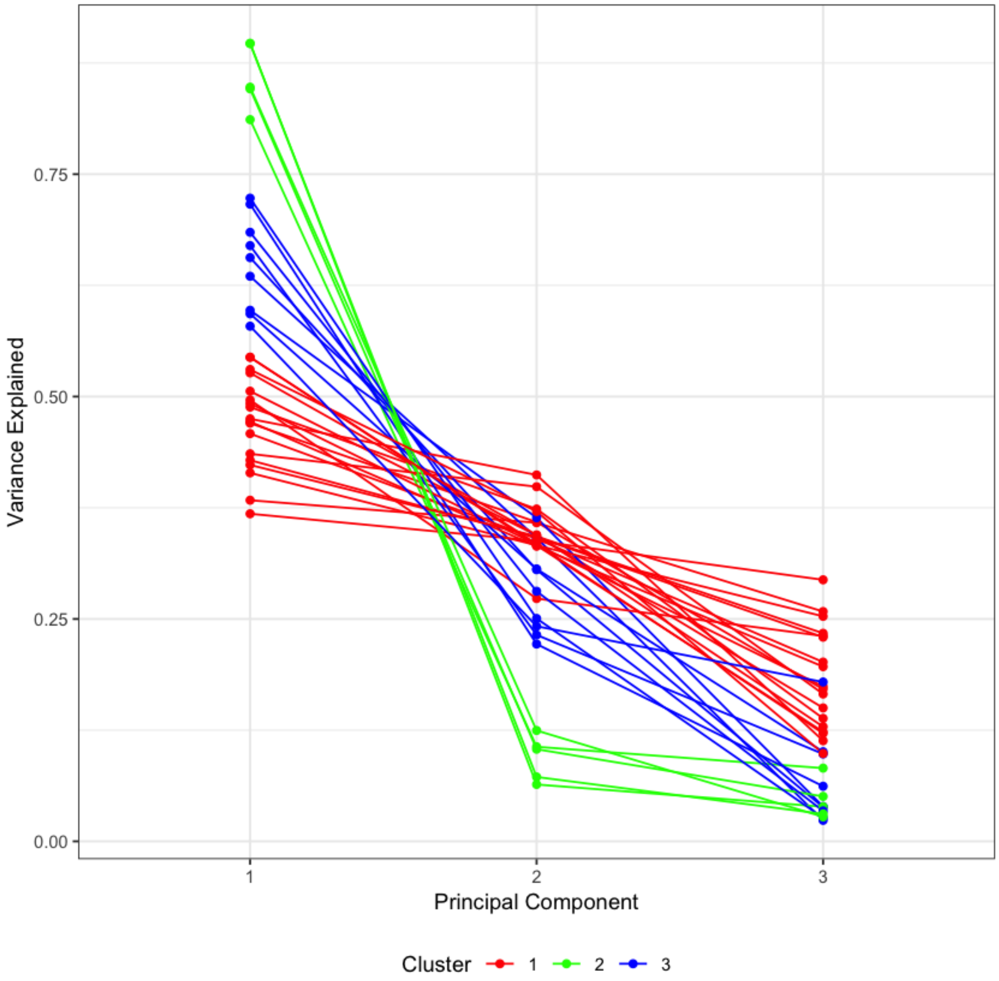

##### All templates combined

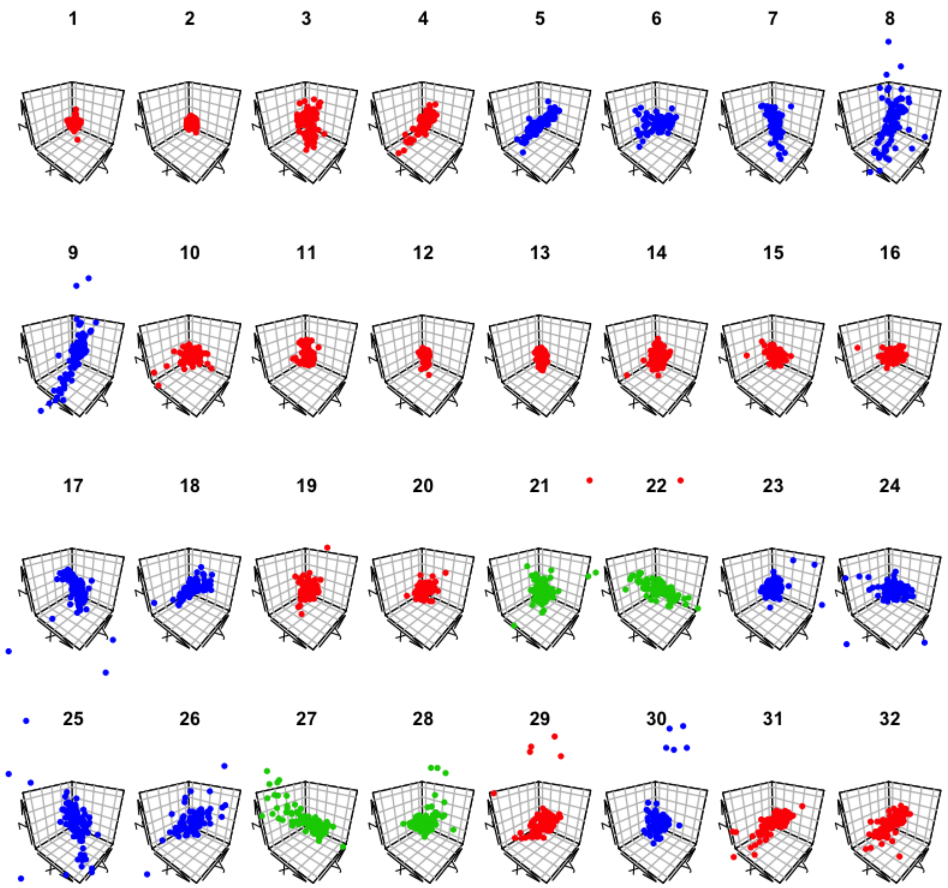

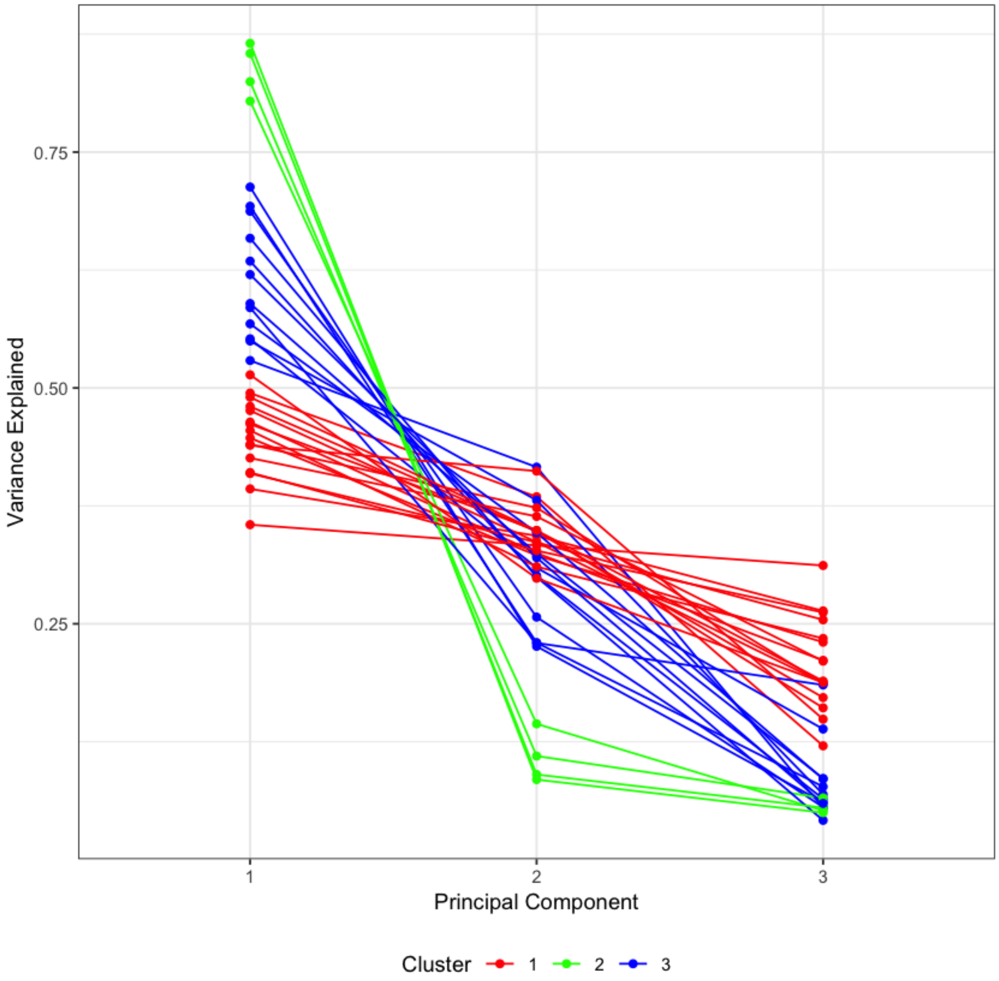

```
R version 3.5.1 (2018-07-02)
Platform: x86_64-apple-darwin13.4.0 (64-bit)
Running under: macOS  10.14.1

Matrix products: default
BLAS/LAPACK: /anaconda3/lib/R/lib/libRblas.dylib

locale:
[1] en_US.UTF-8/en_US.UTF-8/en_US.UTF-8/C/en_US.UTF-8/en_US.UTF-8

attached base packages:
[1] stats     graphics  grDevices utils     datasets  methods   base

other attached packages:
[1] plot3D_1.1.1   ggplot2_3.0.0  reshape2_1.4.3 digest_0.6.15  plyr_1.8.4

loaded via a namespace (and not attached):
 [1] Rcpp_0.12.18     compiler_3.5.1   pillar_1.3.0     bindr_0.1.1
 [5] base64enc_0.1-3  tools_3.5.1      uuid_0.1-2       jsonlite_1.5
 [9] evaluate_0.11    tibble_1.4.2     gtable_0.2.0     pkgconfig_2.0.1
[13] rlang_0.2.1      IRdisplay_0.5.0  IRkernel_0.8.12  bindrcpp_0.2.2
[17] repr_0.15.0      withr_2.1.2      stringr_1.3.1    dplyr_0.7.6
[21] grid_3.5.1       tidyselect_0.2.4 glue_1.3.0       R6_2.2.2
[25] pbdZMQ_0.3-3     purrr_0.2.5      magrittr_1.5     scales_0.5.0
[29] htmltools_0.3.6  misc3d_0.8-4     assertthat_0.2.0 colorspace_1.3-2
[33] labeling_0.3     stringi_1.2.4    lazyeval_0.2.1   munsell_0.5.0
[37] crayon_1.3.4
```
