## Supplementary material for "A framework for evaluating correspondence between brain images using anatomical fiducials": S2_PHASE2_subject_validation.pdf

### Phase 2: Protocol Validation for Individual Subjects

This notebook contains results validating the AFID32 protocol on individual subjects from the OASIS-1 databank.

#### OAS1 Subset: Demographics

Demographics here.

'Total: 58.0 +/- 17.9 years; Range: 25-91'

'Female: 17/30 (56.7%)'

'Total: 1.28 +/- 3.03 mm; Outliers: 28/2880 (0.97%)'

| | fid | subject | mri_session | name | description | mean_AFLE |
|---|---|---|---|---|---|---|
| **1373** | 29 | OAS1_0203 | MR1 | 29 | R ventral occipital horn | 16.19882 |
| **1501** | 29 | OAS1_0216 | MR1 | 29 | R ventral occipital horn | 17.77257 |
| **1502** | 30 | OAS1_0216 | MR1 | 30 | L ventral occipital horn | 11.61197 |
| **2043** | 27 | OAS1_0256 | MR1 | 28 | L indusium griseum origin | 15.80488 |
| **2044** | 28 | OAS1_0256 | MR1 | 27 | R indusium griseum origin | 15.35560 |
| **2141** | 29 | OAS1_0263 | MR1 | 29 | R ventral occipital horn | 39.35092 |
| **2142** | 30 | OAS1_0263 | MR1 | 30 | L ventral occipital horn | 40.64370 |
| **2173** | 29 | OAS1_0263 | MR1 | 29 | R ventral occipital horn | 78.74419 |
| **2174** | 30 | OAS1_0263 | MR1 | 30 | L ventral occipital horn | 80.42163 |
| **2205** | 29 | OAS1_0263 | MR1 | 29 | R ventral occipital horn | 39.39868 |
| **2206** | 30 | OAS1_0263 | MR1 | 30 | L ventral occipital horn | 39.79291 |
| **2235** | 27 | OAS1_0266 | MR1 | 27 | R indusium griseum origin | 23.44415 |
| **2236** | 28 | OAS1_0266 | MR1 | 28 | L indusium griseum origin | 24.30401 |
| **2267** | 27 | OAS1_0266 | MR1 | 27 | R indusium griseum origin | 10.56158 |
| **2268** | 28 | OAS1_0266 | MR1 | 28 | L indusium griseum origin | 12.04423 |
| **2299** | 27 | OAS1_0266 | MR1 | 27 | R indusium griseum origin | 12.98773 |
| **2300** | 28 | OAS1_0266 | MR1 | 28 | L indusium griseum origin | 12.35749 |
| **2534** | 6 | OAS1_0303 | MR1 | 6 | R superior LMS | 14.24872 |
| **2535** | 7 | OAS1_0303 | MR1 | 7 | L superior LMS | 13.98733 |
| **2653** | 29 | OAS1_0343 | MR1 | 29 | R ventral occipital horn | 15.83104 |
| **2942** | 30 | OAS1_0365 | MR1 | 30 | L ventral occipital horn | 10.92964 |
| **3387** | 27 | OAS1_0456 | MR1 | 27 | R indusium griseum origin | 23.38522 |
| **3388** | 28 | OAS1_0456 | MR1 | 28 | L indusium griseum origin | 23.76189 |
| **3390** | 30 | OAS1_0456 | MR1 | 30 | L ventral occipital horn | 17.74944 |
| **3419** | 27 | OAS1_0456 | MR1 | 27 | R induseum griseum origin | 10.64591 |
| **3420** | 28 | OAS1_0456 | MR1 | 28 | L induseum griseum origin | 10.43077 |
| **3451** | 27 | OAS1_0456 | MR1 | 27 | R indusium griseum origin | 12.88382 |
| **3452** | 28 | OAS1_0456 | MR1 | 28 | L indusium griseum origin | 13.53997 |

#### Individual Subject Results: Post-QC

Re-analysis after quality control and filtering of outliers.

'Total: 0.94 +/- 0.73 mm; Outliers: 0/2872 (0.00%)'

#### Inter-Rater AFLE

'Total: 1.58 +/- 1.02 mm'

| AFID | Description | Mean AFLE Pre-QC | Mean AFLE Post-QC | Inter-Rater AFLE Post-QC |
|---|---|---|---|---|
| 01 | AC | 0.36±0.21 (1.29) | 0.36±0.21 (1.29) | 0.60±0.25 (1.38) |
| 02 | PC | 0.34±0.16 (0.88) | 0.34±0.16 (0.88) | 0.57±0.21 (1.22) |
| 03 | infracollicular sulcus | 0.78±0.48 (3.07) | 0.78±0.48 (3.07) | 1.34±0.64 (3.84) |
| 04 | PMJ | 0.83±0.49 (2.44) | 0.83±0.49 (2.44) | 1.41±0.55 (2.55) |
| 05 | superior interpeduncular fossa | 1.20±0.75 (3.50) | 1.20±0.75 (3.50) | 2.04±0.90 (4.25) |
| 06 | R superior LMS | 1.30±1.74 (14.25) | 1.01±0.55 (2.85) | 1.70±0.68 (3.13) |
| 07 | L superior LMS | 1.36±1.71 (13.99) | 1.06±0.61 (3.45) | 1.72±0.71 (3.89) |
| 08 | R inferior LMS | 1.13±0.75 (5.13) | 1.03±0.57 (2.99) | 1.77±0.74 (3.43) |
| 09 | L inferior LMS | 1.10±0.80 (5.31) | 1.01±0.62 (2.72) | 1.71±0.86 (3.71) |
| 10 | culmen | 0.99±0.99 (5.66) | 0.83±0.62 (3.07) | 1.35±0.82 (3.42) |
| 11 | intermammillary sulcus | 0.60±0.31 (1.62) | 0.60±0.31 (1.62) | 1.02±0.41 (1.86) |
| 12 | R MB | 0.40±0.23 (1.11) | 0.40±0.23 (1.11) | 0.69±0.32 (1.52) |
| 13 | L MB | 0.36±0.20 (1.20) | 0.36±0.20 (1.20) | 0.62±0.29 (1.62) |
| 14 | pineal gland | 0.68±0.47 (1.98) | 0.68±0.47 (1.98) | 1.16±0.69 (2.63) |
| 15 | R LV at AC | 1.00±0.90 (5.28) | 0.91±0.72 (4.45) | 1.55±1.08 (5.86) |
| 16 | L LV at AC | 1.01±0.80 (4.53) | 0.94±0.70 (4.53) | 1.60±1.08 (5.47) |
| 17 | R LV at PC | 0.92±0.54 (3.42) | 0.92±0.54 (3.42) | 1.54±0.77 (3.84) |
| 18 | L LV at PC | 0.87±0.42 (2.20) | 0.87±0.42 (2.20) | 1.46±0.55 (2.80) |
| 19 | genu of CC | 0.97±0.81 (5.16) | 0.89±0.63 (3.69) | 1.50±0.89 (4.30) |
| 20 | splenium | 0.54±0.25 (1.24) | 0.54±0.25 (1.24) | 0.91±0.35 (1.66) |
| 21 | R AL temporal horn | 1.44±1.09 (7.01) | 1.30±0.86 (4.45) | 2.21±1.13 (5.92) |
| 22 | L AL temporal horn | 1.22±0.77 (4.11) | 1.22±0.77 (4.11) | 2.04±1.01 (4.47) |
| 23 | R superior AM temporal horn | 1.28±1.27 (8.22) | 1.12±0.88 (4.69) | 1.86±1.19 (4.97) |
| 24 | L superior AM temporal horn | 1.09±1.22 (7.54) | 0.83±0.61 (3.66) | 1.39±0.85 (4.60) |
| 25 | R inferior AM temporal horn | 1.69±1.43 (9.03) | 1.44±0.91 (4.72) | 2.39±1.23 (5.07) |
| 26 | L inferior AM temporal horn | 1.99±1.75 (8.79) | 1.49±1.09 (4.70) | 2.42±1.47 (6.64) |
| 27 | R indusium griseum origin | 3.13±4.19 (23.44) | 1.77±0.99 (4.77) | 2.95±1.20 (5.75) |
| 28 | L indusium griseum origin | 2.99±4.30 (24.30) | 1.68±1.00 (5.00) | 2.75±1.29 (5.78) |
| 29 | R ventral occipital horn | 3.64±10.36 (78.74) | 0.69±0.39 (2.11) | 1.14±0.54 (2.53) |
| 30 | L ventral occipital horn | 3.43±10.38 (80.42) | 0.86±0.67 (4.94) | 1.39±0.98 (5.72) |
| 31 | R olfactory sulcal fundus | 0.99±0.53 (2.29) | 0.99±0.53 (2.29) | 1.71±0.60 (2.84) |
| 32 | L olfactory sulcal fundus | 1.21±0.74 (4.53) | 1.21±0.74 (4.53) | 2.11±0.92 (5.81) |

### Secondary Analyses

We evaluated whether there was any evidence of an effect of demographics on AFLE.

| | | |
|---|---|---|
| **(Intercept)** | 0.7694 | 0e+00 |
| **age** | 0.0030 | 1e-04 |

#### Did AFLE worsen with the age of the subject for specific AFIDs?

We wanted to see if specific AFIDs tended to worsen with age of the OAS1 participant scan. Worsened for AFID17-18: bilateral LV at PC.

| fid | (Intercept) | pval_(Intercept) | age | pval_session | pval_session_adjusted | pval_session_significant |
|---|---|---|---|---|---|---|
| 1 | 0.19 | 0.0102 | 0.00 | 0.0133 | 0.1422 | FALSE |
| 2 | 0.24 | 0.0001 | 0.00 | 0.0964 | 0.3426 | FALSE |
| 3 | 0.93 | 0.0000 | 0.00 | 0.3885 | 0.5920 | FALSE |
| 4 | 0.86 | 0.0000 | 0.00 | 0.8868 | 0.9063 | FALSE |
| 5 | 0.81 | 0.0033 | 0.01 | 0.1364 | 0.4095 | FALSE |
| 6 | 1.24 | 0.0000 | 0.00 | 0.2292 | 0.4584 | FALSE |
| 7 | 0.66 | 0.0035 | 0.01 | 0.0572 | 0.2466 | FALSE |
| 8 | 0.79 | 0.0003 | 0.00 | 0.2276 | 0.4584 | FALSE |
| 9 | 0.60 | 0.0074 | 0.01 | 0.0557 | 0.2466 | FALSE |
| 10 | 0.61 | 0.0075 | 0.00 | 0.3133 | 0.5321 | FALSE |
| 11 | 0.67 | 0.0000 | 0.00 | 0.5306 | 0.7075 | FALSE |
| 12 | 0.52 | 0.0000 | 0.00 | 0.1408 | 0.4095 | FALSE |
| 13 | 0.42 | 0.0000 | 0.00 | 0.4399 | 0.6120 | FALSE |
| 14 | 0.73 | 0.0001 | 0.00 | 0.7578 | 0.8362 | FALSE |
| 15 | 0.82 | 0.0025 | 0.00 | 0.7391 | 0.8362 | FALSE |
| 16 | 0.88 | 0.0008 | 0.00 | 0.8194 | 0.8741 | FALSE |
| 17 | 0.18 | 0.3163 | 0.01 | 0.0000 | 0.0013 | TRUE |
| 18 | 0.44 | 0.0030 | 0.01 | 0.0029 | 0.0461 | TRUE |
| 19 | 0.92 | 0.0002 | 0.00 | 0.9063 | 0.9063 | FALSE |
| 20 | 0.44 | 0.0000 | 0.00 | 0.2772 | 0.5217 | FALSE |
| 21 | 0.92 | 0.0043 | 0.01 | 0.2049 | 0.4584 | FALSE |
| 22 | 1.14 | 0.0001 | 0.00 | 0.7487 | 0.8362 | FALSE |
| 23 | 0.99 | 0.0029 | 0.00 | 0.6622 | 0.8150 | FALSE |
| 24 | 0.62 | 0.0057 | 0.00 | 0.3294 | 0.5321 | FALSE |
| 25 | 1.27 | 0.0002 | 0.00 | 0.5861 | 0.7502 | FALSE |
| 26 | 1.86 | 0.0000 | -0.01 | 0.3325 | 0.5321 | FALSE |
| 27 | 1.13 | 0.0021 | 0.01 | 0.0616 | 0.2466 | FALSE |
| 28 | 1.25 | 0.0008 | 0.01 | 0.2183 | 0.4584 | FALSE |
| 29 | 0.41 | 0.0039 | 0.00 | 0.0404 | 0.2466 | FALSE |
| 30 | 0.36 | 0.1322 | 0.01 | 0.0345 | 0.2466 | FALSE |
| 31 | 0.84 | 0.0000 | 0.00 | 0.4201 | 0.6111 | FALSE |
| 32 | 0.84 | 0.0022 | 0.01 | 0.1563 | 0.4167 | FALSE |

```
R version 3.5.1 (2018-07-02)
Platform: x86_64-apple-darwin14.5.0 (64-bit)
Running under: macOS High Sierra 10.13.2

Matrix products: default
BLAS: /System/Library/Frameworks/Accelerate.framework/Versions/A/Frameworks/vecLib.framework/Versions/A/libBLAS.dylib
LAPACK: /System/Library/Frameworks/Accelerate.framework/Versions/A/Frameworks/vecLib.framework/Versions/A/libLAPACK.dylib

locale:
[1] en_CA.UTF-8/en_CA.UTF-8/en_CA.UTF-8/C/en_CA.UTF-8/en_CA.UTF-8

attached base packages:
[1] stats     graphics  grDevices utils     datasets  methods   base

other attached packages:
[1] plot3D_1.1.1   ggplot2_3.0.0  reshape2_1.4.3 digest_0.6.16  plyr_1.8.4

loaded via a namespace (and not attached):
 [1] Rcpp_0.12.17     compiler_3.5.1   pillar_1.3.0     bindr_0.1.1
 [5] base64enc_0.1-3  tools_3.5.1      uuid_0.1-2       jsonlite_1.5
 [9] evaluate_0.11    tibble_1.4.2     gtable_0.2.0     pkgconfig_2.0.2
[13] rlang_0.2.1      IRdisplay_0.5.0  IRkernel_0.8.12  bindrcpp_0.2.2
[17] repr_0.15.0      withr_2.1.2      stringr_1.3.1    dplyr_0.7.6
[21] grid_3.5.1       tidyselect_0.2.4 glue_1.3.0       R6_2.2.2
[25] pbdZMQ_0.3-3     purrr_0.2.5      magrittr_1.5     scales_1.0.0
[29] htmltools_0.3.6  misc3d_0.8-4     assertthat_0.2.0 colorspace_1.3-2
[33] stringi_1.2.4    lazyeval_0.2.1   munsell_0.5.0    crayon_1.3.4
```
