## Supplementary material for "A framework for evaluating correspondence between brain images using anatomical fiducials": S3_PHASE3_subject_to_template.pdf

### Phase 3: Subject-to-Template Evaluation

This notebook compares voxel overlap measures against AFID-based metrics for evaluating spatial correspondence. The OASIS-1 dataset from from PHASE2 was processed using the Ants-based T1-to-MNI (`MNI152NLin2009bAsym`) registration workflow built-in to fMRIPrep.

```
Attaching package: ‘dplyr’

The following objects are masked from ‘package:plyr’:

    arrange, count, desc, failwith, id, mutate, rename, summarise,
    summarize

The following objects are masked from ‘package:stats’:

    filter, lag

The following objects are masked from ‘package:base’:

    intersect, setdiff, setequal, union

Loading required package: magrittr

Attaching package: ‘ggpubr’

The following object is masked from ‘package:plyr’:

    mutate
```

#### ROI Overlap

Values for pallidum, striatum, and thalamus.

| roi | side | jaccard_lin | jaccard_nlin | jaccard_lin_vs_nlin | kappa_lin | kappa_nlin | kappa_lin_vs_nlin |
|---|---|---|---|---|---|---|---|
| pallidum | left | 0.54±0.13 | 0.80±0.03 | * | 0.69±0.11 | 0.89±0.02 | * |
| pallidum | right | 0.55±0.12 | 0.79±0.05 | * | 0.70±0.11 | 0.88±0.03 | * |
| striatum | left | 0.53±0.14 | 0.83±0.03 | * | 0.68±0.13 | 0.91±0.02 | * |
| striatum | right | 0.55±0.15 | 0.82±0.05 | * | 0.70±0.13 | 0.90±0.03 | * |
| thalamus | left | 0.70±0.11 | 0.86±0.03 | * | 0.82±0.08 | 0.93±0.02 | * |
| thalamus | right | 0.69±0.11 | 0.87±0.03 | * | 0.81±0.08 | 0.93±0.02 | * |

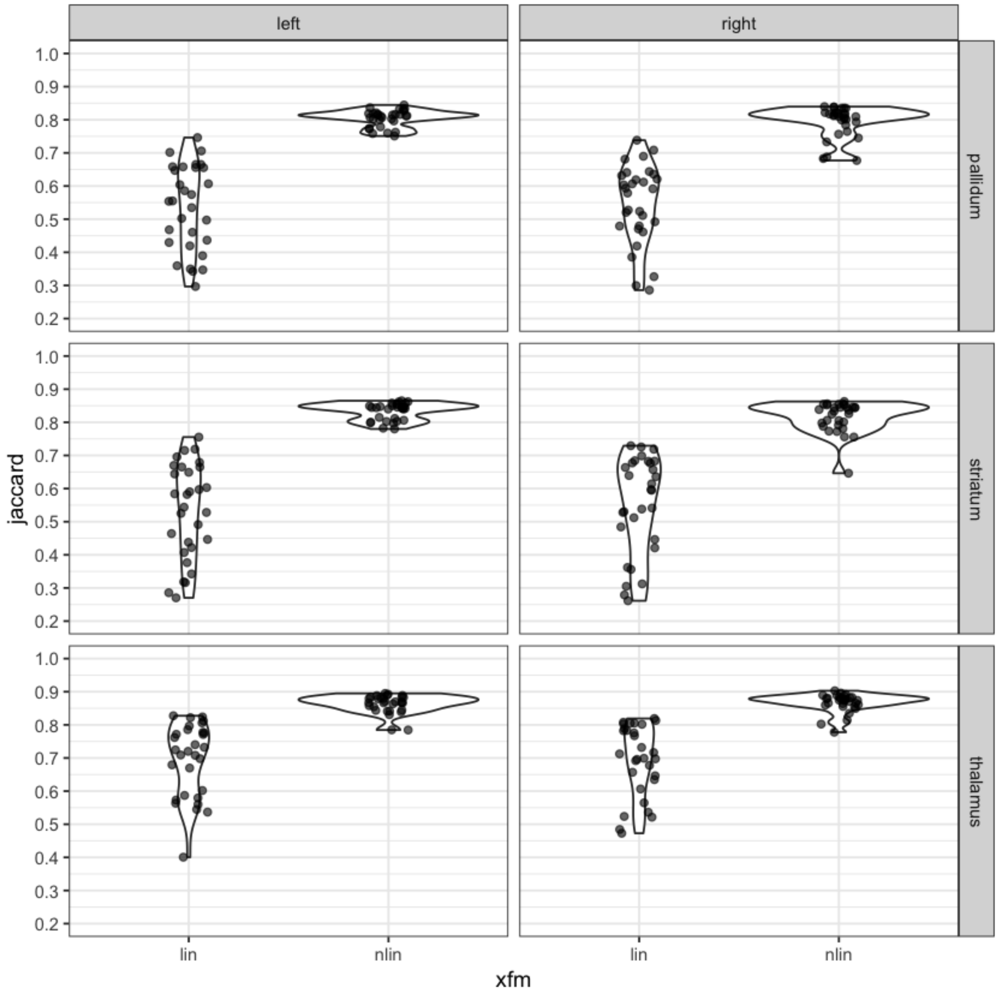
left
right
pallidum
striatum
thalamus
jaccard
1.0
0.9
0.8
0.7
0.6
0.5
0.4
0.3
0.2
lin
nlin
xfm

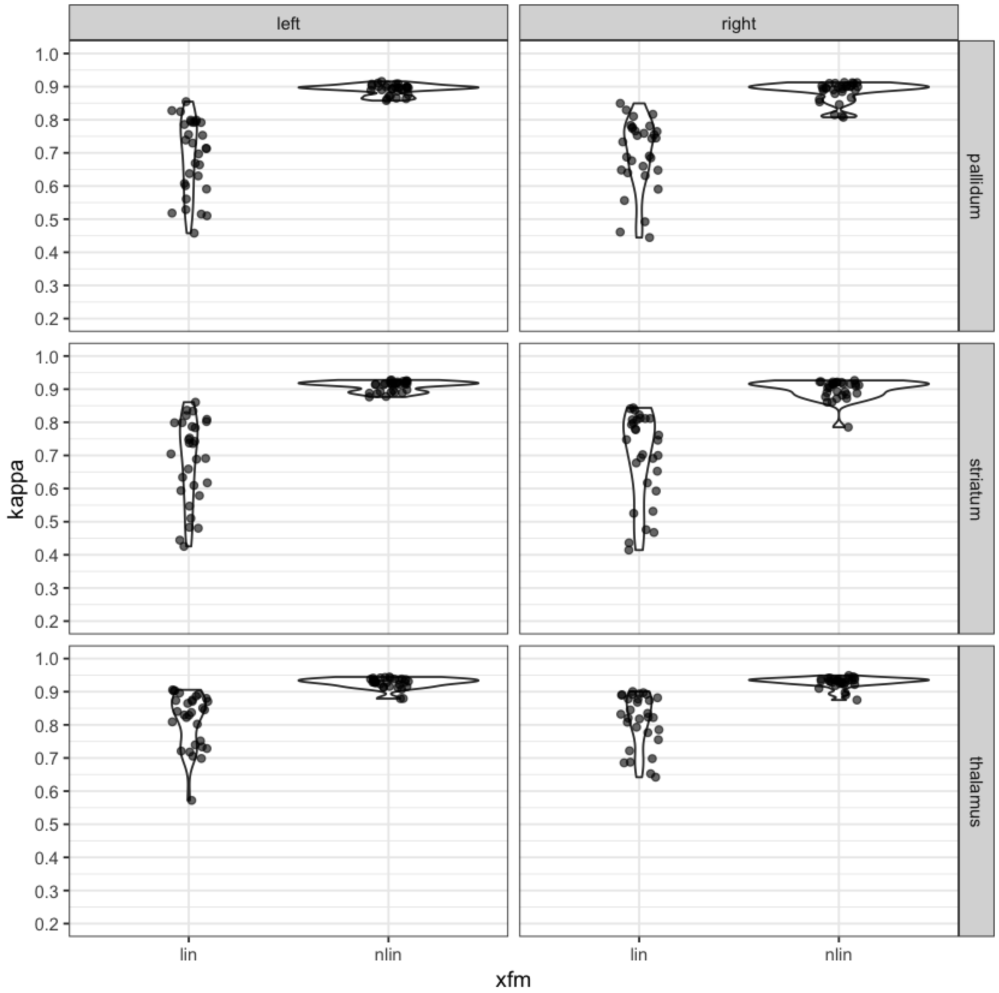

### AFRE

Anatomical FRE is evaluated here as a metric for looking at the spatial correspondence between images. Here they are summarized globally, for each AFID, and for each subject. Qualitatively, the ventricles are misaligned for OAS1_0109 which accounts for the maximally error observed in this analysis of > 30 mm AFRE.

#### Nonlinear Transform Results

'Total: 1.80 +/- 2.09 mm; Range: 0.07-32.78'

'Mean Max: 7.55 mm'

#### Linear Transform Results

'Total: 3.40 +/- 2.55 mm; Range: 0.28-36.26; Mean Max: '

'Mean Max: 10.25 mm'

```
        Wilcoxon rank sum test with continuity correction

data:  df_subjects_lin$AFRE and df_subjects_nlin$AFRE
W = 716930, p-value < 2.2e-16
alternative hypothesis: true location shift is not equal to 0
```

| AFID | Description | Mean AFRE lin | Mean AFRE nlin | lin vs nlin |
|---|---|---|---|---|
| 01 | AC | 2.15±0.97 (4.96) | 0.36±0.21 (0.99) | * |
| 02 | PC | 1.83±0.96 (4.58) | 0.57±0.29 (1.64) | * |
| 03 | infracollicular sulcus | 2.20±1.23 (5.71) | 0.93±0.53 (2.11) | * |
| 04 | PMJ | 2.50±1.36 (6.06) | 0.68±0.43 (2.13) | * |
| 05 | superior interpeduncular fossa | 2.35±1.06 (4.75) | 0.76±0.37 (1.69) | * |
| 06 | R superior LMS | 2.07±0.95 (4.32) | 1.17±0.74 (3.52) | * |
| 07 | L superior LMS | 2.03±0.85 (4.22) | 1.43±0.77 (2.88) | * |
| 08 | R inferior LMS | 2.45±1.37 (7.50) | 1.78±1.11 (5.41) | * |
| 09 | L inferior LMS | 2.54±1.26 (6.63) | 1.83±0.96 (3.99) | * |
| 10 | culmen | 4.50±2.93 (12.72) | 2.73±2.81 (10.12) | * |
| 11 | intermammillary sulcus | 2.81±1.62 (6.30) | 1.44±0.60 (2.73) | * |
| 12 | R MB | 2.72±1.67 (6.90) | 0.93±0.48 (1.90) | * |
| 13 | L MB | 2.84±1.70 (6.14) | 1.01±0.62 (2.93) | * |
| 14 | pineal gland | 2.53±1.39 (5.70) | 2.01±1.24 (6.16) | |
| 15 | R LV at AC | 4.44±1.84 (7.90) | 2.70±1.59 (7.85) | * |
| 16 | L LV at AC | 4.50±1.95 (8.40) | 2.11±1.72 (7.92) | * |
| 17 | R LV at PC | 4.81±2.54 (10.07) | 2.96±2.42 (9.46) | * |
| 18 | L LV at PC | 4.80±2.64 (10.34) | 3.01±2.22 (8.13) | * |
| 19 | genu of CC | 3.73±1.82 (7.88) | 1.56±0.76 (3.32) | * |
| 20 | splenium | 2.96±1.88 (7.57) | 0.97±0.60 (2.93) | * |
| 21 | R AL temporal horn | 3.79±1.71 (7.50) | 1.70±1.09 (5.23) | * |
| 22 | L AL temporal horn | 3.62±1.45 (6.98) | 1.67±0.98 (4.31) | * |
| 23 | R superior AM temporal horn | 3.34±1.63 (7.25) | 1.93±1.34 (6.85) | * |
| 24 | L superior AM temporal horn | 3.44±1.80 (8.20) | 1.67±1.25 (5.80) | * |
| 25 | R inferior AM temporal horn | 4.02±1.97 (8.32) | 2.41±1.16 (5.61) | * |
| 26 | L inferior AM temporal horn | 4.13±1.70 (8.20) | 2.21±1.09 (4.84) | * |
| 27 | R indusium griseum origin | 3.36±2.07 (8.46) | 2.06±1.49 (6.40) | * |
| 28 | L indusium griseum origin | 3.60±1.68 (8.83) | 2.05±1.37 (5.00) | * |
| 29 | R ventral occipital horn | 5.86±6.32 (36.26) | 3.44±5.77 (32.78) | * |
| 30 | L ventral occipital horn | 6.99±6.72 (33.74) | 4.51±6.28 (29.76) | * |
| 31 | R olfactory sulcal fundus | 2.83±1.36 (7.50) | 1.37±0.95 (3.44) | * |
| 32 | L olfactory sulcal fundus | 2.94±1.28 (6.49) | 1.57±0.84 (3.41) | * |

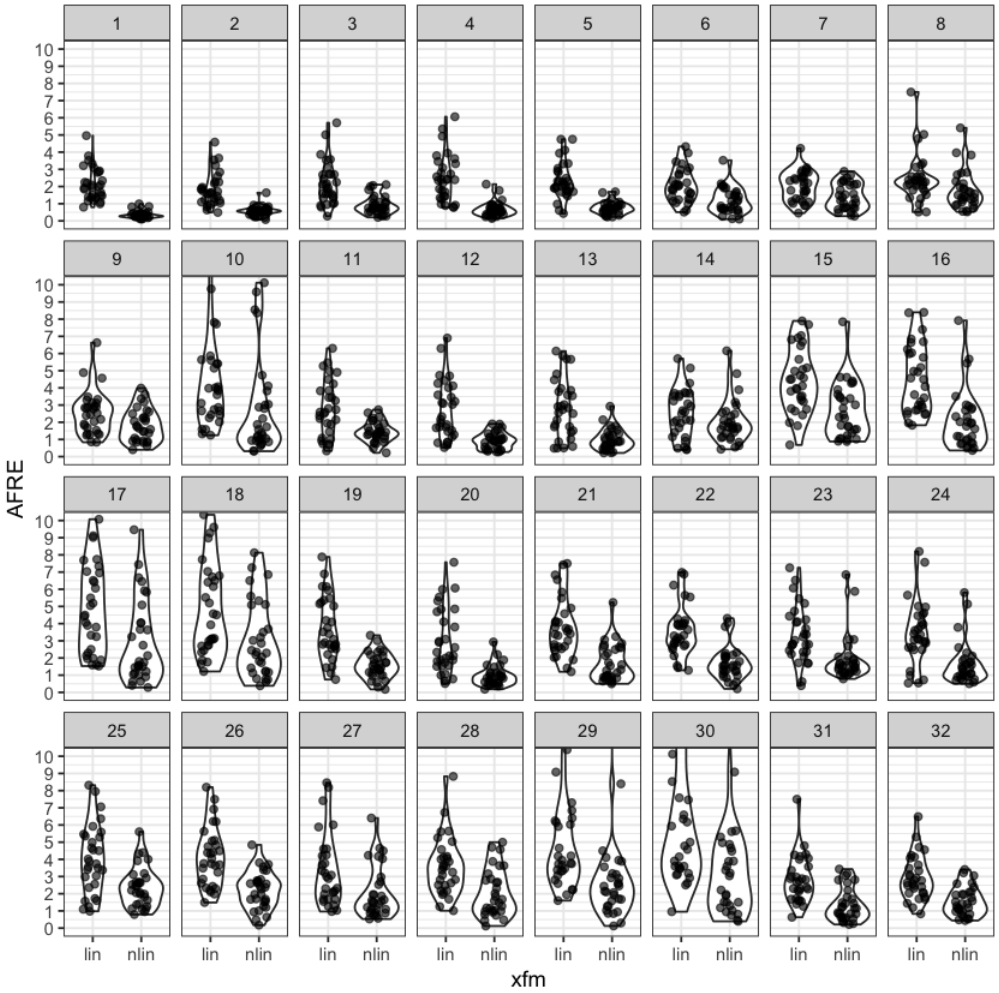

#### Subject level analysis of lin versus nlin

Revealed 3 subjects where mean AFRE was not statistically different. However, individual afids demonstrated high AFRE.

One subject appeared to be well registered with linear registration alone. The other two had extreme registration errors (over 8 mm AFRE).

| subject | AFRE_lin | AFRE_nlin | AFRE_qval_significant |
|---|---|---|---|
| OAS1_0010 | 4.44±2.06 (11.19) | 1.82±1.25 (5.61) | * |
| OAS1_0086 | 3.19±1.41 (6.40) | 1.33±1.14 (5.66) | * |
| OAS1_0101 | 3.10±2.22 (8.83) | 2.46±2.14 (8.36) | |
| OAS1_0109 | 4.86±8.12 (36.26) | 3.89±7.34 (32.78) | |
| OAS1_0114 | 3.31±2.03 (8.32) | 1.52±1.21 (5.88) | * |
| OAS1_0117 | 4.08±2.15 (9.76) | 1.74±1.81 (10.12) | * |
| OAS1_0145 | 2.33±1.69 (7.57) | 1.27±1.45 (6.85) | * |
| OAS1_0177 | 2.84±1.78 (7.06) | 1.54±0.82 (2.83) | * |
| OAS1_0180 | 4.08±2.20 (10.13) | 2.52±1.69 (6.45) | * |
| OAS1_0188 | 3.35±1.80 (9.08) | 1.65±1.21 (5.08) | * |
| OAS1_0200 | 2.56±1.45 (7.88) | 1.47±0.96 (4.84) | * |
| OAS1_0203 | 3.78±3.89 (23.96) | 2.46±3.89 (22.40) | * |
| OAS1_0216 | 2.19±1.61 (7.58) | 1.73±1.02 (4.48) | |
| OAS1_0239 | 2.89±2.03 (11.68) | 1.56±1.63 (8.55) | * |
| OAS1_0249 | 3.34±1.65 (8.53) | 1.63±1.08 (4.66) | * |
| OAS1_0255 | 3.48±1.77 (6.68) | 1.29±0.75 (3.16) | * |
| OAS1_0256 | 4.16±2.00 (7.90) | 1.52±0.97 (3.56) | * |
| OAS1_0263 | 3.97±2.36 (10.34) | 1.29±0.96 (3.96) | * |
| OAS1_0266 | 3.67±1.18 (7.50) | 1.60±1.22 (6.16) | * |
| OAS1_0274 | 2.90±1.87 (7.73) | 1.99±2.34 (8.13) | * |
| OAS1_0284 | 3.90±2.59 (13.41) | 1.84±1.85 (8.39) | * |
| OAS1_0303 | 2.70±1.23 (5.41) | 1.47±0.85 (4.03) | * |
| OAS1_0343 | 3.32±1.69 (7.95) | 2.43±1.95 (9.46) | * |
| OAS1_0345 | 2.31±1.30 (6.16) | 1.57±1.10 (4.11) | * |
| OAS1_0357 | 2.58±1.61 (7.43) | 1.47±1.35 (5.30) | * |
| OAS1_0365 | 4.18±2.07 (9.61) | 1.65±1.35 (6.50) | * |
| OAS1_0371 | 2.64±1.31 (6.92) | 1.33±0.82 (3.81) | * |
| OAS1_0395 | 3.27±1.96 (11.14) | 1.68±1.28 (6.68) | * |
| OAS1_0398 | 3.85±3.04 (12.40) | 1.81±1.80 (9.09) | * |
| OAS1_0456 | 4.64±2.73 (12.72) | 2.42±2.01 (9.59) | * |

#### Finding Outlier Misregistrations

Here we identify all AFID points across subjects above some minimum threshold of AFRE error (arbitrarily set to 5 mm for notebook presentation purposes).

| | fid | subject | name | description | AFRE |
|---|---|---|---|---|---|
| **212** | 8 | OAS1_0180 | 8 | R inferior LMS | 5.411179 |
| **271** | 10 | OAS1_0456 | 10 | culmen | 9.585257 |
| **277** | 10 | OAS1_0239 | 10 | culmen | 8.550554 |
| **278** | 10 | OAS1_0117 | 10 | culmen | 10.118020 |
| **289** | 10 | OAS1_0101 | 10 | culmen | 8.359801 |
| **402** | 14 | OAS1_0266 | 14 | pineal gland | 6.158929 |
| **435** | 15 | OAS1_0274 | 15 | R LV at AC | 7.848187 |
| **451** | 16 | OAS1_0456 | 16 | L LV at AC | 5.677177 |
| **463** | 16 | OAS1_0274 | 16 | L LV at AC | 7.922394 |
| **471** | 16 | OAS1_0101 | 16 | L LV at AC | 5.436124 |
| **485** | 17 | OAS1_0101 | 17 | R LV at PC | 5.918516 |
| **486** | 17 | OAS1_0180 | 17 | R LV at PC | 6.450083 |
| **490** | 17 | OAS1_0395 | 17 | R LV at PC | 6.676601 |
| **491** | 17 | OAS1_0274 | 17 | R LV at PC | 7.440260 |
| **505** | 17 | OAS1_0188 | 17 | R LV at PC | 5.084890 |
| **506** | 17 | OAS1_0343 | 17 | R LV at PC | 9.463697 |
| **507** | 17 | OAS1_0284 | 17 | R LV at PC | 5.832009 |
| **518** | 18 | OAS1_0274 | 18 | L LV at PC | 8.125230 |
| **520** | 18 | OAS1_0343 | 18 | L LV at PC | 6.852881 |
| **527** | 18 | OAS1_0456 | 18 | L LV at PC | 5.597730 |
| **528** | 18 | OAS1_0180 | 18 | L LV at PC | 5.111037 |
| **534** | 18 | OAS1_0101 | 18 | L LV at PC | 7.257325 |
| **535** | 18 | OAS1_0365 | 18 | L LV at PC | 6.501676 |
| **537** | 18 | OAS1_0284 | 18 | L LV at PC | 5.089356 |
| **538** | 18 | OAS1_0010 | 18 | L LV at PC | 5.331059 |
| **621** | 21 | OAS1_0180 | 21 | R AL temporal horn | 5.232758 |
| **667** | 23 | OAS1_0145 | 23 | R superior AM temporal horn | 6.847816 |
| **674** | 23 | OAS1_0114 | 23 | R superior AM temporal horn | 5.875959 |
| **718** | 24 | OAS1_0398 | 24 | L superior AM temporal horn | 5.141291 |
| **719** | 24 | OAS1_0145 | 24 | L superior AM temporal horn | 5.803107 |
| **737** | 25 | OAS1_0180 | 25 | R inferior AM temporal horn | 5.605086 |
| **782** | 27 | OAS1_0109 | 27 | R indusium griseum origin | 6.403877 |
| **851** | 29 | OAS1_0109 | 29 | R ventral occipital horn | 32.777012 |
| **866** | 29 | OAS1_0284 | 29 | R ventral occipital horn | 8.394882 |
| **871** | 30 | OAS1_0109 | 30 | L ventral occipital horn | 29.762330 |
| **872** | 30 | OAS1_0203 | 30 | L ventral occipital horn | 22.404038 |
| **877** | 30 | OAS1_0010 | 30 | L ventral occipital horn | 5.611475 |
| **879** | 30 | OAS1_0357 | 30 | L ventral occipital horn | 5.304235 |
| **886** | 30 | OAS1_0398 | 30 | L ventral occipital horn | 9.086490 |
| **896** | 30 | OAS1_0086 | 30 | L ventral occipital horn | 5.660475 |

OAS1_0180 OAS1_0456 OAS1_0239 OAS1_0117 OAS1_0101 OAS1_0266 OAS1_0274 OAS1_0395 OAS1_0188 OAS1_0343 OAS1_0284
OAS1_0365 OAS1_0010 OAS1_0145 OAS1_0114 OAS1_0398 OAS1_0109 OAS1_0203 OAS1_0357 OAS1_0086

▸ **Levels**:

#### Investigation of individual OASIS-1 Subject AFIDs (OAS1_0180)

| | fid | subject | name | description | AFRE | xfm |
|---|---|---|---|---|---|---|
| **15** | 1 | OAS1_0180 | 1 | AC | 0.3333971 | nlin |
| **33** | 2 | OAS1_0180 | 2 | PC | 0.6045653 | nlin |
| **62** | 3 | OAS1_0180 | 3 | infracollicular sulcus | 0.8774145 | nlin |
| **99** | 4 | OAS1_0180 | 4 | PMJ | 0.9738354 | nlin |
| **125** | 5 | OAS1_0180 | 5 | superior interpeduncular fossa | 1.6461190 | nlin |
| **177** | 6 | OAS1_0180 | 6 | R superior LMS | 0.6837562 | nlin |
| **190** | 7 | OAS1_0180 | 7 | L superior LMS | 0.4821263 | nlin |
| **212** | 8 | OAS1_0180 | 8 | R inferior LMS | 5.4111786 | nlin |
| **246** | 9 | OAS1_0180 | 9 | L inferior LMS | 3.1787138 | nlin |
| **293** | 10 | OAS1_0180 | 10 | culmen | 3.8186248 | nlin |
| **315** | 11 | OAS1_0180 | 11 | intermammillary sulcus | 2.3186715 | nlin |
| **335** | 12 | OAS1_0180 | 12 | R MB | 1.7114213 | nlin |
| **365** | 13 | OAS1_0180 | 13 | L MB | 2.9269330 | nlin |
| **403** | 14 | OAS1_0180 | 14 | pineal gland | 2.6319054 | nlin |
| **432** | 15 | OAS1_0180 | 15 | R LV at AC | 1.7596951 | nlin |
| **459** | 16 | OAS1_0180 | 16 | L LV at AC | 1.0219584 | nlin |
| **486** | 17 | OAS1_0180 | 17 | R LV at PC | 6.4500826 | nlin |
| **528** | 18 | OAS1_0180 | 18 | L LV at PC | 5.1110374 | nlin |
| **557** | 19 | OAS1_0180 | 19 | genu of CC | 1.4385252 | nlin |
| **584** | 20 | OAS1_0180 | 20 | splenium | 0.9289616 | nlin |
| **621** | 21 | OAS1_0180 | 21 | R AL temporal horn | 5.2327578 | nlin |
| **641** | 22 | OAS1_0180 | 22 | L AL temporal horn | 2.0260472 | nlin |
| **663** | 23 | OAS1_0180 | 23 | R superior AM temporal horn | 1.4644496 | nlin |
| **709** | 24 | OAS1_0180 | 24 | L superior AM temporal horn | 1.8499234 | nlin |
| **737** | 25 | OAS1_0180 | 25 | R inferior AM temporal horn | 5.6050862 | nlin |
| **766** | 26 | OAS1_0180 | 26 | L inferior AM temporal horn | 2.7429303 | nlin |
| **783** | 27 | OAS1_0180 | 27 | R indusium griseum origin | 1.3961954 | nlin |
| **821** | 28 | OAS1_0180 | 28 | L indusium griseum origin | 2.6464879 | nlin |
| **863** | 29 | OAS1_0180 | 29 | R ventral occipital horn | 3.1210472 | nlin |
| **891** | 30 | OAS1_0180 | 30 | L ventral occipital horn | 4.8718270 | nlin |
| **905** | 31 | OAS1_0180 | 31 | R olfactory sulcal fundus | 2.2000853 | nlin |
| **933** | 32 | OAS1_0180 | 32 | L olfactory sulcal fundus | 3.2115890 | nlin |

#### Comparison of AFRE and Voxel Overlap

Evidence for focal misregistrations not captured using voxel overlap measures alone.

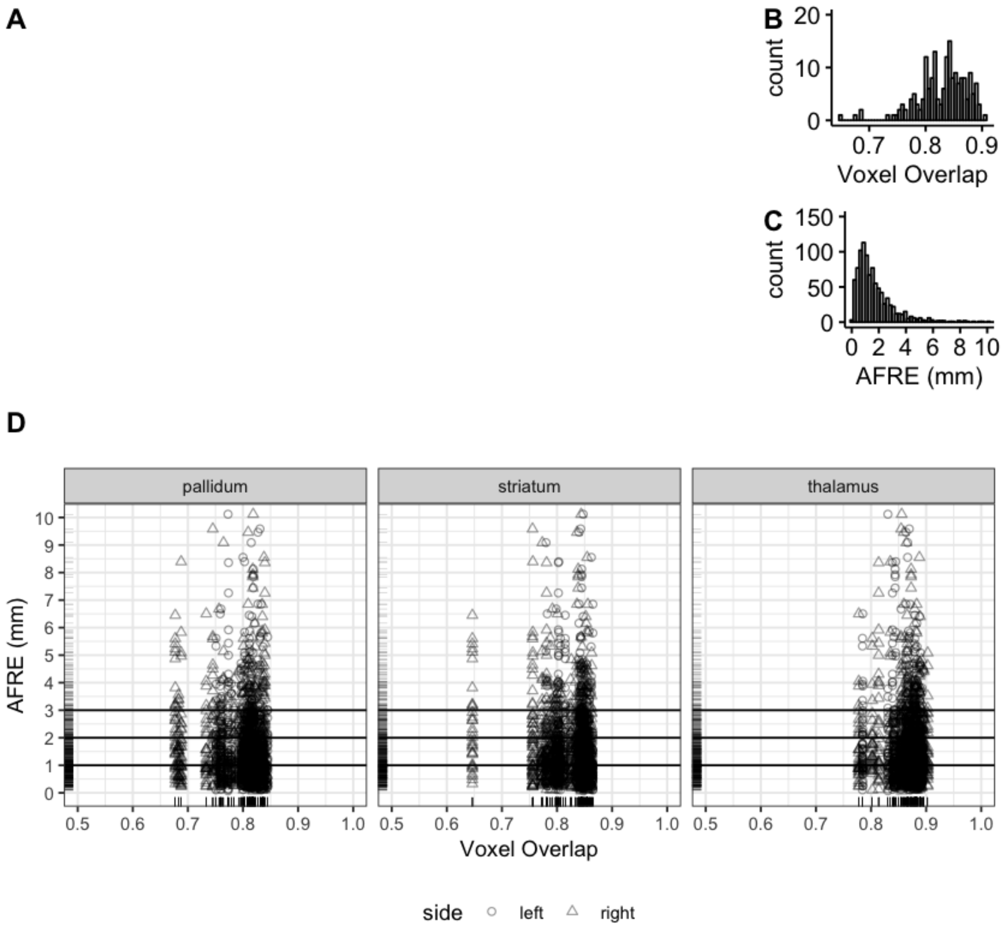

#### Correlation between AFRE and voxel overlap

A weak negative correlation was found between AFRE and standard voxel overlap measures at the global dataset level and for each specific ROI in isolation.

```
        Kendall's rank correlation tau

data:  compare_overlap_AFRE_nlin$jaccard and compare_overlap_AFRE_nlin$AFRE
z = -2.1686, p-value = 0.03011
alternative hypothesis: true tau is not equal to 0
sample estimates:
        tau
-0.01911451

        Kendall's rank correlation tau

data:  compare_overlap_AFRE_nlin$kappa and compare_overlap_AFRE_nlin$AFRE
z = -2.1686, p-value = 0.03011
alternative hypothesis: true tau is not equal to 0
sample estimates:
        tau
-0.01911451

        Kendall's rank correlation tau

data:  compare_overlap_AFRE_nlin$jaccard and compare_overlap_AFRE_nlin$kappa
z = 113.2, p-value < 2.2e-16
alternative hypothesis: true tau is not equal to 0
sample estimates:
tau
  1
```

| roi | side | AFRE_jaccard | AFRE_jaccard_pval | AFRE_kappa | AFRE_kappa_pval | AFRE_jaccard_pval_adjusted | AFRE_jaccard_pval_significant | AFRE_kappa_pv |
|---|---|---|---|---|---|---|---|---|
| pallidum | left | -0.021863046 | 0.31813186 | -0.021863046 | 0.31813186 | 0.3817582 | FALSE | 0.3817582 |
| pallidum | right | -0.045837311 | 0.03634823 | -0.045837311 | 0.03634823 | 0.1090447 | FALSE | 0.1090447 |
| striatum | left | -0.034786535 | 0.11219299 | -0.034786535 | 0.11219299 | 0.2243860 | FALSE | 0.2243860 |
| striatum | right | -0.049644573 | 0.02339900 | -0.049644573 | 0.02339900 | 0.1090447 | FALSE | 0.1090447 |
| thalamus | left | 0.006607498 | 0.76287336 | 0.006607498 | 0.76287336 | 0.7628734 | FALSE | 0.7628734 |
| thalamus | right | -0.029857412 | 0.17277515 | -0.029857412 | 0.17277515 | 0.2591627 | FALSE | 0.2591627 |

#### For each AFID and ROI

No correlation between voxel overlap measures and individual AFID AFREs were identified. However, plotting of voxel overlap against individual AFREs demonstrate the added sensitivity to misregistration when looking at individual AFID plots along the y-axis.

'Number of significant correlations (individual AFIDs vs voxel overlap): 0/192 (0.0%)'

##### Pallidum

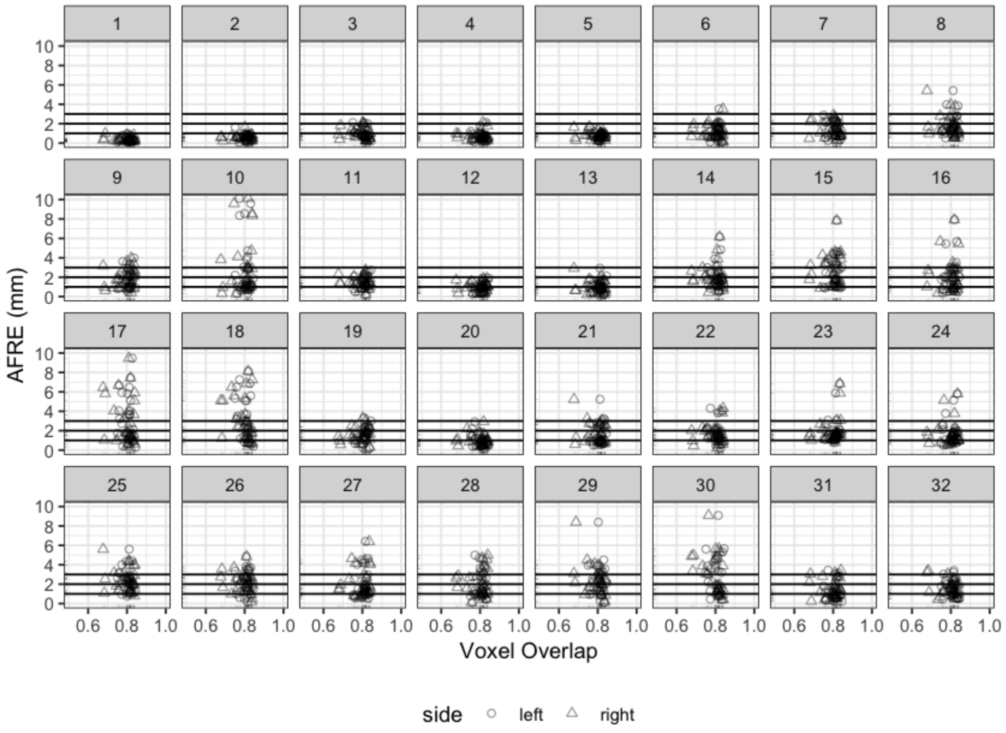
AFRE (mm)
Voxel Overlap
0.6 0.8 1.0
0 2 4 6 8 10
side
left
right

**Striatum**

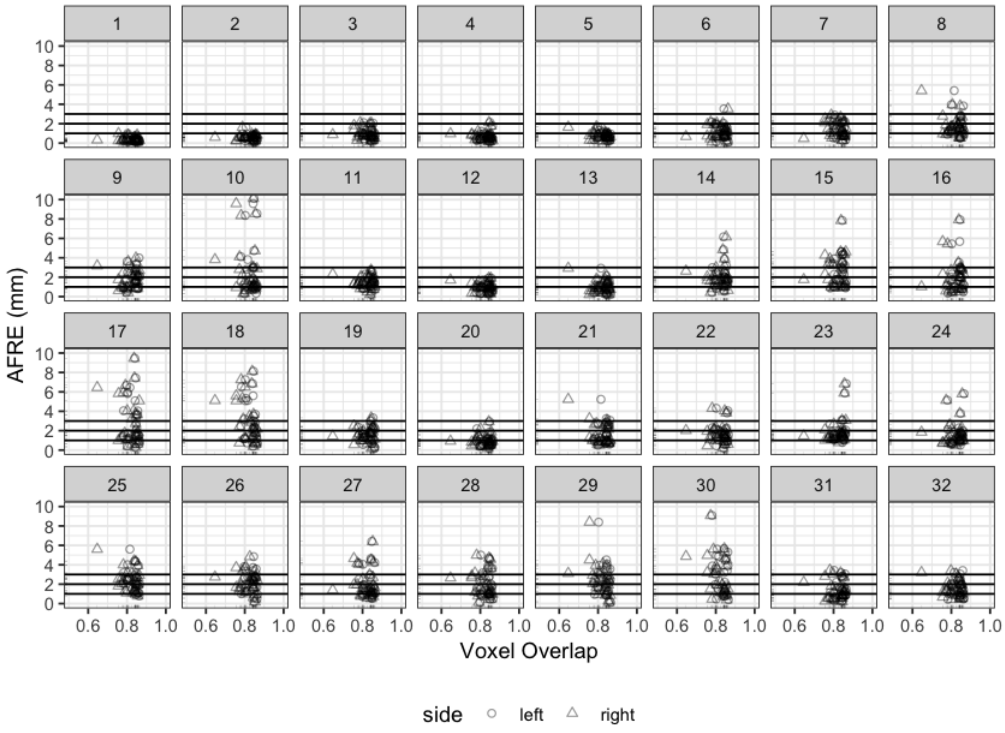

AFRE (mm)
0
2
4
6
8
10
0.6
0.8
1.0
Voxel Overlap
side
left
right

**Thalamus**

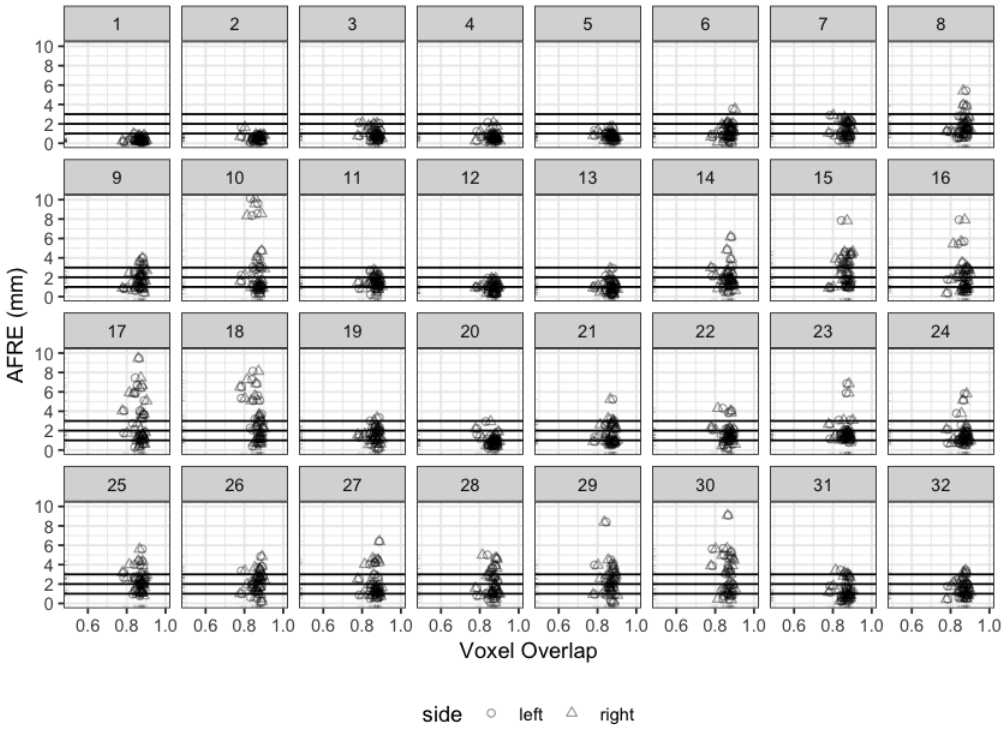

#### AFLE versus AFRE

In this section, we examine whether AFLE and AFRE are correlated and establish a baseline for when AFRE should be considered beyond placement-related (AFLE) error. The first vertical line is the mean AFLE for OASIS-1 subjects. Second is 1 s.d., third is 2 s.d.

A positive correlation between AFLE and AFRE was found to be statistically significant although the actual effect size of the correlation was small.

'Num Outlier AFIDs (> 2 s.d. above mean AFLE): 135/960 (14.06%)'

'Num Unique Outlier AFIDs (> 2 s.d. above mean AFLE): 22/32 (68.75%)'

```
        Kendall's rank correlation tau

data:  AFLE_vs_AFRE$AFRE and AFLE_vs_AFRE$mean_AFLE
z = 6.9796, p-value = 2.959e-12
alternative hypothesis: true tau is not equal to 0
sample estimates:
      tau
0.1504519
```

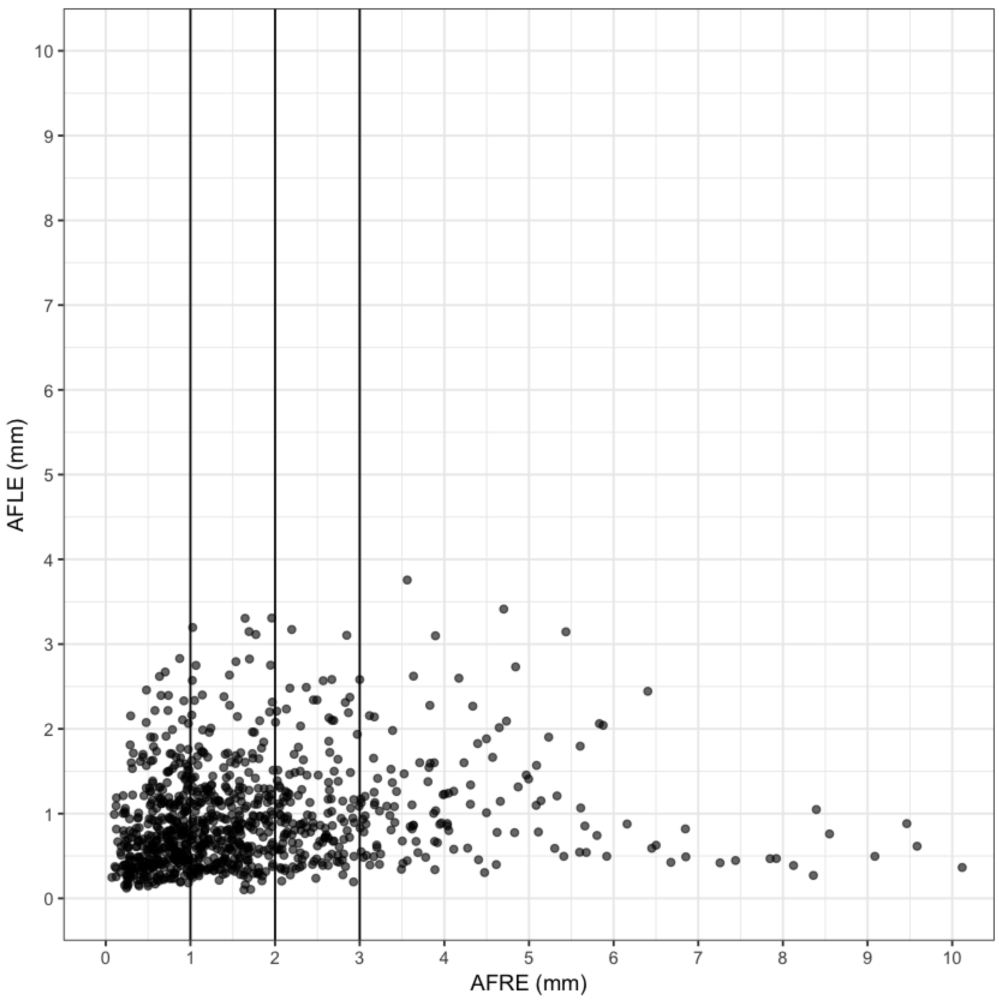

#### Secondary Analyses

We evaluated whether there was any evidence of an effect of demographics (e.g. age of the participants being registered) on AFRE. Age resulted in a global AFRE change of 0.0075 mm/year (i.e. a small but statistically significant effect). No specific AFIDs were found to contribute to this age-related AFRE change after multiple comparisons correction.

| | | |
|---|---|---|
| **(Intercept)** | 2.1635 | 0.0000 |
| **age** | 0.0075 | 0.0191 |

| fid | (Intercept) | pval_(Intercept) | age | pval_age | pval_age_adjusted | pval_age_significant |
|---|---|---|---|---|---|---|
| 1 | 1.15 | 0.03 | 0.00 | 0.83 | 0.86 | FALSE |
| 2 | 1.22 | 0.01 | 0.00 | 0.96 | 0.96 | FALSE |
| 3 | 1.96 | 0.00 | -0.01 | 0.42 | 0.75 | FALSE |
| 4 | 0.72 | 0.23 | 0.01 | 0.13 | 0.55 | FALSE |
| 5 | 1.77 | 0.00 | 0.00 | 0.66 | 0.83 | FALSE |
| 6 | 1.74 | 0.00 | 0.00 | 0.78 | 0.83 | FALSE |
| 7 | 1.03 | 0.01 | 0.01 | 0.05 | 0.43 | FALSE |
| 8 | 0.62 | 0.25 | 0.03 | 0.01 | 0.16 | FALSE |
| 9 | 1.54 | 0.00 | 0.01 | 0.20 | 0.55 | FALSE |
| 10 | 6.53 | 0.00 | -0.05 | 0.02 | 0.32 | FALSE |
| 11 | 1.41 | 0.03 | 0.01 | 0.23 | 0.56 | FALSE |
| 12 | 0.90 | 0.18 | 0.02 | 0.15 | 0.55 | FALSE |
| 13 | 0.75 | 0.28 | 0.02 | 0.08 | 0.50 | FALSE |
| 14 | 2.07 | 0.00 | 0.00 | 0.72 | 0.83 | FALSE |
| 15 | 2.72 | 0.00 | 0.01 | 0.30 | 0.64 | FALSE |
| 16 | 2.12 | 0.03 | 0.02 | 0.20 | 0.55 | FALSE |
| 17 | 2.32 | 0.05 | 0.03 | 0.16 | 0.55 | FALSE |
| 18 | 1.65 | 0.14 | 0.04 | 0.04 | 0.40 | FALSE |
| 19 | 2.30 | 0.01 | 0.01 | 0.65 | 0.83 | FALSE |
| 20 | 2.34 | 0.00 | -0.01 | 0.61 | 0.83 | FALSE |
| 21 | 2.52 | 0.00 | 0.00 | 0.77 | 0.83 | FALSE |
| 22 | 2.87 | 0.00 | 0.00 | 0.74 | 0.83 | FALSE |
| 23 | 3.42 | 0.00 | -0.01 | 0.26 | 0.60 | FALSE |
| 24 | 2.88 | 0.00 | -0.01 | 0.67 | 0.83 | FALSE |
| 25 | 2.51 | 0.00 | 0.01 | 0.36 | 0.71 | FALSE |
| 26 | 2.21 | 0.01 | 0.02 | 0.19 | 0.55 | FALSE |
| 27 | 3.19 | 0.00 | -0.01 | 0.56 | 0.83 | FALSE |
| 28 | 3.20 | 0.00 | -0.01 | 0.61 | 0.83 | FALSE |
| 29 | 2.43 | 0.38 | 0.04 | 0.40 | 0.75 | FALSE |
| 30 | 2.19 | 0.45 | 0.06 | 0.21 | 0.55 | FALSE |
| 31 | 2.48 | 0.00 | -0.01 | 0.52 | 0.83 | FALSE |
| 32 | 2.47 | 0.00 | 0.00 | 0.70 | 0.83 | FALSE |

```
R version 3.5.1 (2018-07-02)
Platform: x86_64-apple-darwin14.5.0 (64-bit)
Running under: macOS High Sierra 10.13.2

Matrix products: default
BLAS: /System/Library/Frameworks/Accelerate.framework/Versions/A/Frameworks/vecLib.framework/Versions/A/libBLAS.dylib
LAPACK: /System/Library/Frameworks/Accelerate.framework/Versions/A/Frameworks/vecLib.framework/Versions/A/libLAPACK.dylib

locale:
[1] en_CA.UTF-8/en_CA.UTF-8/en_CA.UTF-8/C/en_CA.UTF-8/en_CA.UTF-8

attached base packages:
[1] stats     graphics  grDevices utils     datasets  methods   base

other attached packages:
[1] bindrcpp_0.2.2 ggpubr_0.1.8   magrittr_1.5   ggplot2_3.0.0  reshape2_1.4.3
[6] digest_0.6.16  dplyr_0.7.6    plyr_1.8.4

loaded via a namespace (and not attached):
 [1] Rcpp_0.12.17     pillar_1.3.0     compiler_3.5.1   bindr_0.1.1
 [5] base64enc_0.1-3  tools_3.5.1      uuid_0.1-2       jsonlite_1.5
 [9] evaluate_0.11    tibble_1.4.2     gtable_0.2.0     pkgconfig_2.0.2
[13] rlang_0.2.1      IRdisplay_0.5.0  IRkernel_0.8.12  repr_0.15.0
[17] withr_2.1.2      stringr_1.3.1    cowplot_0.9.3    grid_3.5.1
[21] tidyselect_0.2.4 glue_1.3.0       R6_2.2.2         pbdZMQ_0.3-3
[25] purrr_0.2.5      scales_1.0.0     htmltools_0.3.6  assertthat_0.2.0
[29] colorspace_1.3-2 labeling_0.3     stringi_1.2.4    lazyeval_0.2.1
[33] munsell_0.5.0    crayon_1.3.4
```
