## Supplementary material for "A framework for evaluating correspondence between brain images using anatomical fiducials": S4_PHASE4_template_to_template.pdf

### Phase 4: Template-to-Template Evaluation

This notebook contains results evaluating the correspondence between BigBrain and ICBM2009b. Use Case for BigBrain vs ICBM2009b (Sym).

### Validation of AFID Placements

Two expert raters and one additional expert rater overlooking the placements.

'Total: 0.59 +/- 0.40 mm; Outliers: 0/128 (0.00%)'

| template | mean | sd |
|---|---|---|
| BigBrain | 0.63 | 0.50 |
| MNI152NLin2009bSym | 0.55 | 0.26 |

### BigBrainSym versus ICBM2009b Sym

BigBrain has been pre-registered to ICBM2009b Sym and available as a package online. Here we evaluated the spatial correspondence between these two templates.

'Total: 2.16 +/- 1.99 mm'

#### Is there any correlation of the errors reported with FLE?

Here we take our computed AFLE values for BigBrain-Sym and ICBM2009b-Sym and find that there is no correlation with the AFRE found.

```
Warning message in cor.test.default(summary_bbsym_vs_sym$AFRE, summary_bbsym_df$mean, :
“Cannot compute exact p-value with ties”

        Kendall's rank correlation tau

data:  summary_bbsym_vs_sym$AFRE and summary_bbsym_df$mean
z = -0.37303, p-value = 0.7091
alternative hypothesis: true tau is not equal to 0
sample estimates:
        tau
-0.04641778
```

```
Warning message in cor.test.default(summary_bbsym_vs_sym$AFRE, summary_sym_df$mean, :
“Cannot compute exact p-value with ties”

        Kendall's rank correlation tau

data:  summary_bbsym_vs_sym$AFRE and summary_sym_df$mean
z = 0.56765, p-value = 0.5703
alternative hypothesis: true tau is not equal to 0
sample estimates:
       tau
0.07063576
```

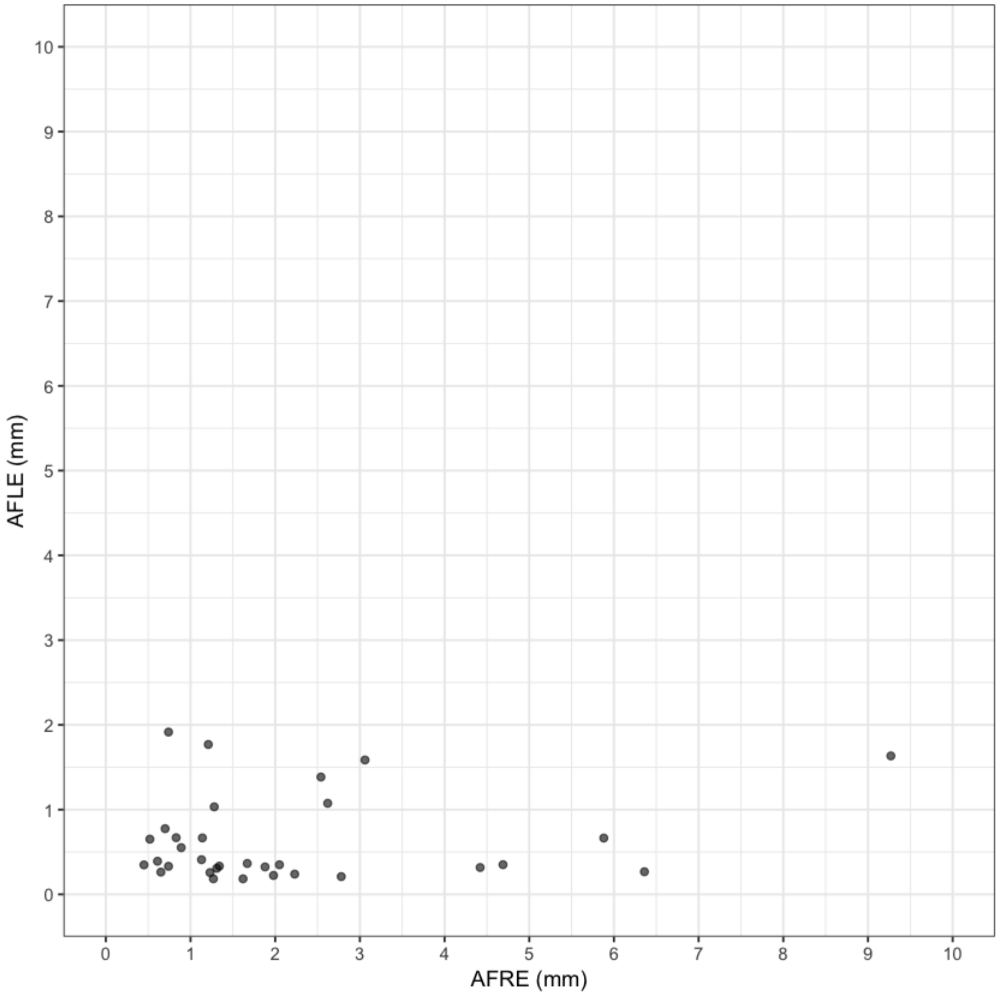


#### BigBrainSym versus ICBM2009b Asym

Here we evaluated the spatial correspondence between BigBrainSym and MNI2009bAsym (asymmetric) knowing that BigBrainSym was registered to MNI2009bSym rather than MNI2009bAsym. AFRE should be higher than for MNI2009bSym.

'Total: 2.30 +/- 1.83 mm'

#### ICBM2009b: Sym versus Asym

Here we evaluated the distance between AFIDs for ICBM2009b sym and asym templates. Note that calling the difference AFRE is not technically correct as the two templates are not aligned to one another. However, the syntax was kept the same for simplicity.

'Total: 0.88 +/- 0.68 mm'

##### Is there any correlation of the errors reported with FLE?

Here we take our computed AFLE values for ICBM2009b-Asym and ICBM2009b-Sym and find that there is no correlation with the AFRE found.

```
Warning message in cor.test.default(summary_asym_vs_sym$AFRE, summary_sym_df$mean, :
“Cannot compute exact p-value with ties”

        Kendall's rank correlation tau

data:  summary_asym_vs_sym$AFRE and summary_sym_df$mean
z = 1.687, p-value = 0.09161
alternative hypothesis: true tau is not equal to 0
sample estimates:
      tau
0.2101014
```

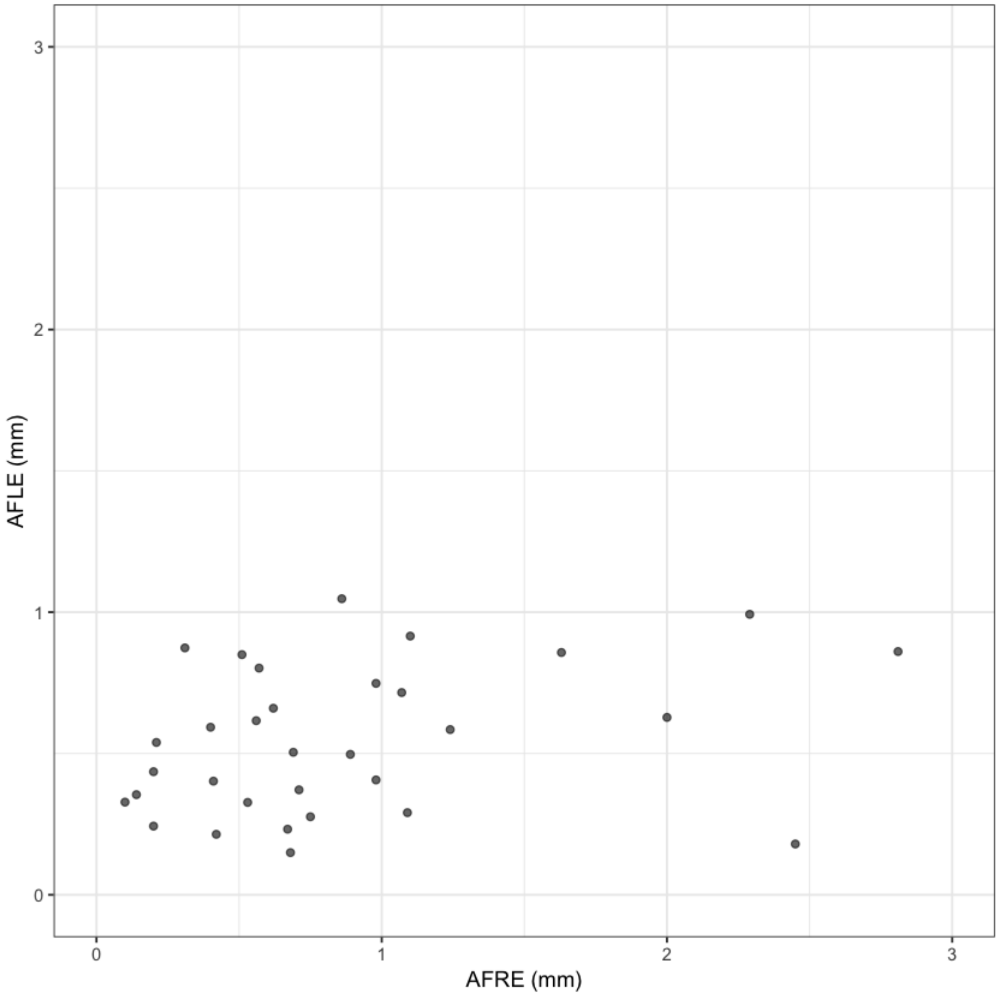


| | AFID | Description | AFRE for BigBrainSym vs MNI2009bSym | Star: BigBrain and Sym | AFRE for BigBrainSym vs MNI2009bAsym | Star: BigBrain and Asym | AFRE for MNI2009b: Asym vs Sym | Star: Asym vs Sym |
|---|---|---|---|---|---|---|---|---|
| **3** | 03 | infracollicular sulcus | 6.36 | * | 5.48 | * | 0.98 | |
| **9** | 09 | L inferior LMS | 2.78 | * | 2.48 | * | 0.68 | |
| **10** | 10 | culmen | 9.27 | * | 9.39 | * | 0.21 | |
| **14** | 14 | pineal gland | 4.42 | * | 4.16 | * | 0.41 | |
| **16** | 16 | L LV at AC | 2.05 | * | 1.22 | | 0.86 | |
| **20** | 20 | splenium | 2.23 | * | 2.20 | * | 0.10 | |
| **22** | 22 | L AL temporal horn | 4.69 | * | 3.44 | * | 2.45 | * |
| **26** | 26 | L inferior AM temporal horn | 1.88 | | 2.58 | * | 0.98 | |
| **27** | 27 | R indusium griseum origin | 1.21 | | 3.60 | * | 2.81 | * |
| **28** | 28 | L indusium griseum origin | 0.74 | | 2.88 | * | 2.29 | * |
| **29** | 29 | R ventral occipital horn | 2.54 | * | 3.99 | * | 1.63 | |
| **30** | 30 | L ventral occipital horn | 5.88 | * | 4.22 | * | 2.00 | * |
| **31** | 31 | R olfactory sulcal fundus | 2.62 | * | 1.84 | | 1.10 | |
| **32** | 32 | L olfactory sulcal fundus | 3.06 | * | 4.21 | * | 1.24 | |

```
R version 3.5.1 (2018-07-02)
Platform: x86_64-apple-darwin14.5.0 (64-bit)
Running under: macOS High Sierra 10.13.2

Matrix products: default
BLAS: /System/Library/Frameworks/Accelerate.framework/Versions/A/Frameworks/vecLib.framework/Versions/A/libBLAS.dylib
LAPACK: /System/Library/Frameworks/Accelerate.framework/Versions/A/Frameworks/vecLib.framework/Versions/A/libLAPACK.dylib

locale:
[1] en_CA.UTF-8/en_CA.UTF-8/en_CA.UTF-8/C/en_CA.UTF-8/en_CA.UTF-8

attached base packages:
[1] stats     graphics  grDevices utils     datasets  methods   base

other attached packages:
[1] ggplot2_3.0.0  reshape2_1.4.3 digest_0.6.16  plyr_1.8.4

loaded via a namespace (and not attached):
 [1] Rcpp_0.12.17     bindr_0.1.1      magrittr_1.5     tidyselect_0.2.4
 [5] munsell_0.5.0    uuid_0.1-2       colorspace_1.3-2 R6_2.2.2
 [9] rlang_0.2.1      dplyr_0.7.6      stringr_1.3.1    tools_3.5.1
[13] grid_3.5.1       gtable_0.2.0     withr_2.1.2      htmltools_0.3.6
[17] assertthat_0.2.0 lazyeval_0.2.1   tibble_1.4.2     crayon_1.3.4
[21] bindrcpp_0.2.2   IRdisplay_0.5.0  purrr_0.2.5      repr_0.15.0
[25] base64enc_0.1-3  IRkernel_0.8.12  glue_1.3.0       evaluate_0.11
[29] pbdZMQ_0.3-3     stringi_1.2.4    pillar_1.3.0     compiler_3.5.1
[33] scales_1.0.0     jsonlite_1.5     pkgconfig_2.0.2
```
