## Supplementary material for "A framework for evaluating correspondence between brain images using anatomical fiducials": S5_afids_protocol.pdf

afids / **afids-protocol**

Branch: master

afids-protocol / **protocol.md**

Find file | Copy path

**Jonathan Lau** updated figures and text 465894e on Jan 9

3 contributors

236 lines (154 sloc) 13.1 KB

Raw | Blame | History

### Fiducial Protocol

#### Preparation

- Download and use Slicer 4.8.1

#### Naming Scheme for Fiducial Files

- [VolumeID]*[Contrast]*[Rater]_[N] (e.g. MNI2009b_T1_JL_1_20170511.fcsv)
  - [VolumeID] = the identifier for the volume on which you are performing the fiducial placements; for the tutorial it will be one of the well known MRI templates:
    - Colin27: average of 27 Colin brains
    - MNI2009b: average of 152 healthy controls
    - Agile12v1.0: average of 12 healthy controls at 7T
  - [Contrast] = T1, T2, PD, other (typically will be T1)
  - [Rater] = the unique identifier for the rater performing the fiducial placement; convention will be first initial and last name to prevent overlap
  - [N] = reference for fiducial placement session (helpful if performing placements more than once; starting with 1)

- [YYYYMMDD] = year month and date

#### Place AC and PC Fiducials

After creating the Markups list as described above, the process begins by placing the anterior commissure landmark (for more information, see the Section below entitled Fiducial Placement Details). Once placed, click on the corresponding row in the Markups Module list and change the Name to "1" and Description to "AC". Next, place the posterior commissure landmark (see Section Fiducial Placement Details). Click on the corresponding row in the Markups Module list and change the Name to "2" and Description to "PC".

#### Create new AC-PC Transform

The images first have to be AC-PC aligned using the **ACPC Transform Module**. Within the Markups Module, we next have to create two new lists: “ACPC Line” and “Midline” by clicking "Create New MarkupsFiducial as..." then entering the name "ACPC Line" and separately "Midline".

Next copy both AC and PC fiducials into “ACPC Line” and “Midline” Lists. To do this, right click on the individual fiducials (AC and PC) and select “Copy fiducial to another list” then add to both “ACPC Line” and “Midline” lists.

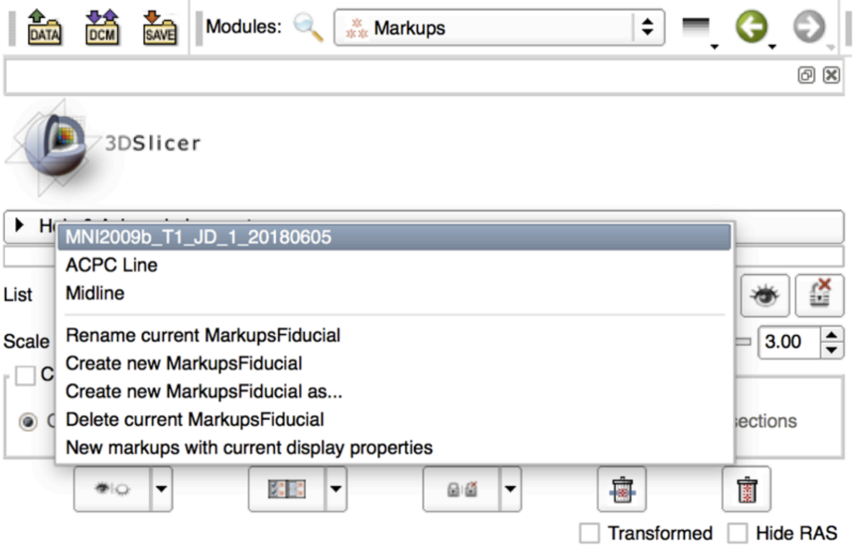

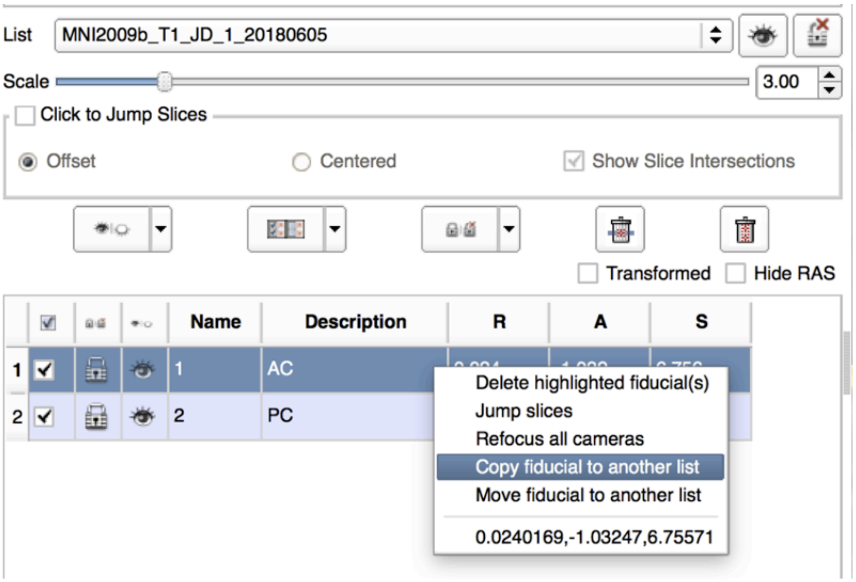


Next, you will need to place a third fiducial marker in the “Midline” list. Select “Midline” list and place a fiducial marker in the superior interpeduncular fossa (Fid03 below; axial view to targeting) and name it Midline. Alternatively, use the intermamillary sulcus (Fid11 below).

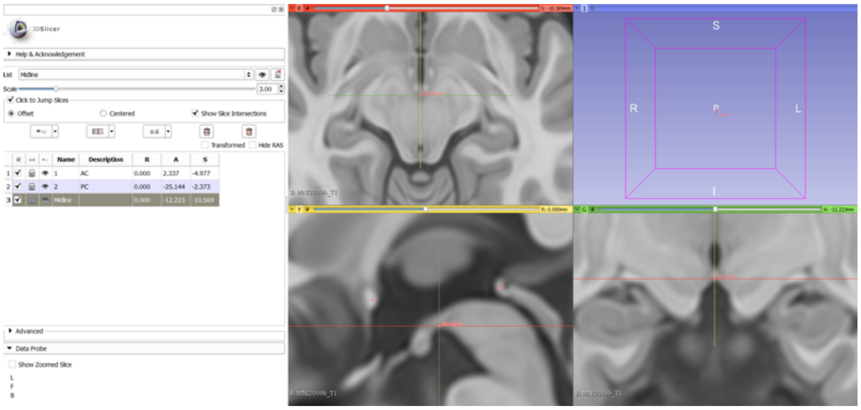

Next, you will perform the ACPC transform to realign the MRI volume. Under modules select **Registration>Specialized>ACPC Transform** (Left image below). Under the “ACPC transform” tab, select the “Parameter Set” dropdown tab and select “ACPC transform”. Below that is the “Transform Panel”. Under “ACPC Line” dropdown tab select “ACPC Line”, under “Midline” dropdown tab select “Midline”, and under “Output transform” dropdown tab select “New Linear Transform” and name it “Output Transform”. Next, click “Apply” at the bottom of the window (image below).

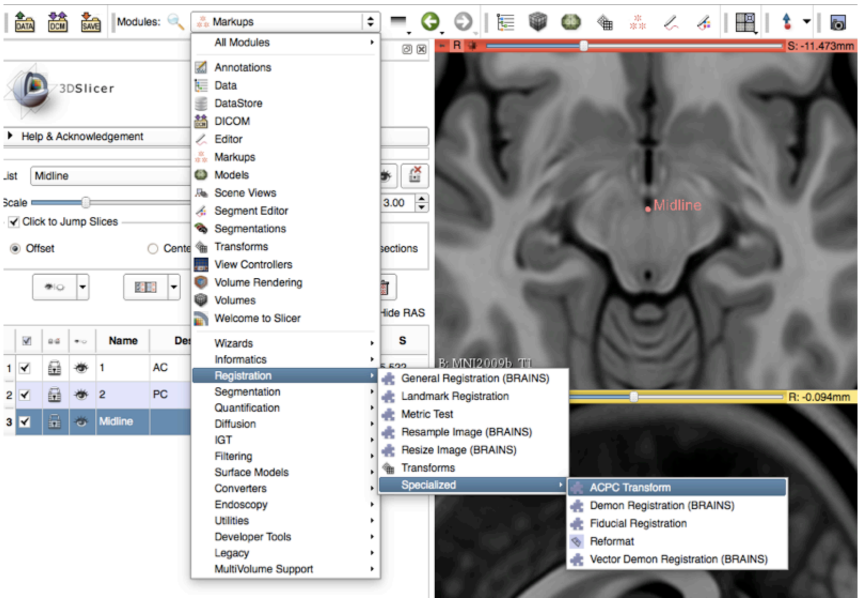


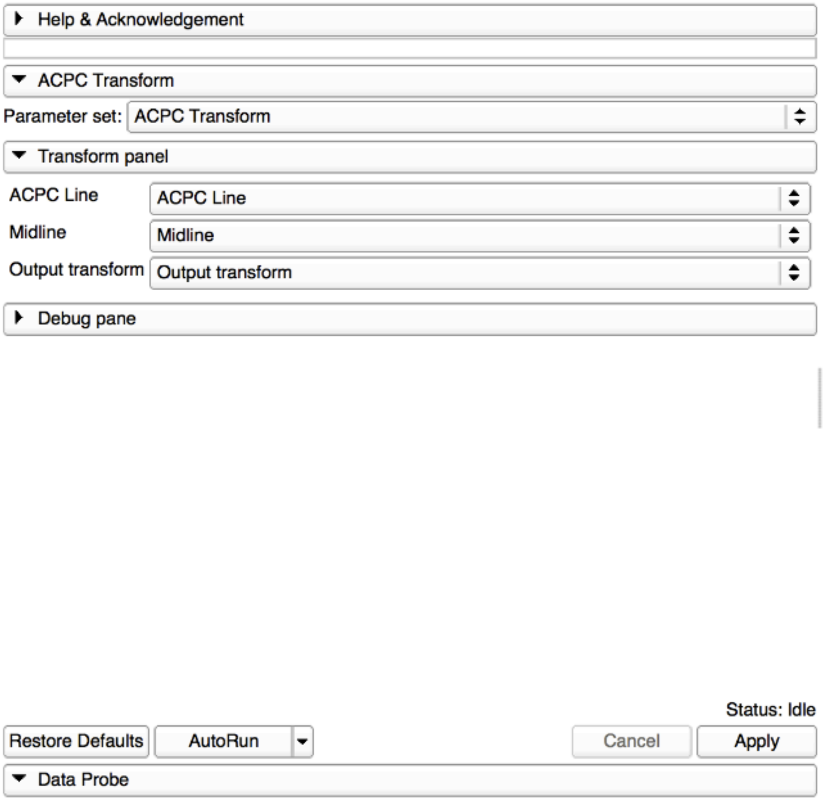


Finally, under “Modules” go to “Transforms.” Under the “Active Transform” dropdown tab select “Output Transform” as the active transform. Next, under the “Apply Transform” menu, Select all 4 lists (i.e the original loaded volume, ACPC Line, Midline etc.) and click the arrow pointing right to transfer them from “Transformable” to “Transformed.” Your image and fiducials should shift to realign with the proper orthogonal axis.

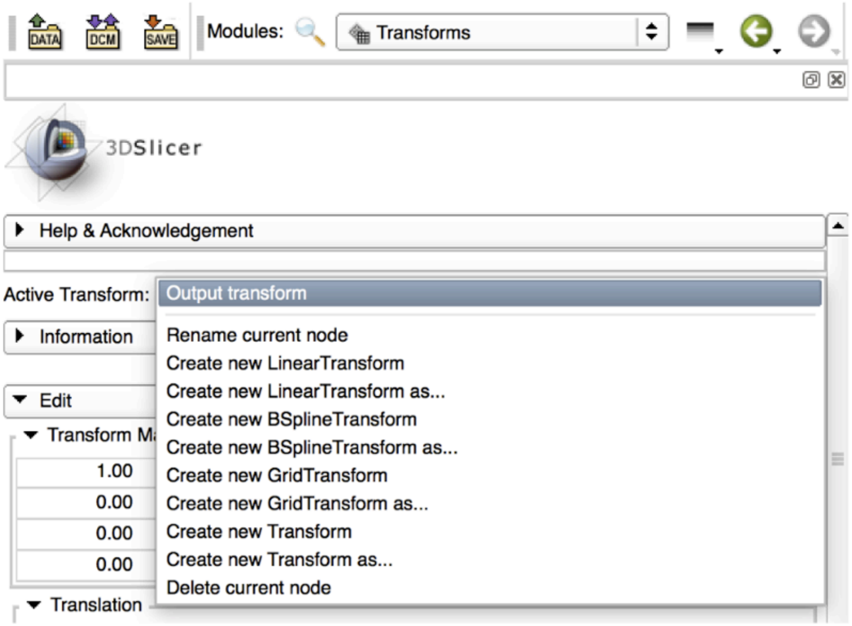


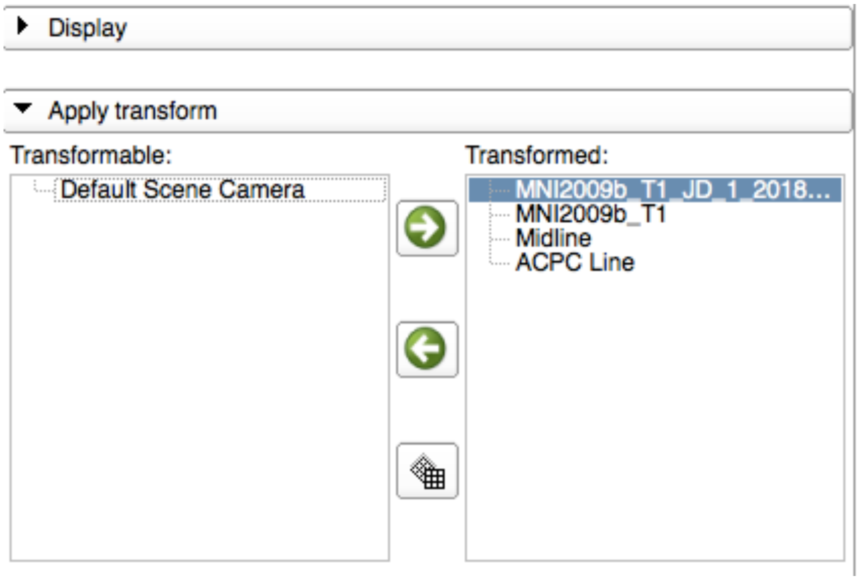


### General Fiducial Placement Strategies

Use the **"Jump to Slice"** feature to center your view on the fiducial of interest and ensure that the placed landmark appears accurate on all three standard views (axial, sagittal, coronal). Once a fiducial is placed, **dragging** the fiducial can allow for more refined placement. Holding down **shift** centers the view in all views on the cursor (use along with crosshair function). If a given fiducial is classified as **[midline]**, jump to an existing midline fiducial (e.g. AC or PC) and start by placing the fiducial on the **sagittal** view and refine placement using the other views. Try to place fiducials at the **boundary/edge** of the feature of interest. For some of the fiducials, the instructions for placement will explicitly say to place the landmark using information mostly from one view (e.g. axial view for olfactory sulcus). Be aware that changing the windowing of your images (and lighting in the room) may affect your perception of where landmarks should be placed. When you're satisfied with the location of a fiducial, **lock it in place** to prevent yourself from displacing it later. **NOTE: there is no UNDO feature for fiducial placements.**

### Placement of Fiducial Series

Create a new Markup list entitled **Fid32_[VolumeID]*[Rater]*[N]**. Click on **ACPC Line** and copy over AC and PC to your new list by right clicking each fiducial, choosing "Copy fiducial to another list", and selecting **Fid32_[VolumeID]*[Rater]*[N]**. Place the following **30 fiducials**, enter the number corresponding to the fiducial in the Name textbox and enter the underlined anatomical structure in the corresponding Description textbox:

### 1. AC [midline]

- Place at the center of the commissure

01. Anterior Commissure

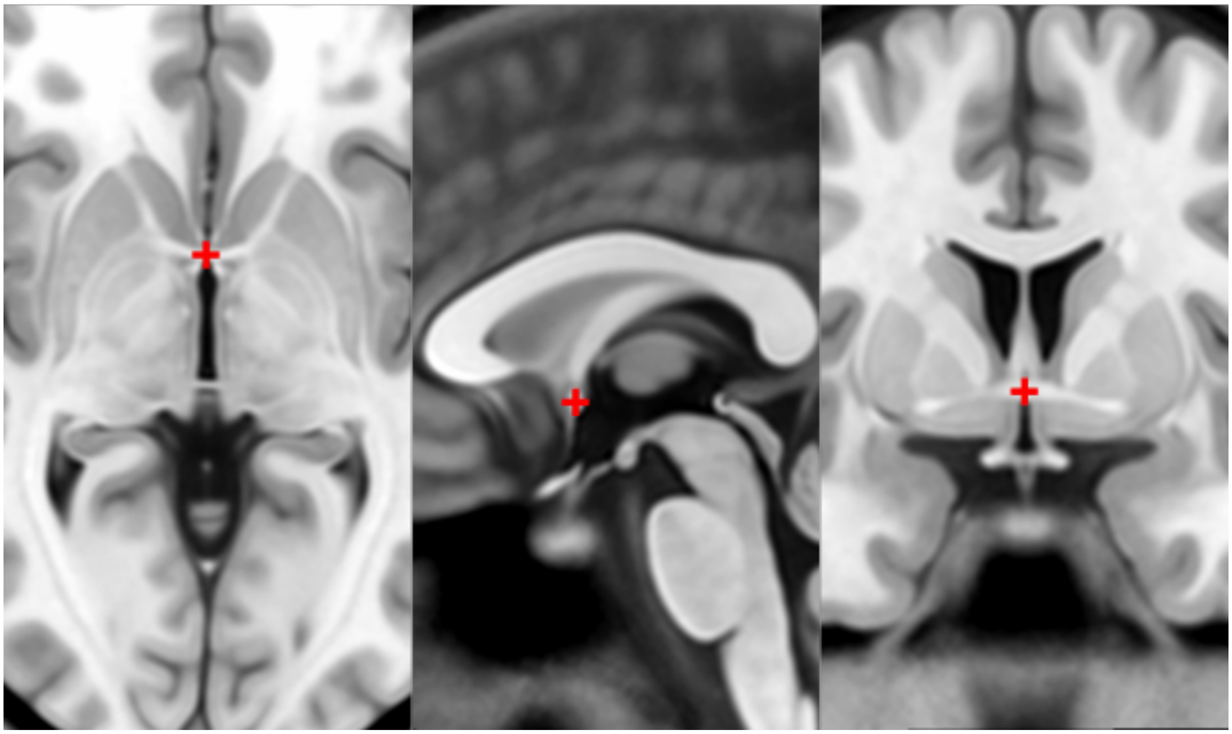

#### 2. PC [midline]

- Place at the center of the commissure

02. Posterior Commissure

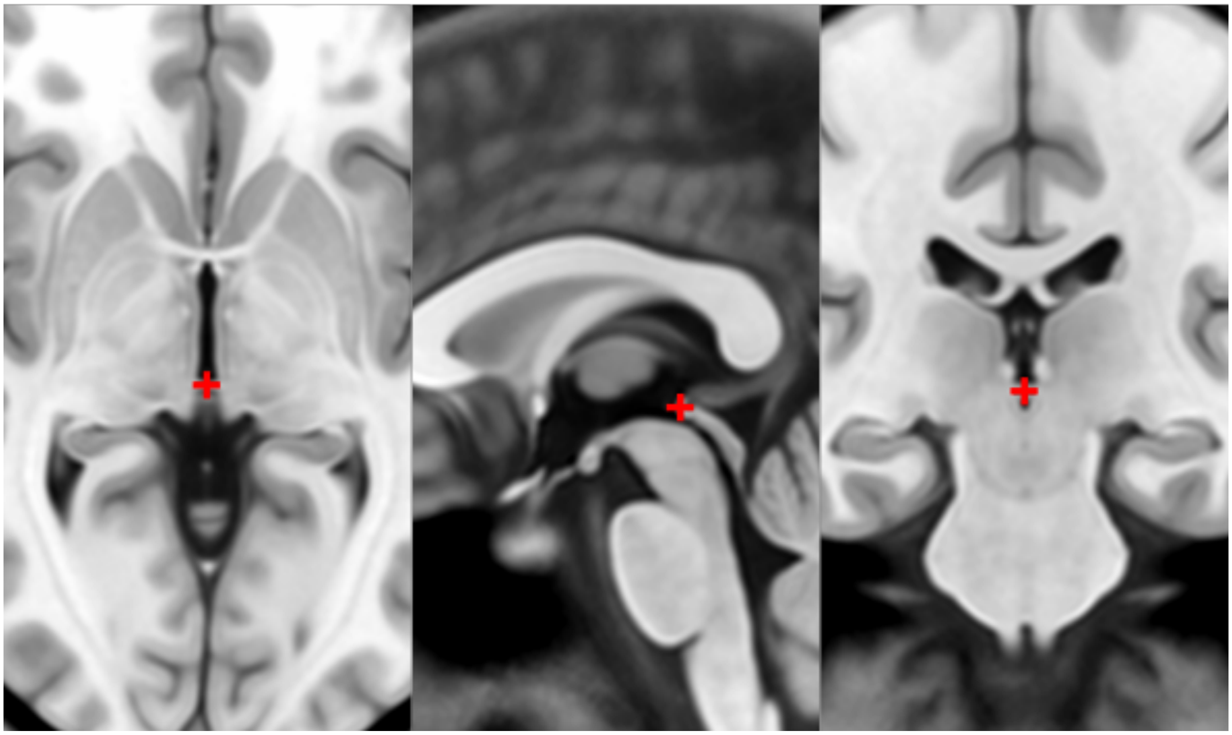

#### 3. infracollicular sulcus [midline]

- Inferior part of sulcus of inferior colliculi at the midline junction of inferior colliculi
- Inferior most boundary of longitudinal intercollicular sulcus

03. Infracollicular Sulcus

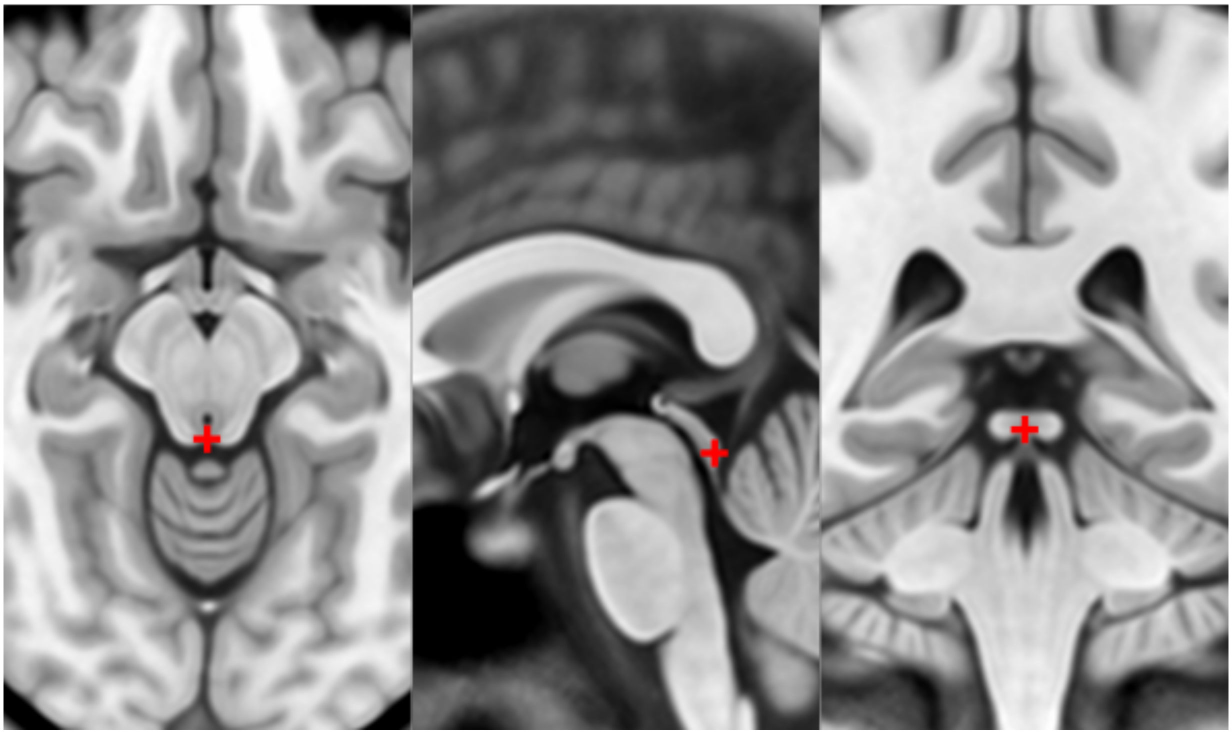

#### 4. PMJ = pontomesencephalic junction [midline]

- At the junction but because the junction tapers off gradually, choose the ventral/inferior/pontine side of the junction using the sagittal and coronal views

04. PMJ

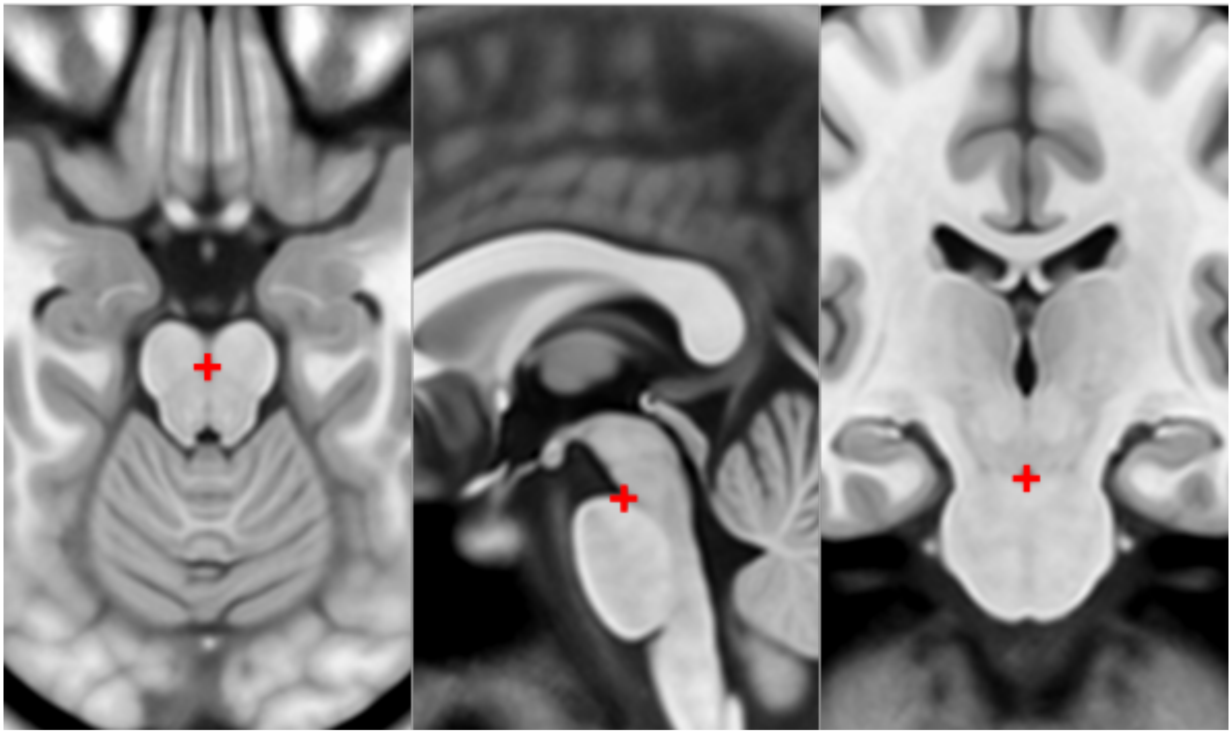

#### 5. superior interpeduncular fossa [midline]

- Most superior axial slice showing the interpeduncular fossa
- Use coronal slice to help optimize location at boundary of 3rd ventricle and surrounding brain
- Commentary: useful fiducial location for DBS since subthalamic nucleus close by

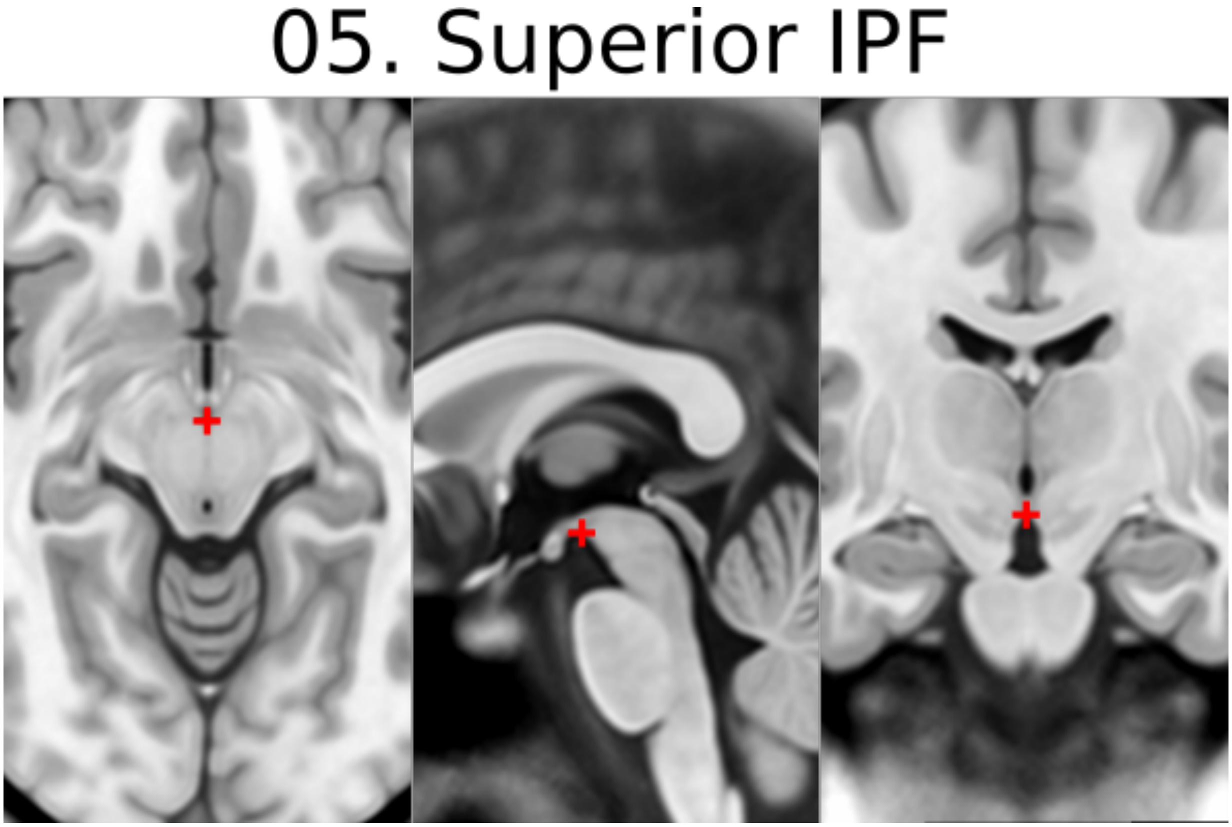


#### 6. R superior LMS = right superior lateral mesencephalic sulcus

- Localize using axial slices; at boundary of CSF and brain
- Use the coronal and sagittal views to optimize the location so that this point is in the angle created in all views

06. Right Superior LMS

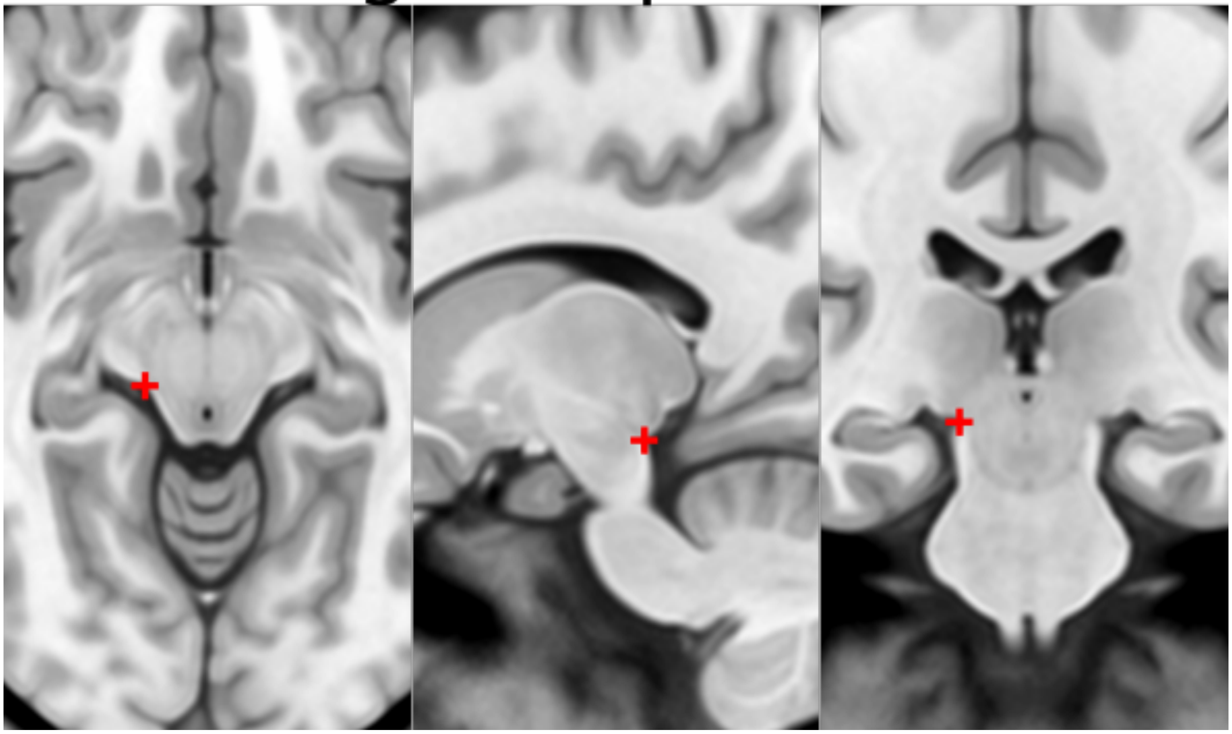

#### 7. L superior LMS = left superior lateral mesencephalic sulcus

- As in 6

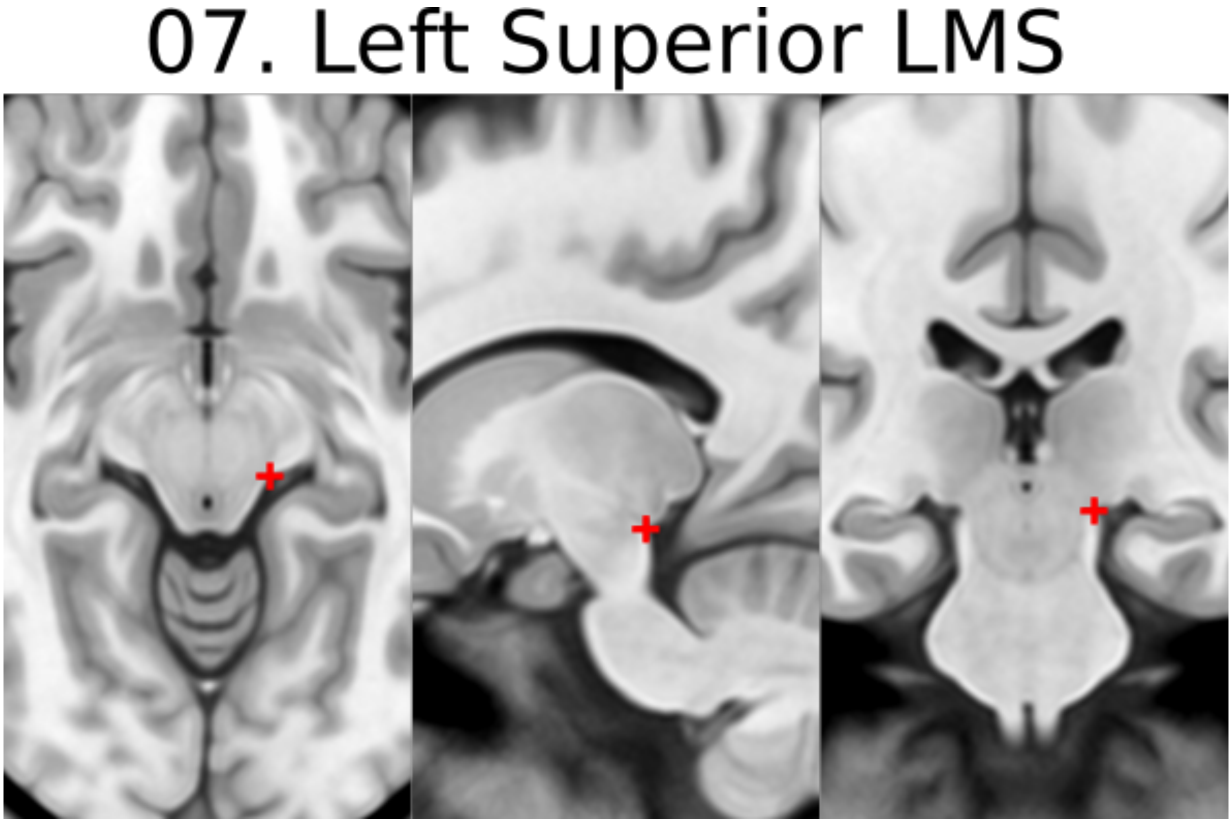


#### 8. R inferior LMS = right inferior lateral mesencephalic sulcus

- Localize at junction between midbrain and pons first using axial slices
- Refine positioning using sagittal view (at the change in angle of brainstem at the PMJ)

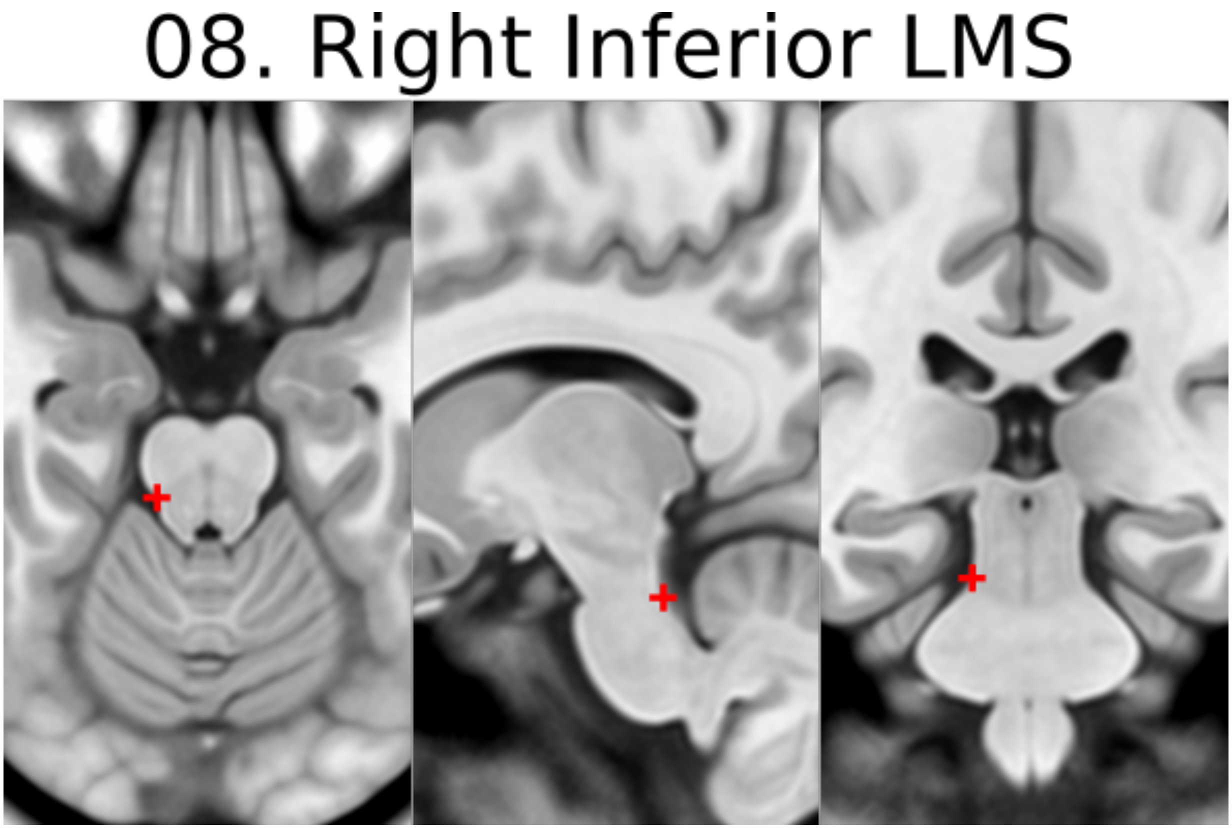


#### 9. L inferior LMS = left inferior lateral mesencephalic sulcus

- As in 8

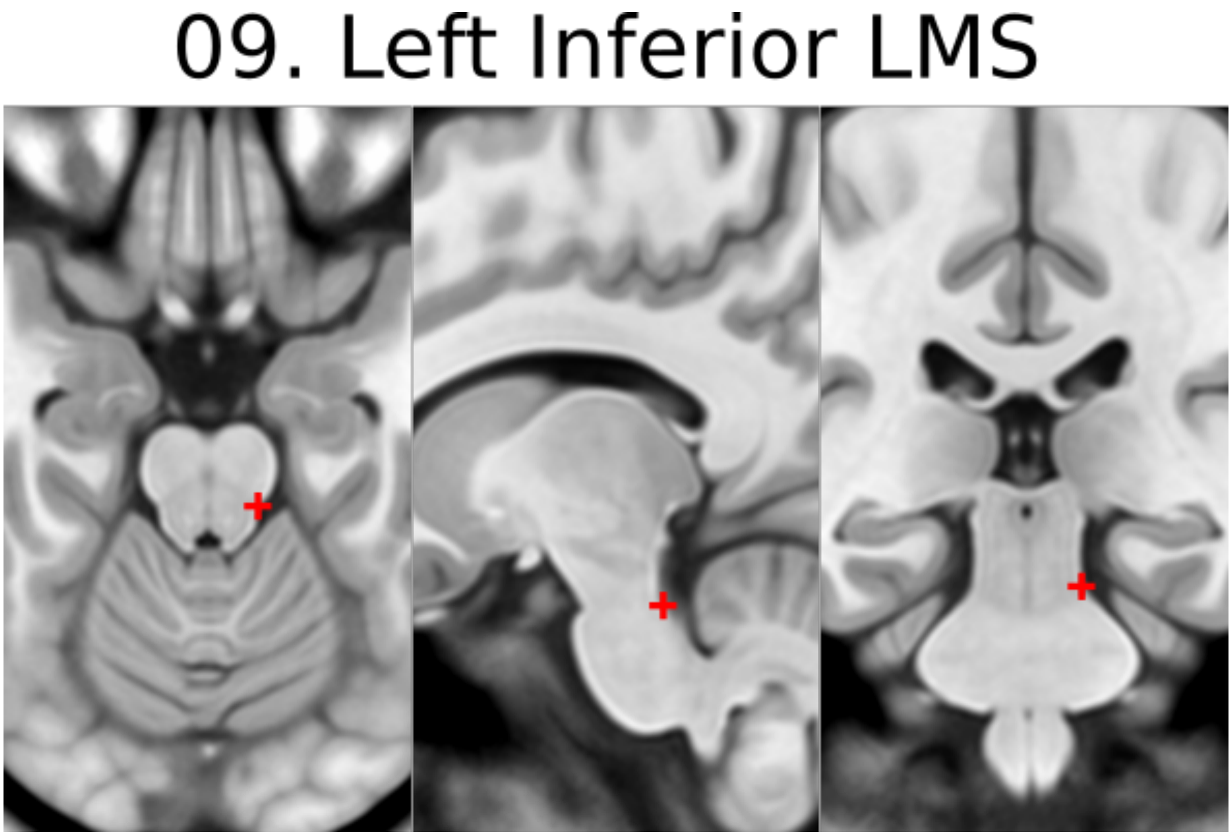


#### 10. Culmen [midline]

- Jump to AC or another midline AFID to get to the mid-sagittal slice, then place using the sagittal view
- Most superior point of cerebellar vermis; one of the vermian lobules

### 10. Culmen

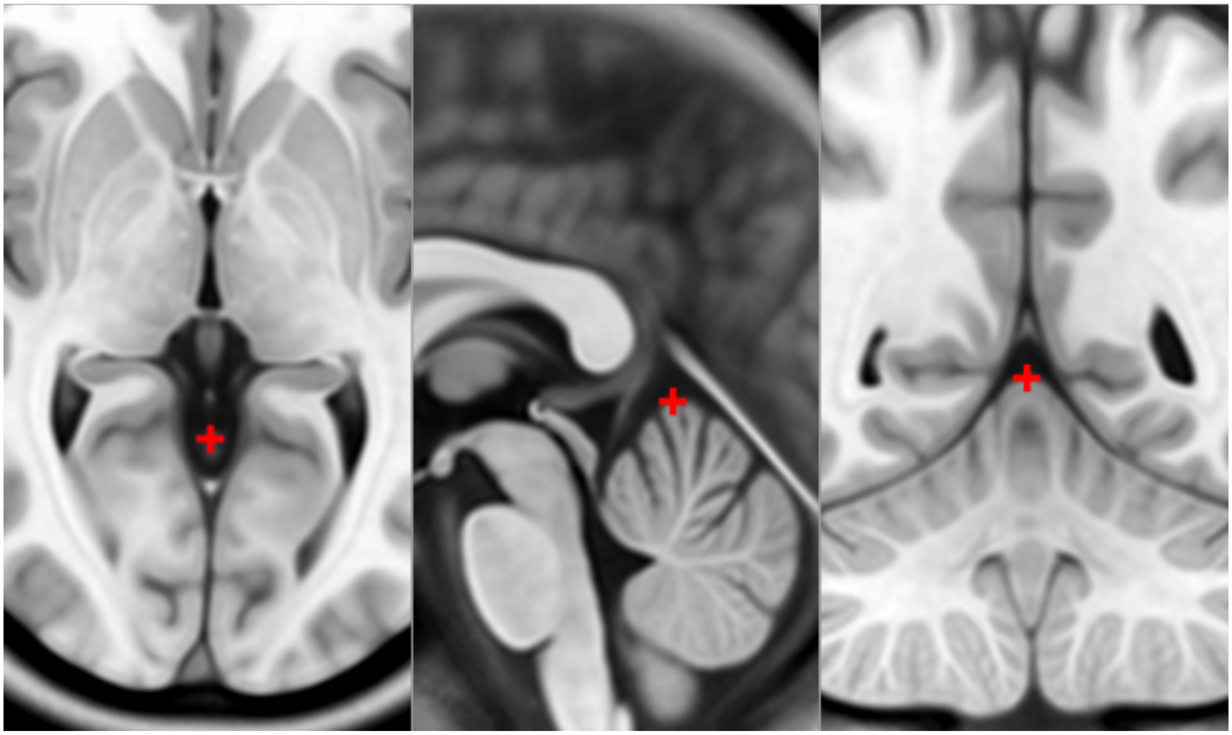

#### 11. Intermammillary sulcus [midline]

- Click to jump to AC landmark and place using the sagittal view
- Midpoint between the mamillary bodies; remember to place at the border of the grey matter

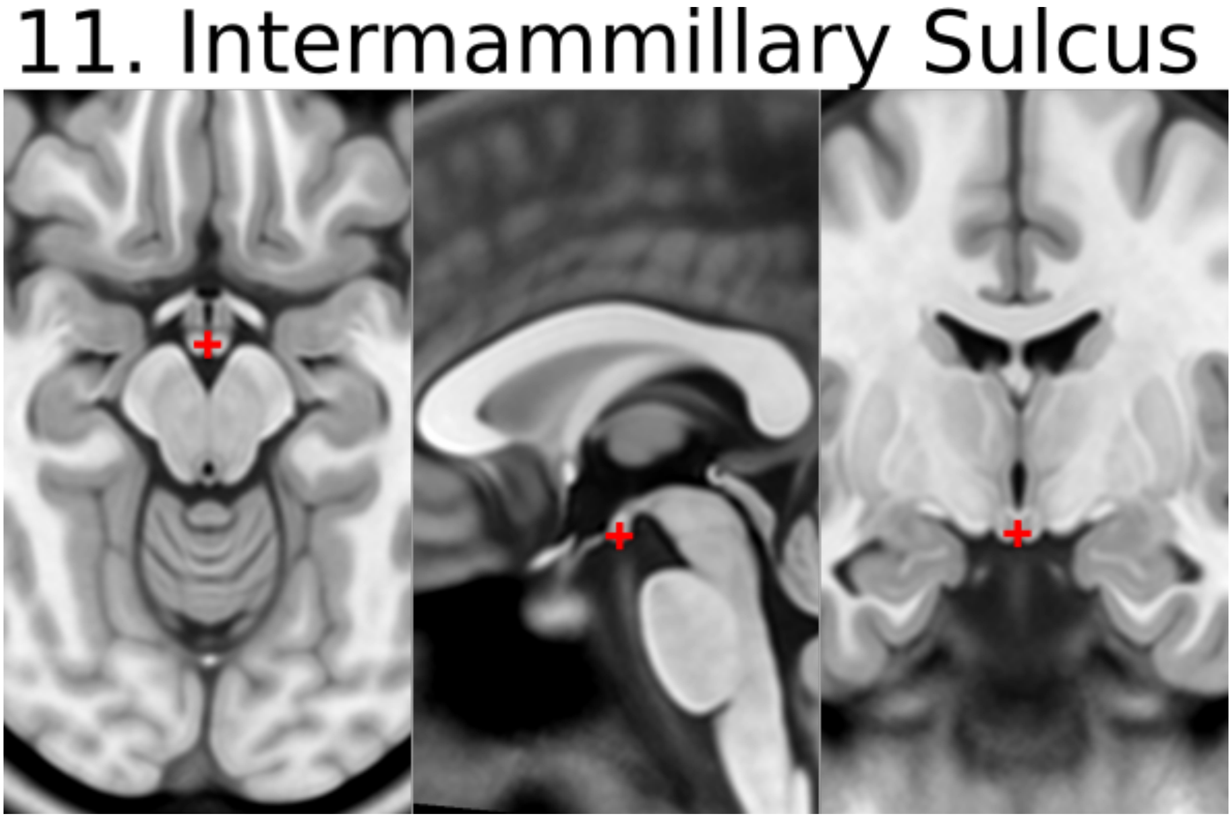


#### 12. R MB = right mammillary body

- Place at the center of the mammillary body

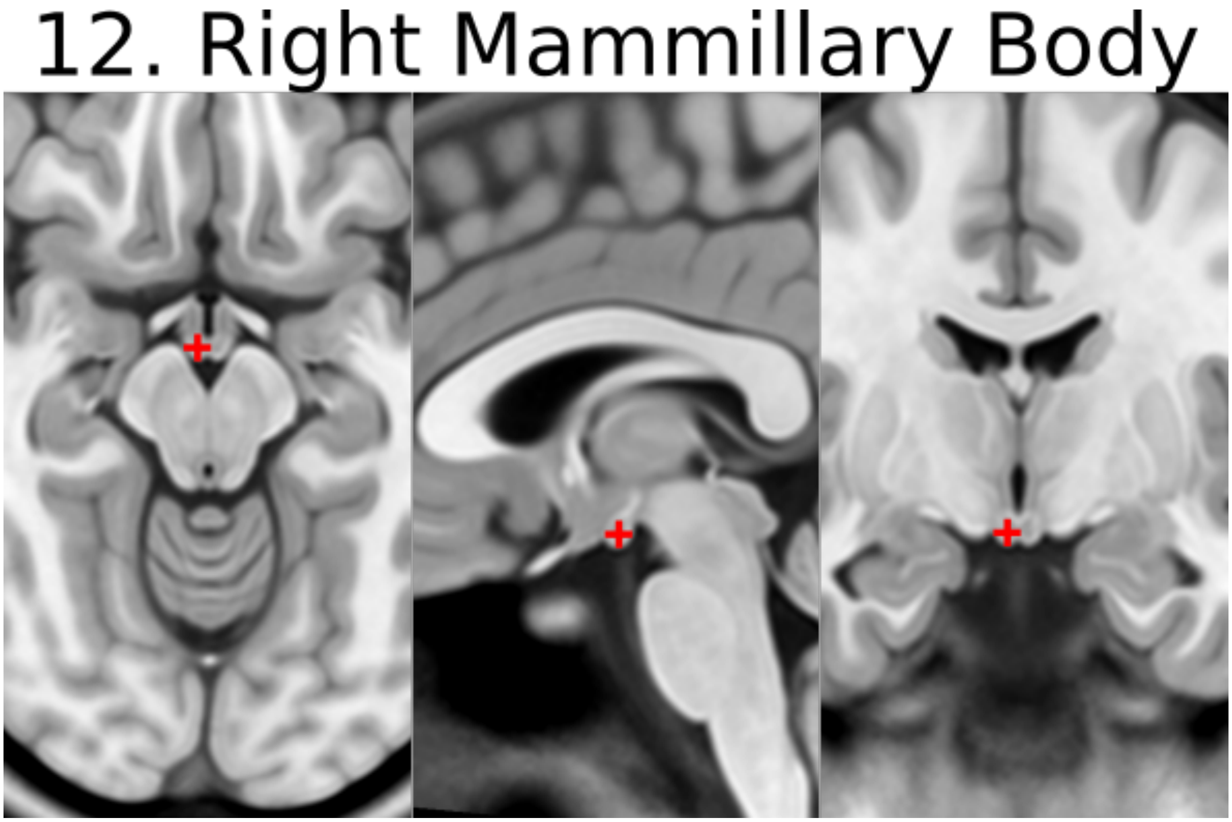

#### 13. L MB = left mamillary body

- As in 12

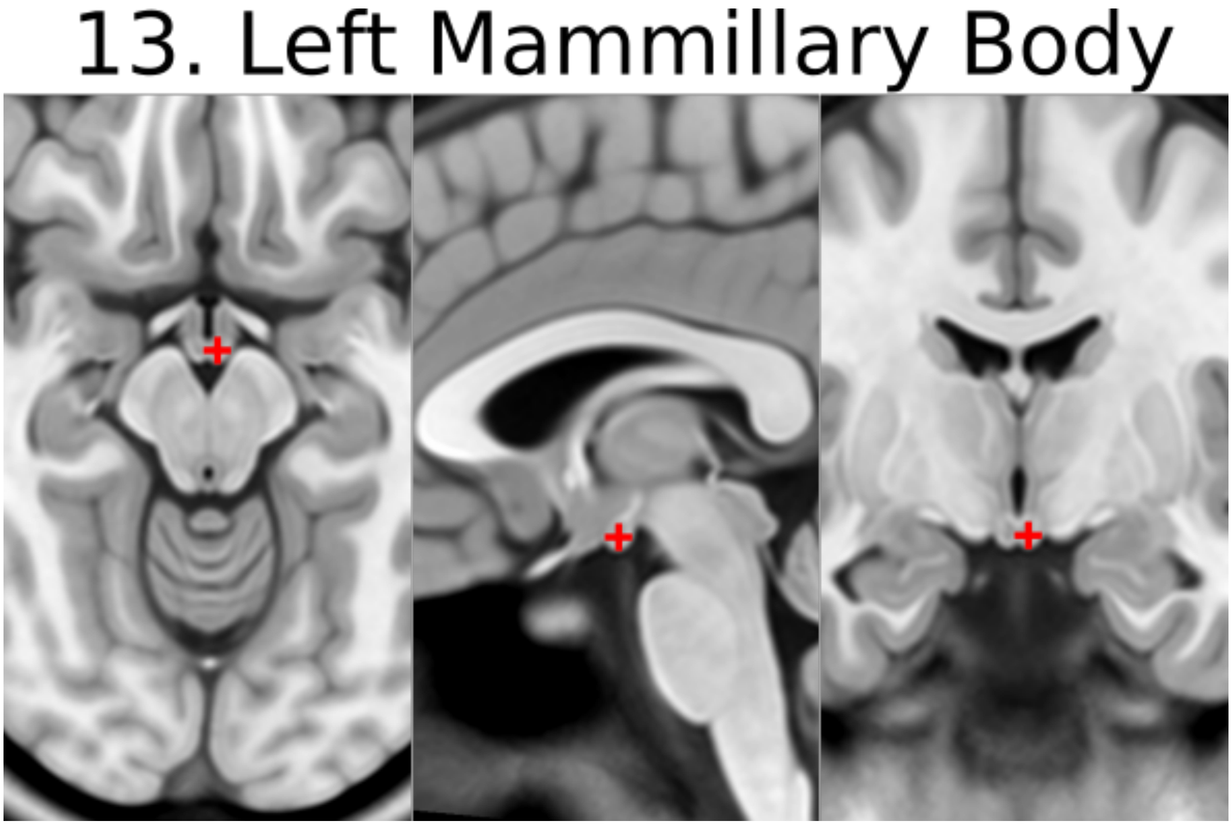


#### 14. pineal gland [midline]

- Click to jump to the AC landmark on the sagittal view and place AFID in the middle of gland (use all views to correctly place this point)
- Occasionally the pineal gland is calcified, which makes it more difficult to find the center of the gland. Be sure to scroll back and forth in all views to find the true center point regardless of asymmetry of calcifications

#### 14. Pineal Gland

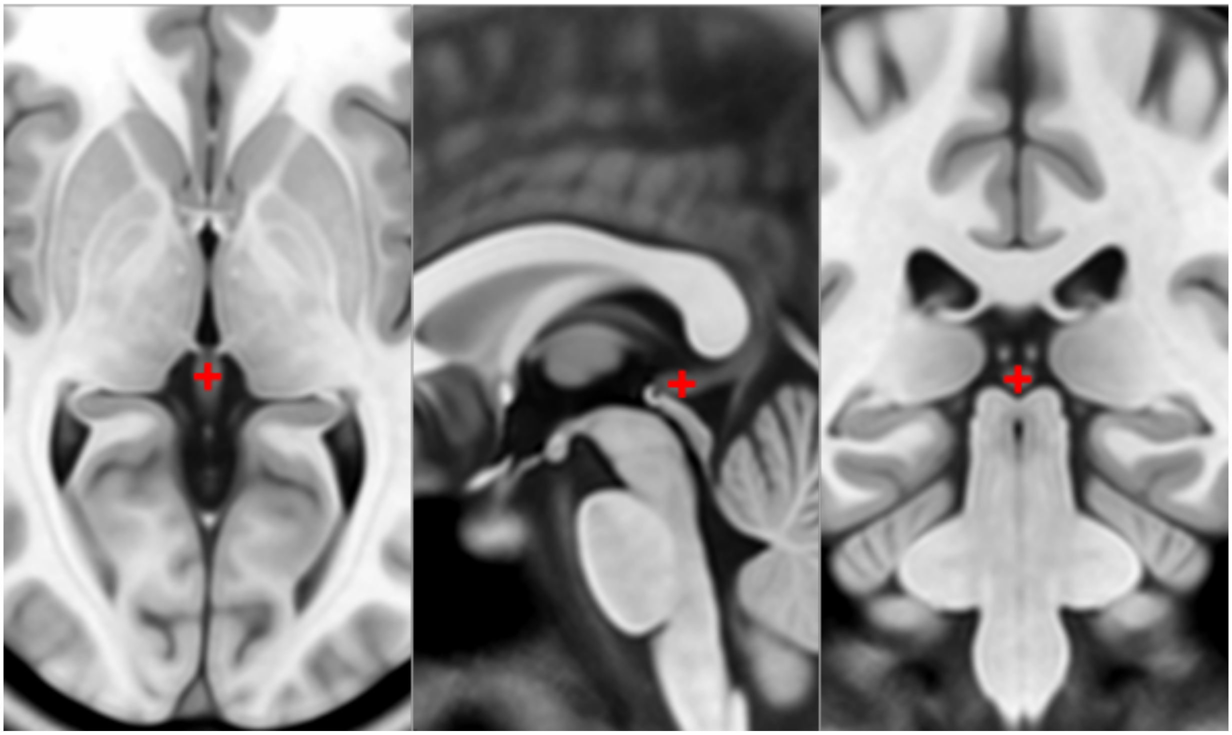

#### 15. R LV at AC = right lateral aspect of frontal horn on coronal section of AC

- Defined at same coronal slice as AC (jump to it)

15. Right LV at AC

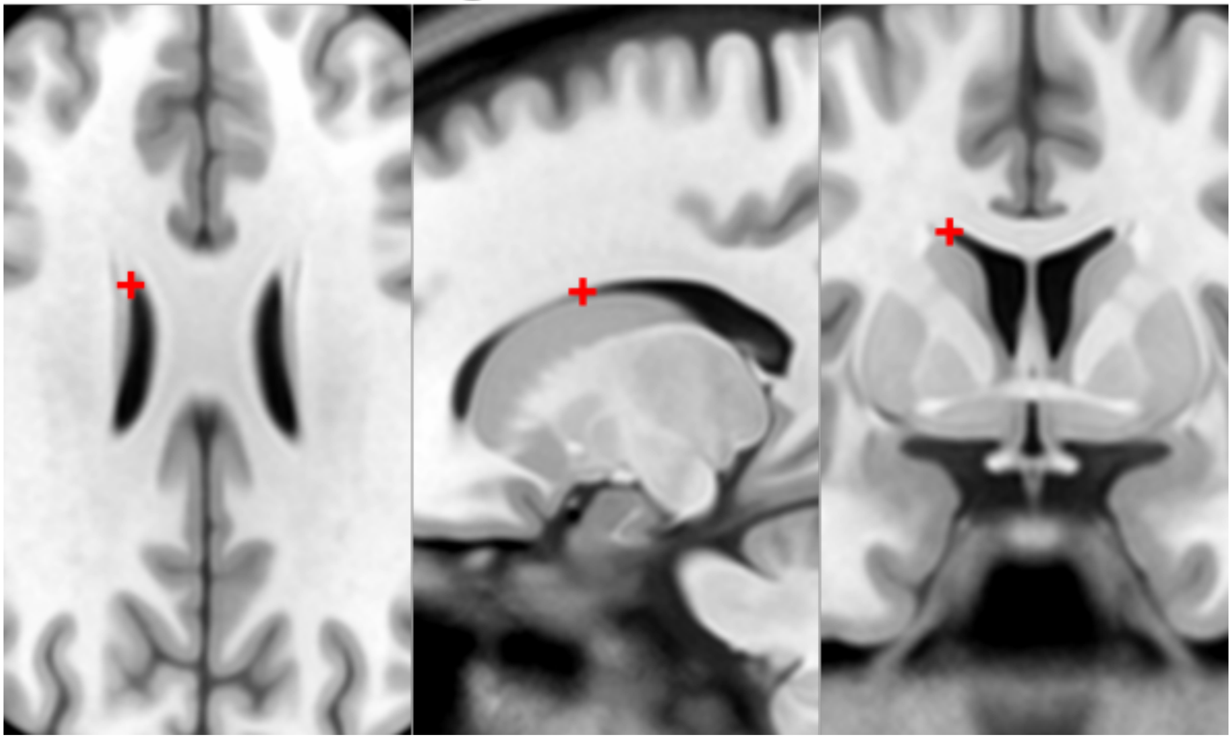

#### 16. L LV at AC = left lateral aspect of frontal horn on coronal section of AC

- As in 15

16. Left LV at AC

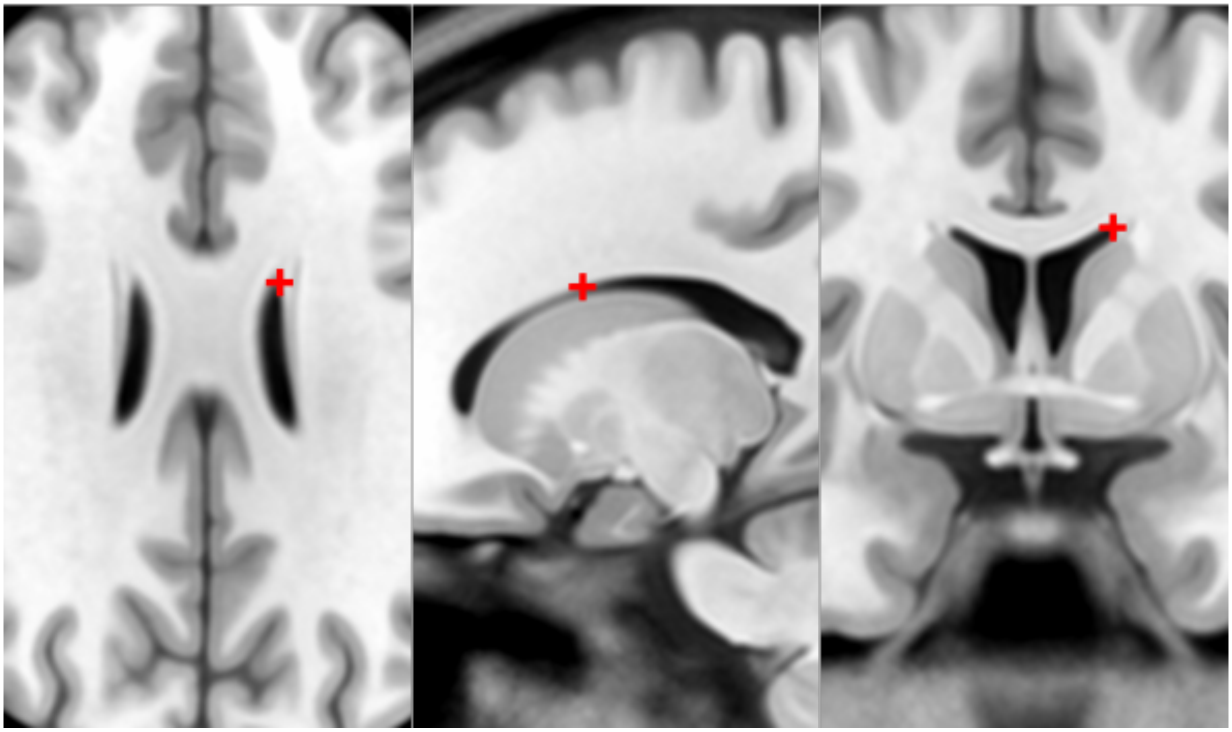

##### 17. R LV at PC = right lateral aspect of frontal horn on coronal section of PC

- Defined at same coronal slice as PC (jump to it)

17. Right LV at PC

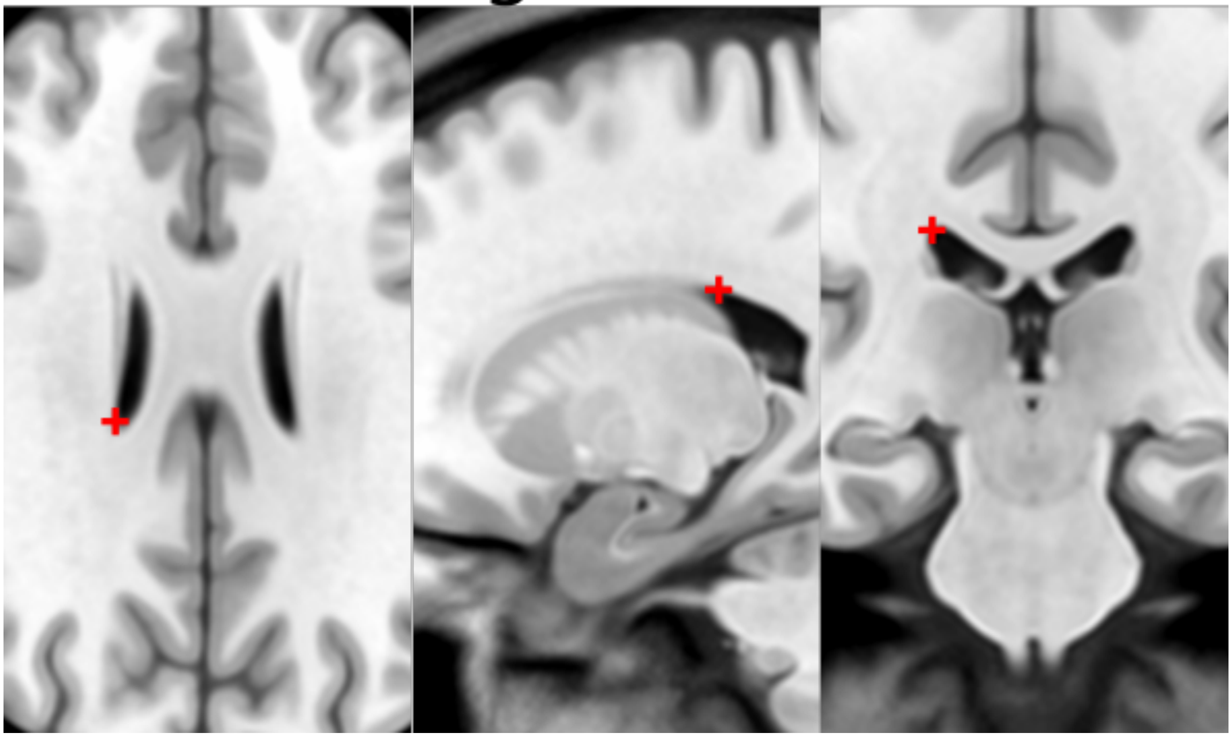

#### 18. L LV at PC = left lateral aspect of frontal horn on coronal section of PC

- As in 17

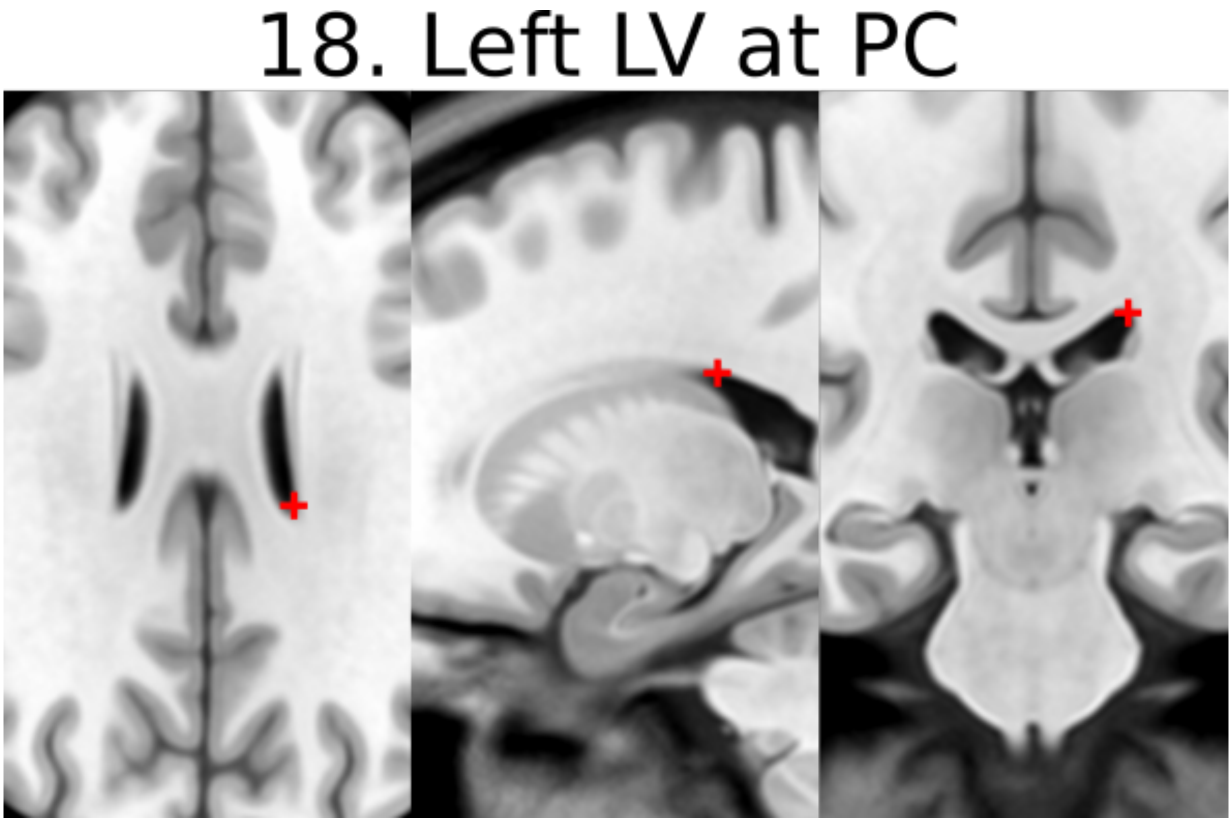


#### 19. Genu of CC = genu of corpus callosum [midline]

- Jump to AC and place using sagittal view
- Optimize using coronal view as most anterior point of the corpus callosum on coronal slice

19. Genu of CC

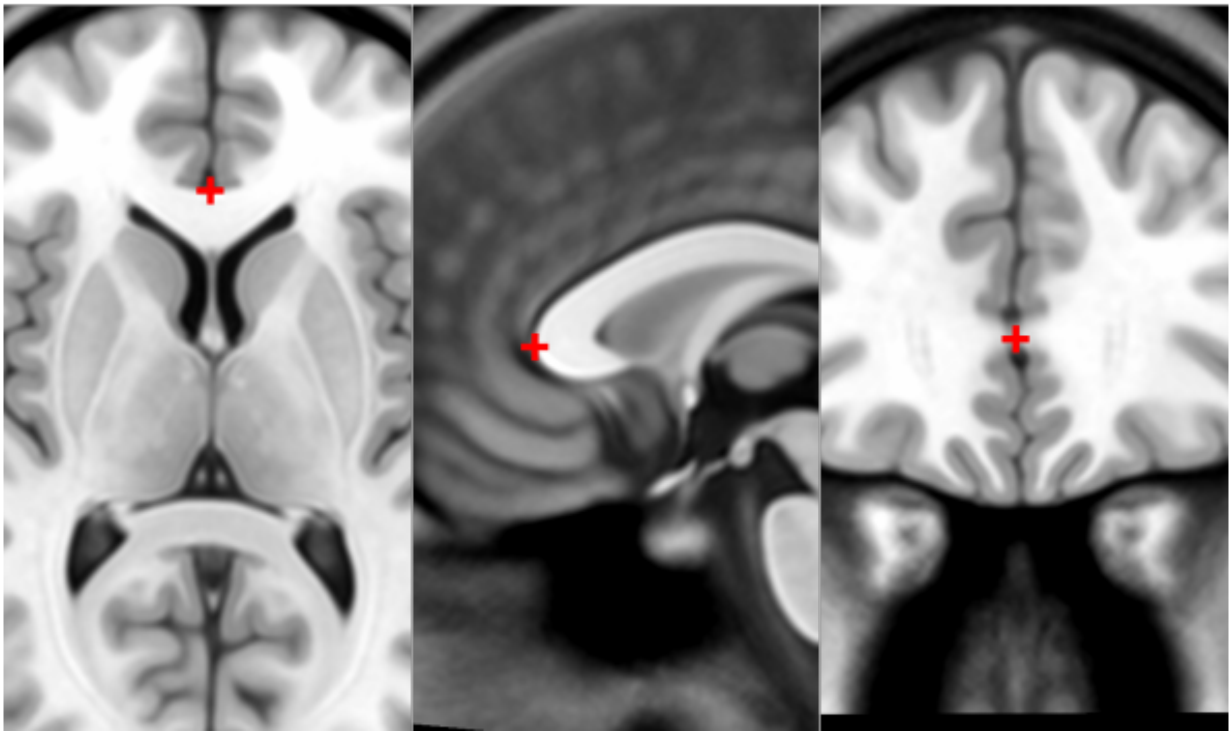

#### 20. Splenium of CC = splenium of the corpus callosum [midline]

- Jump to AC and place using sagittal view.
- Optimize using axial view as the inferior-most point on axial section

20. Splenium of CC

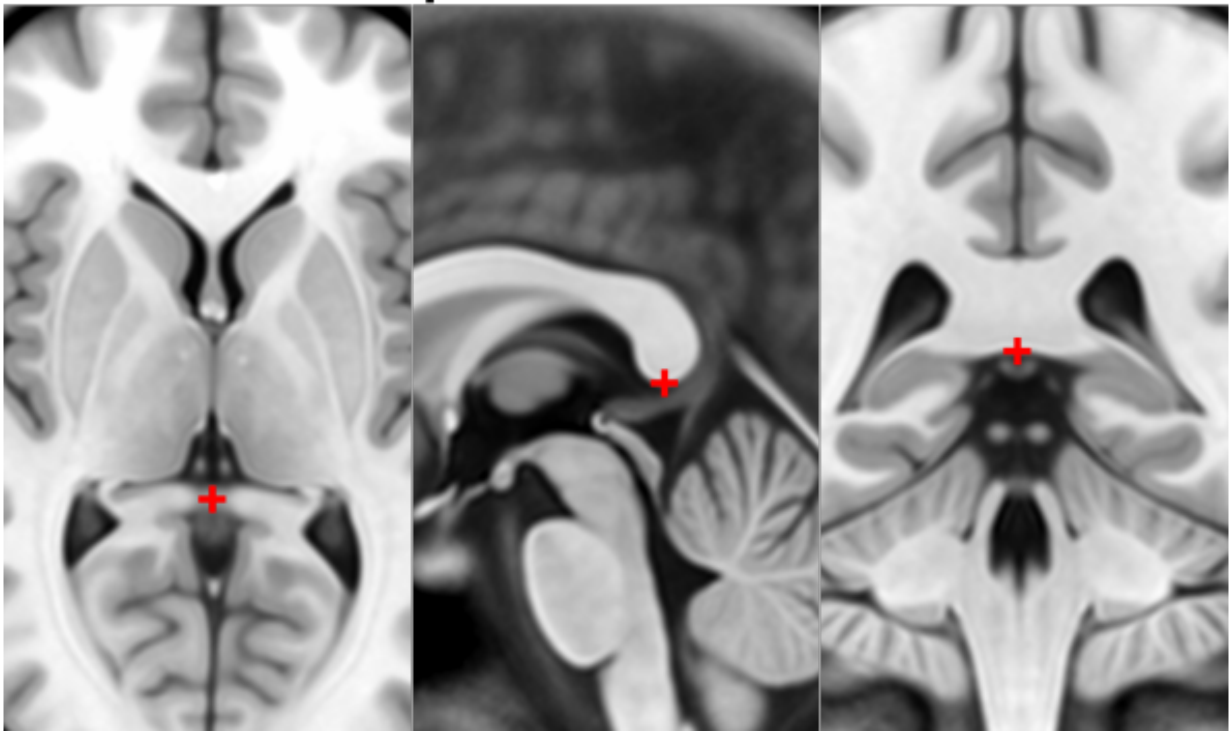

#### 21. R AL temporal horn = right anterolateral temporal horn

- Place using coronal view as the anterior-most (and lateral) point of temporal horn
- Choose a more ventral/inferior point on the coronal view
- Place at the boundary of CSF and brain

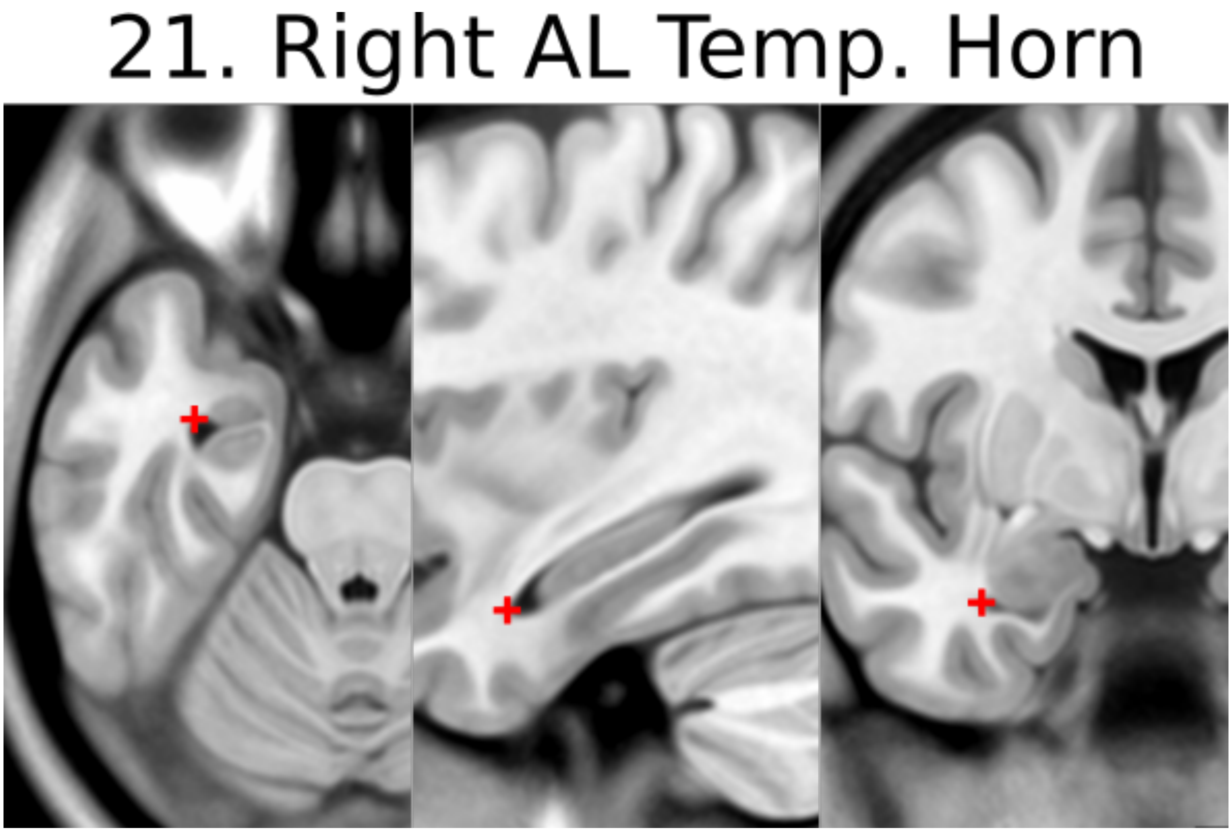


#### 22. L AL temporal horn = left anterolateral temporal horn

- As in 21

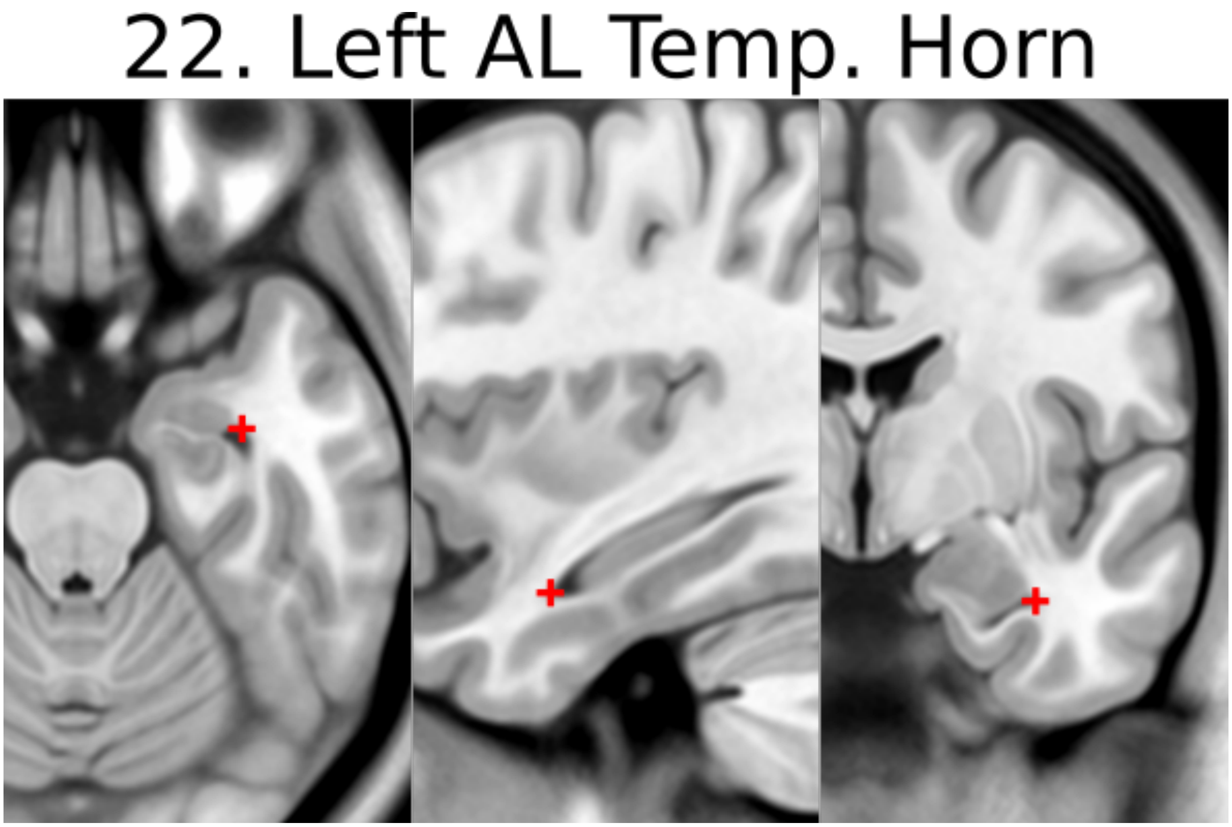


#### 23. R superior AM temporal horn = Rhoton's R uncal recess

- At the superior hippocampal-amygdalar transition area (HATA)
- NOTE: there is also an inferior anteromedial temporal horn
- Rhoton's uncal recess:
  - "narrow medially projecting space between hippocampal head & ventricular surface of amygdala located lateral to uncal apex")
- Place at the boundary of CSF and brain

23. R Sup. AM Temp. Horn

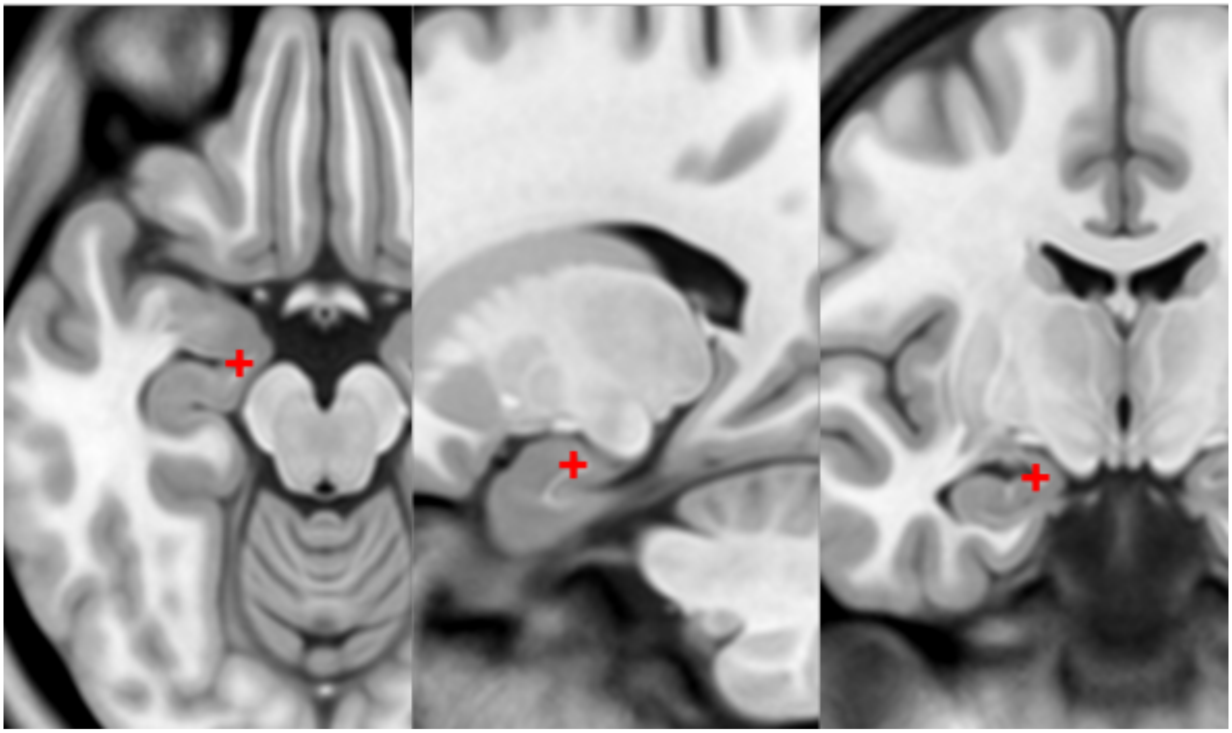

#### 24. L superior AM temporal horn = Rhoton's L uncal recess

- As in 23

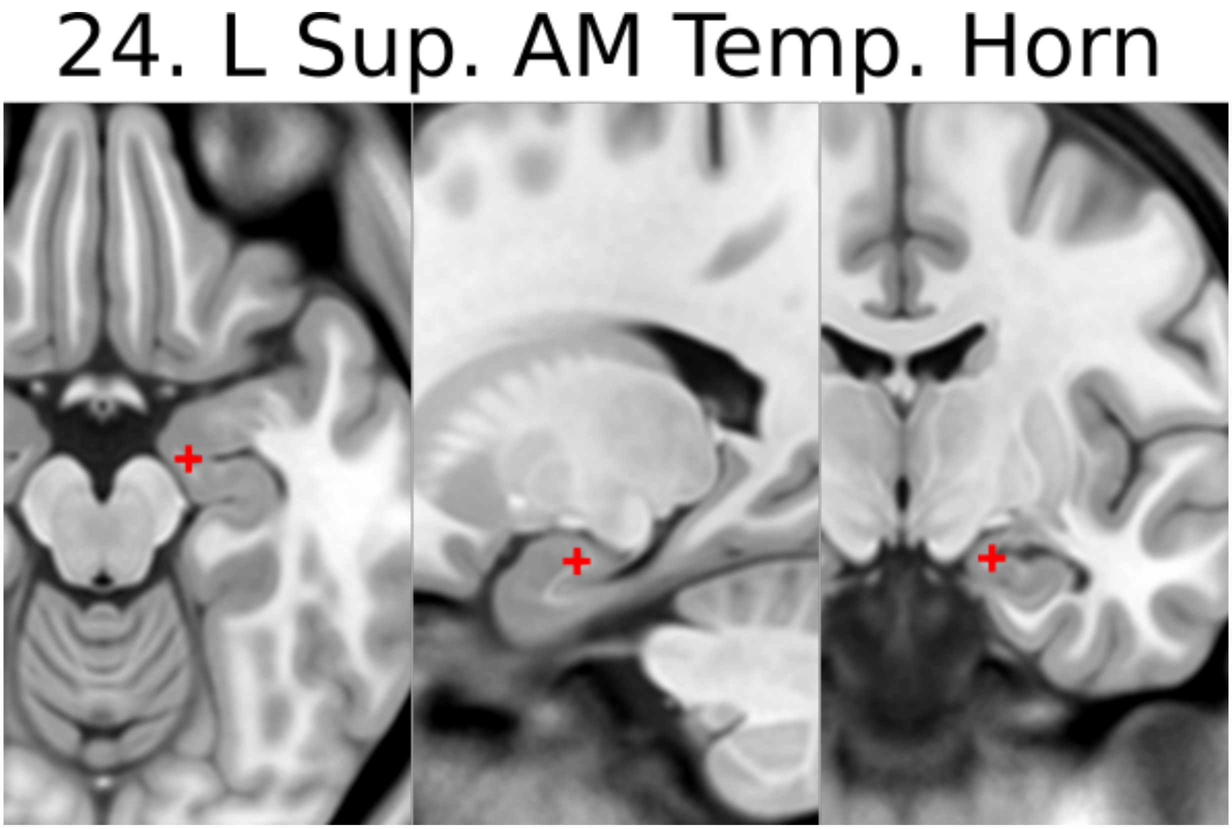


#### 25. R inferior AM temporal horn

- Initially place using coronal view
- Jump to 21 (right AL temporal horn) and scroll to find the most medial (and anterior) showing of the CSF
- Optimize using the axial view again to find the most anteromedial showing of the CSF

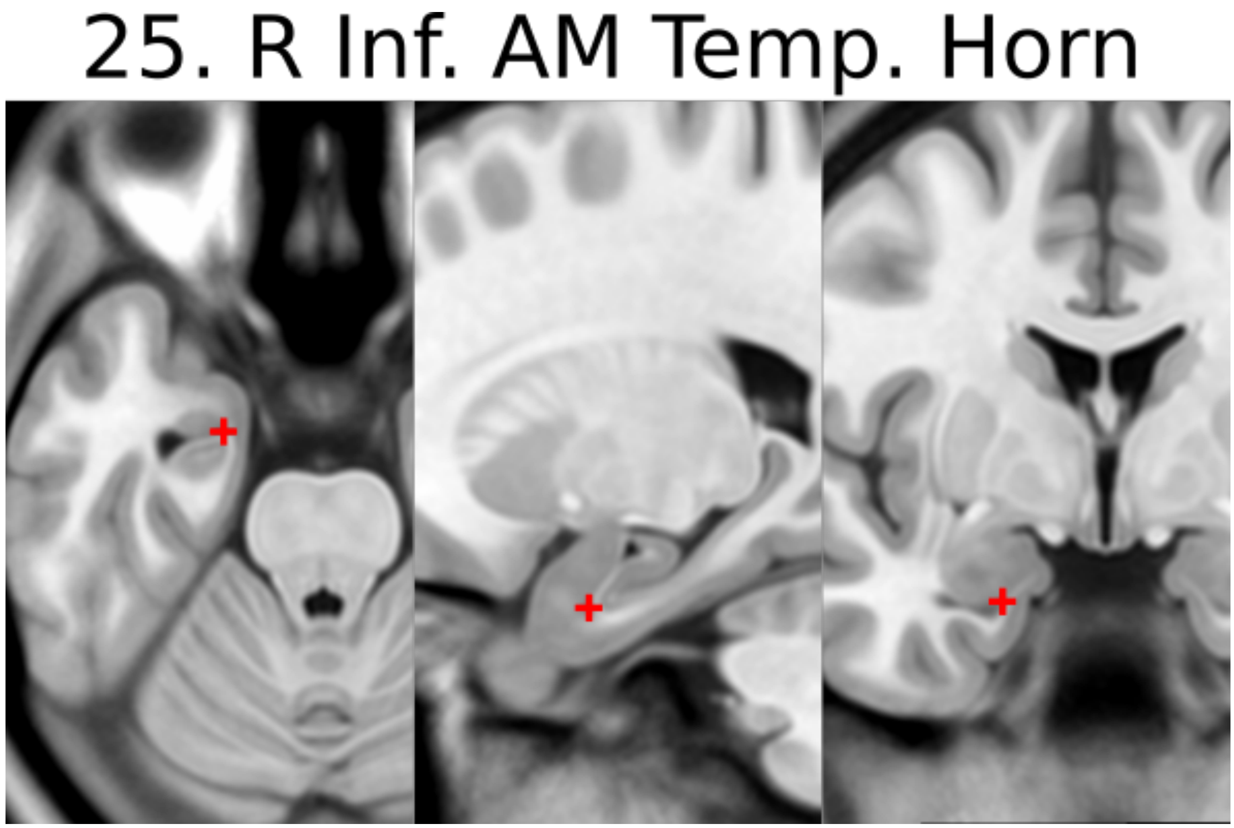


#### 26. L inferior AM temporal horn

- Like in 25
- Jump to 22 (left AL temporal horn) and scroll the find the most medial showing of the CSF

#### 26. L Inf. AM Temp. Horn

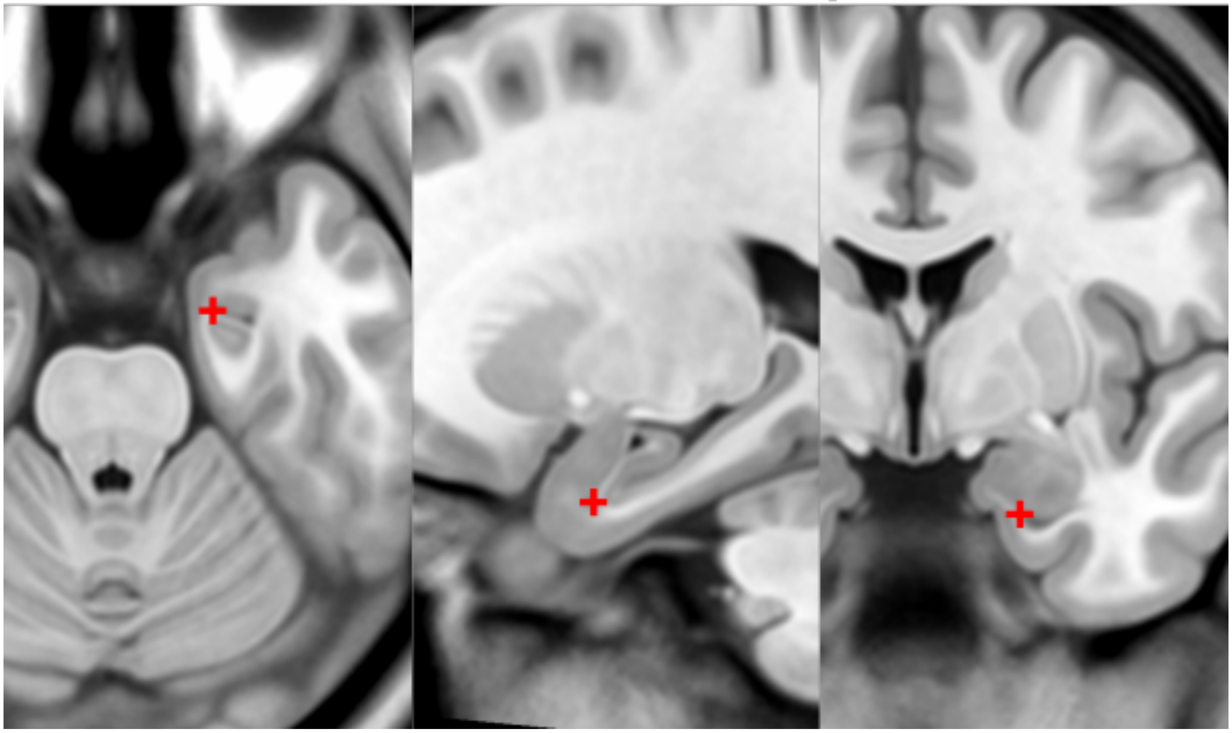

#### 27. R indusium griseum origin

- Defined on sagittal slice at takeoff from posterior hippocampus below splenium
- Begin on the sagittal view (make sure the view is on the right side), scroll back and forrth to find the point where the tail of the hippocampus begins to become pointed and "takeoff"

27. Right IG Origin

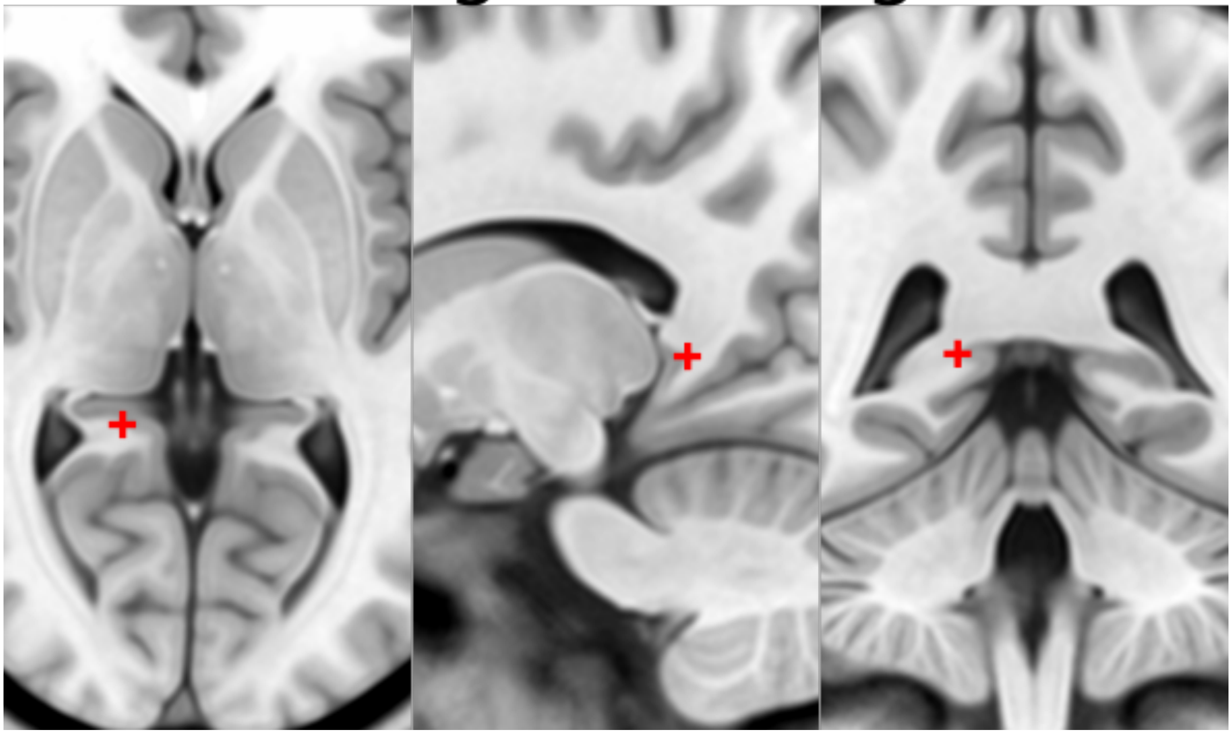

#### 28. L indusium griseum origin

- As in 27

28. Left IG Origin

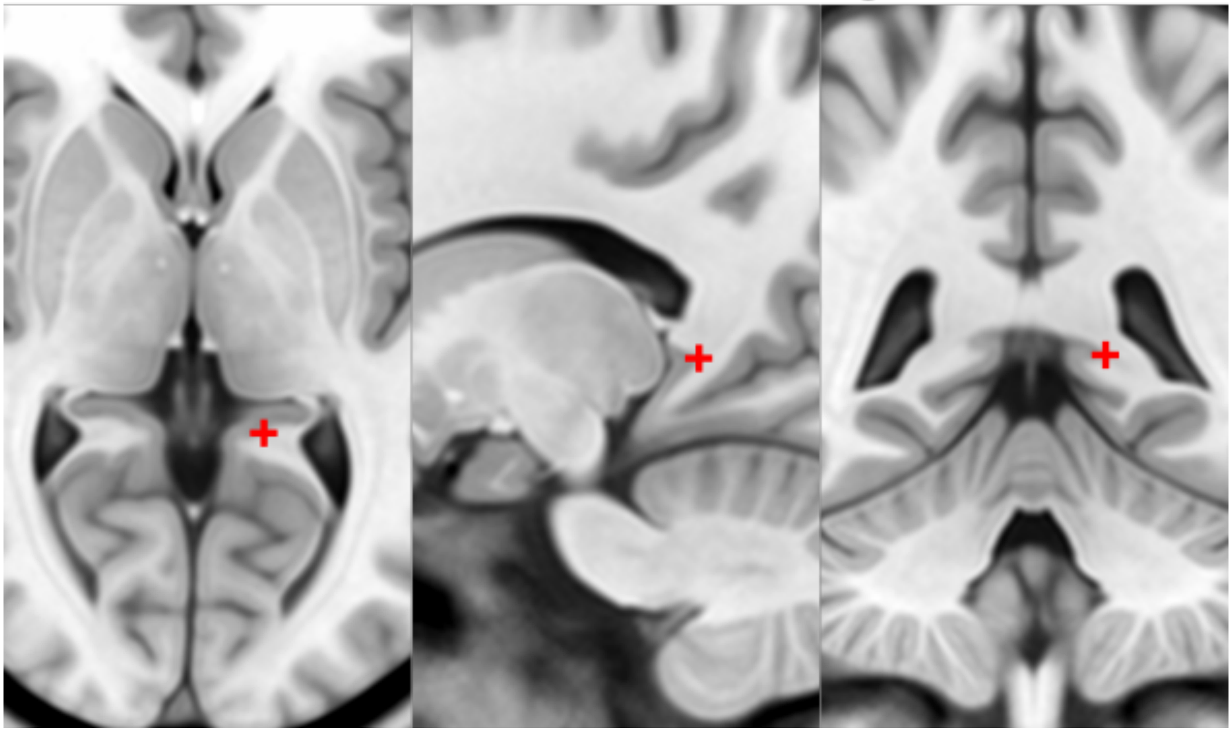

#### 29. R ventral occipital horn

- Defined on ventral/inferior portion of last visible coronal slice with occipital horn
- If it is hard to see on the coronal view then you can make the first placement using the axial view (make sure the view is on the right side of the brain).
- Optimize using other views

29. R Ventral Occipital Horn

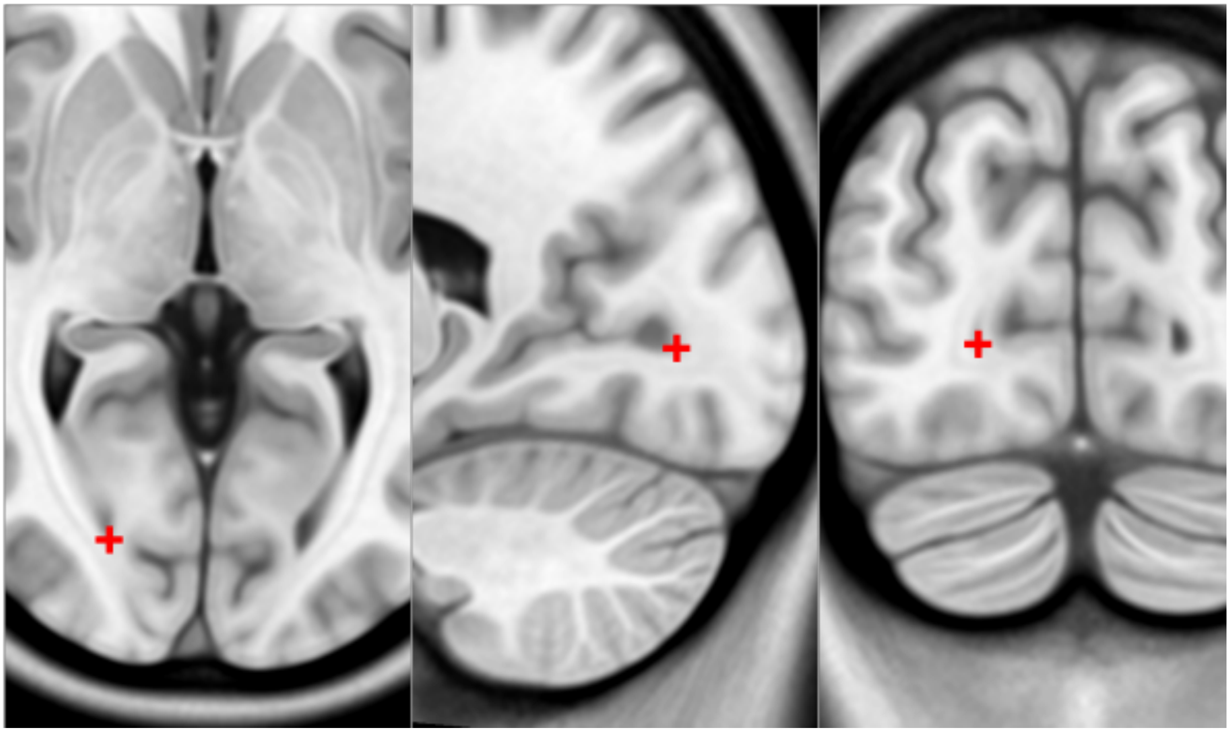

#### 30. L ventral occipital horn

- As in 29

### 30. L Ventral Occipital Horn

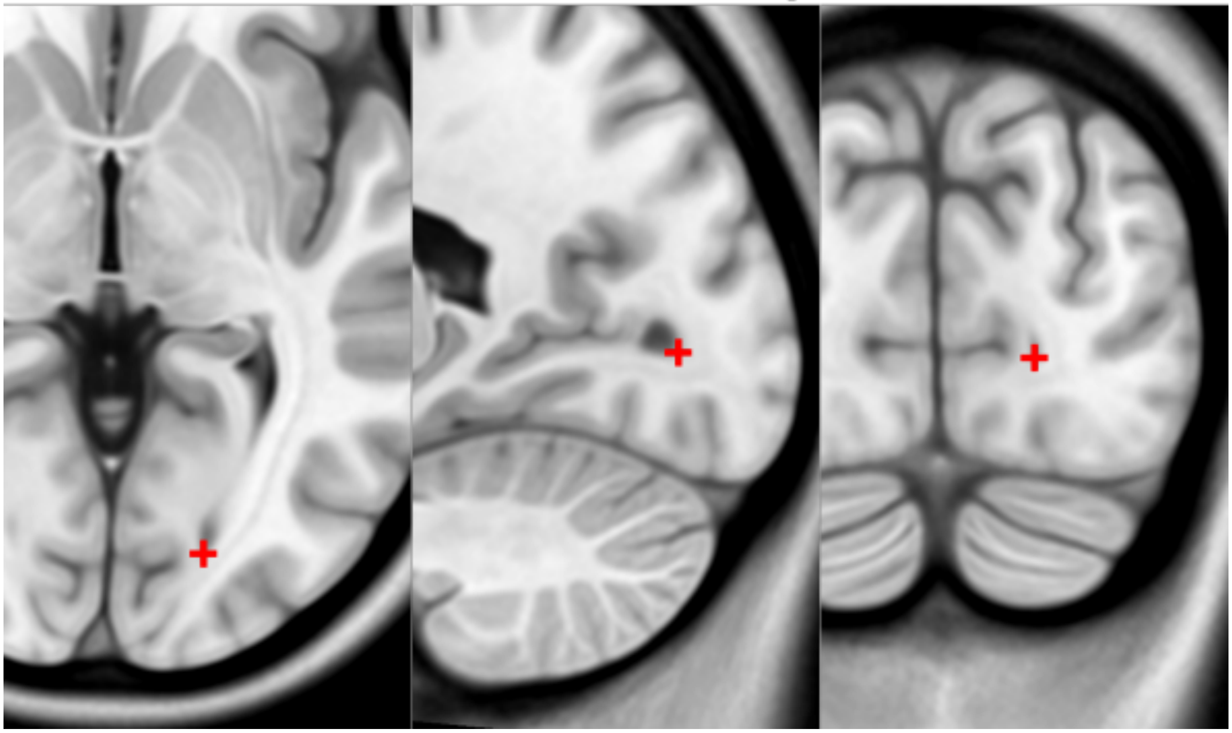

#### 31. R olfactory sulcal fundus

- Sulcal fundus = at depth of sulcus and boundary of gray matter-white matter
- Posterior and most superior portion visible on axial slice

31. R Olfactory Fundus

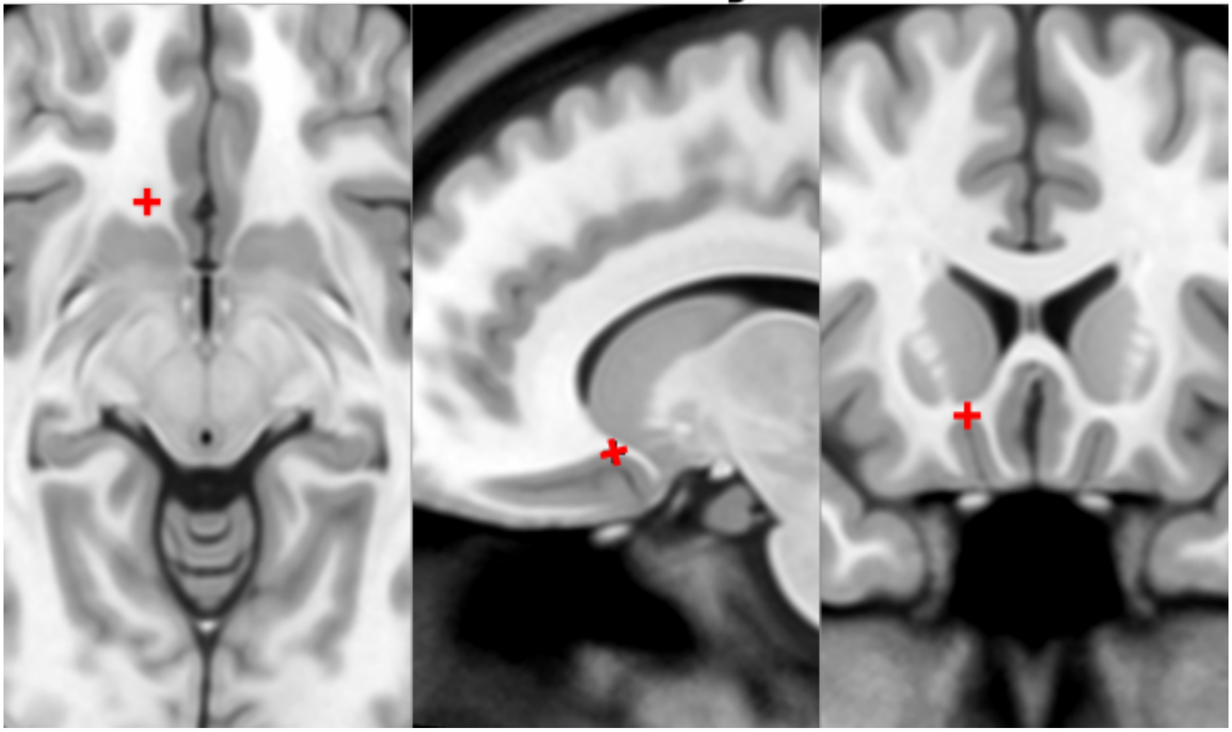

#### 32. L olfactory sulcal fundus

- As in 31

32. L Olfactory Fundus

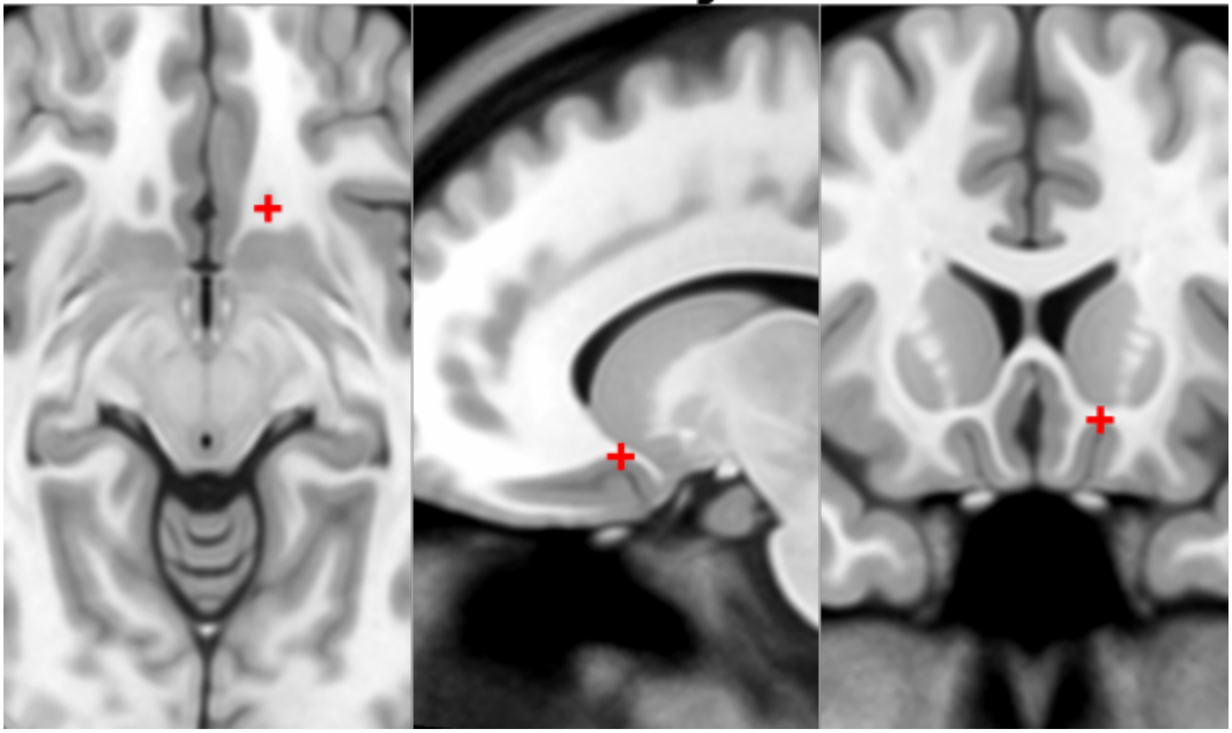

Save the fiducial file (.fcsv) locally.

#### Resources:

- Tutorial given as part of Brainhack Western 2018: https://www.youtube.com/watch?v=huGtd19_uiM
